# Recent rapid radiation of apex predators suggests dramatic biodiversity turnover in an ancient lake

**DOI:** 10.1101/2025.01.14.633002

**Authors:** Jessica A. Rick, Julian Junker, Alexander L. Lewanski, Brittany Swope, Michael M. McGlue, Emmanuel A. Sweke, Ismael A. Kimirei, Ole Seehausen, Catherine E. Wagner

## Abstract

Top predators have oversized influence on food webs and ecosystem dynamics, and the arrival of a predator in a naive environment can have dramatic consequences for endemic biodiversity. Using a combination of whole genome and reduced-representation genomic data, and pairing these with phylogenetic and demographic modeling approaches, we find that the colonization of East Africa’s Lake Tanganyika by *Lates* fishes—the top predators in this ancient lake—occurred *∼*1–2 Mya, which is more recent than other major and diverse clades within the lake. Diversification from one into four endemic *Lates* species occurred within the lake during a time of dramatic changes in lake levels driven by glacial–interglacial cycles, supporting the the role that environmental conditions likely played in speciation between these taxa. The lake level fluctuations also occurred during a time with multiple inferred admixture events among *Lates* species during the mid- to late Pleistocene (*∼*90–500 Kya). Together, our findings suggest a dynamic and environmentally-linked evolutionary history of this predator radiation, and that their colonization of the lake and subsequent diversification likely had dramatic ecosystem consequences for taxa already present in Lake Tanganyika. This provides novel insight into the role that climatic cycles play in evolution in freshwater lakes, as well as insight into the evolutionary and ecological history of this extremely biodiverse ecosystem.

**Teaser Text:** The arrival of top predators in new ecosystems can substantially alter biodiversity and food webs. These new habitats can also provide invading taxa with ecological opportunities that facilitate species diversification. In this research, Rick *et al*. find evidence that the radiation of endemic top predators in East Africa’s Lake Tanganyika originated surprisingly recently, and that these species have experienced repeated periods of hybridization with a congeneric, widespread riverine species throughout their history. These findings have major implications for the history of the lake and suggest that the anthropogenic introduction of Nile perch into Lake Victoria, which caused dramatic ecosystem and food web changes, may be a contemporary analog to the natural historical events in Lake Tanganyika.

## Introduction

Predators have strong direct and indirect effects on biodiversity, as well as a strong influence on ecosystem dynamics (Carpenter et al., 1985; Estes et al., 2011). The appearance of top predators in naive ecosystems can cause major alterations to species diversity, trophic interactions, and community structure (Alston et al., 2019; Schmitz et al., 2004; Wainright et al., 2021), both within and across ecosystems (Koel et al., 2019), and can induce evolutionary change in native species (Mooney and Cleland, 2001). While many recent predator colonization events have been human-mediated, predators can also reach new habitats without human assistance, often as a result of novel habitat formation or reorganization caused by changing environmental conditions.

The same environmental changes that facilitate predator colonization of new habitats can also drive broader patterns of speciation and hybridization through facilitating cycles of isolation and contact among populations (He et al., 2019; Linck and Battey, 2019; Nevado et al., 2018). Pleistocene glacial–interglacial cycles transformed high latitude landscapes, caused wide-ranging global climatic changes (DeMoncal, 1995; Lyons et al., 2015; Tierney et al., 2008b), and have been invoked to explain speciation patterns in mammals (e.g., Aubry et al., 2009; Barnosky et al., 2004), birds (e.g., Claramunt and Cracraft, 2015; Weir et al., 2016; Weir and Schluter, 2004), arachnids (e.g., Ayoub and Riechert, 2004), plants (e.g., DeChaine et al., 2014; Nevado et al., 2018), marine invertebrates (e.g., Clarke and Crame, 2010), and freshwater fishes (e.g., April et al., 2013). Global glacial periods are also associated with megadrought conditions in Africa (Fig. S1; Russell et al., 2020; Tierney et al., 2008a; Verschuren et al., 2009), and these periods had dramatic consequences for African freshwater ecosystems, many of which contracted or disappeared during drought periods (Burnett et al., 2011; Cohen et al., 2007; Lyons et al., 2015; Scholz et al., 2007). Tropical East Africa—and in particular, the East African Rift Lake system—is particularly sensitive to precessional-cycle climate shifts (i.e., those due to the rotational axis of the planet; Clement et al., 2004) due to its location within the continent. Changes to these cycles have triggered rapid climate change in the region, resulting in a shift from wet to dry conditions periodically during the Plio-Pleistocene (Trauth et al., 2010, 2007). As a consequence, both climatic and tectonic events since the early Miocene have periodically altered environmental suitability and barriers to dispersal for organisms inhabiting the region, which in turn shaped patterns of biodiversity in both aquatic and terrestrial taxa (Salzburger et al., 2014; Stewart, 2001), including early hominids (Levin, 2015; Trauth et al., 2010).

Ancient lakes are among the best records of past environmental change; these lakes not only record long histories of environmental variation and human activity in their sediments, but also harbor high levels of biodiversity and endemism. The African Great Lakes region is home to numerous species radiations and abundant biodiversity (Salzburger et al., 2014). Lake Tanganyika (LT; Fig. 1)—the largest (by volume; 18,880km^3^), deepest (1,470m), and oldest (*∼*9–12 My; Cohen et al., 1993) of the African Great Lakes—represents one of the most diverse aquatic ecosystems in the world (Groombridge and Jenkins, 1998) and is home to >1500 aquatic animal species, of which >600 are endemic (Coulter, 1991a; Salzburger et al., 2014) and >200 belong to a radiation of endemic cichlid fishes. Evolutionary diversification in these speciose clades is often linked to divergences in niche space as ecological opportunity arises (Wagner et al., 2012), and the fish radiations have diversified along many common ecological axes (Salzburger et al., 2014; Seehausen and Wagner, 2014).

**Fig. 1.**
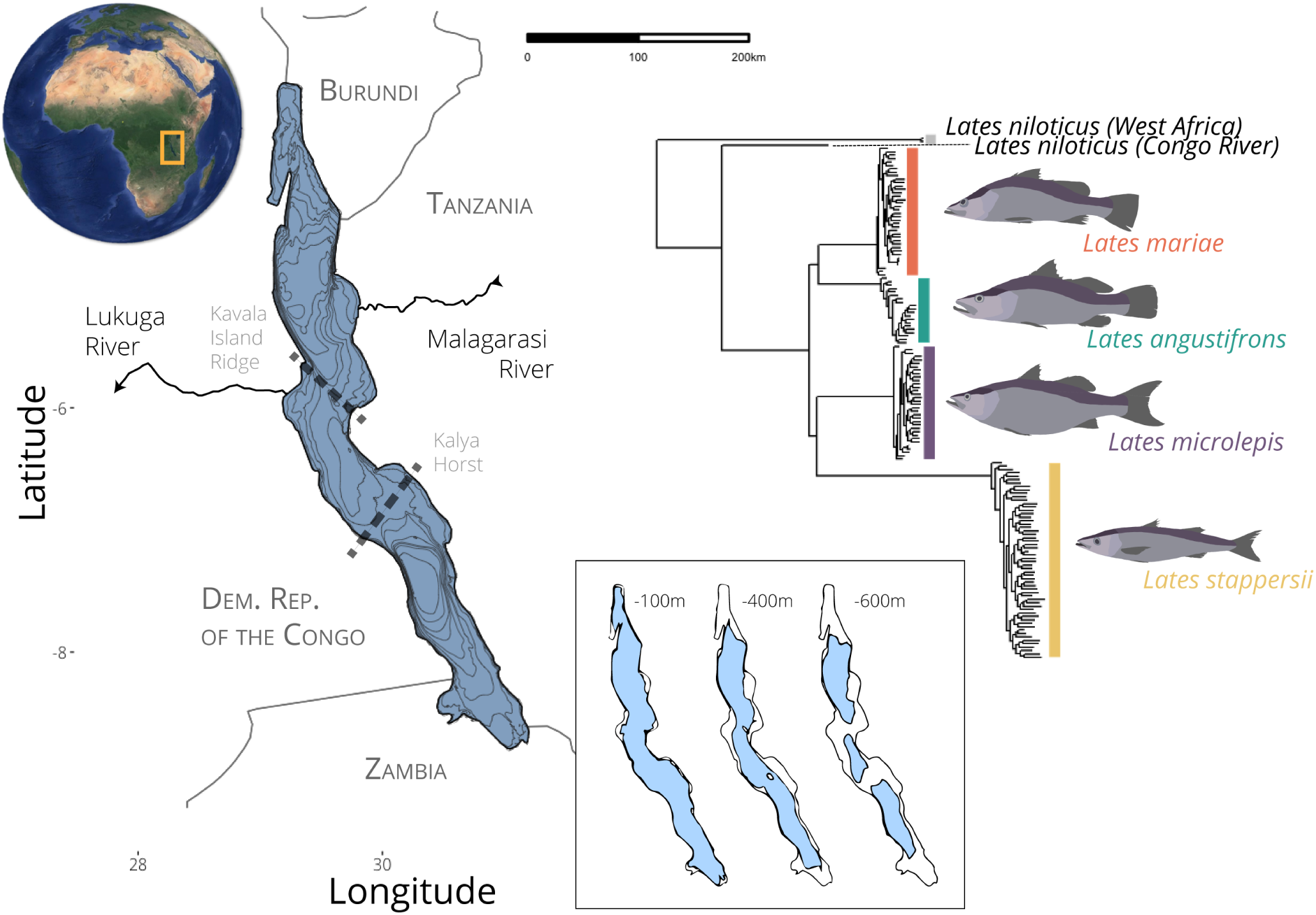
Map of Lake Tanganyika, with inflow via the Malagarasi River and outflow via the Lukuga River indicated, as well as the location of the Kavala Island Ridge and Kalya Horst, separating the north and middle, and middle and south basins, respectively. The inset map shows the inferred lake extent at 100m, 400m, and 600m below the present surface of the lake based on present-day bathymetry, all levels that are likely to have occurred as a result of climatic cycling during the late Pleistocene (McGlue et al., 2008). The phylogeny indicates relationships among species inferred using concatenated genotyping-by-sequencing data from Rick et al. (2021), with the tree rooted using *Lates calcarifer* as an outgroup.

Recent studies based on large scale whole genome sequencing have led to important insights into diversification and hybridization dynamics of fish fauna within the African Great Lakes. Whole-genome sequencing of Lake Malawi’s cichlid radiation revealed that hybridization among ancient lineages fueled its extraordinary diversification, with admixture events providing genetic variation that was subsequently acted upon by selection to produce novel adaptive phenotypes (Malinsky et al., 2018; Svardal et al., 2020). Similarly, genome-wide analyses of the Lake Victoria Region Superflock—the large group of haplochromine cichlids that occupy Lake Victoria and nearby lakes—demonstrated that it originated from a hybrid swarm formed by the merging of divergent Congo and Nile drainage cichlid lineages (Meier et al., 2017, 2023), with this ancestral admixture contributing functional genetic variation that facilitated rapid adaptation to novel ecological niches (McGee et al., 2020; Meier et al., 2023). In Lake Tanganyika, whole-genome data have revealed complex histories of introgression among cichlid species, with signatures of ancient and recent admixture pervading the radiation (Irisarri et al., 2018; Ronco et al., 2021). In other taxa, the signal of admixture is usually distributed heterogeneously across the genome, with regions of low recombination tending to be more resistant to introgression, while regions of high recombination are more permeable to gene flow, a signal that is often interpreted as selection against admixed ancestry (Martin et al., 2019; Moran et al., 2021; Schumer et al., 2018). Alternatively, positive selection on introgressed regions can also produce a negative correlation between recombination rate and admixed ancestry. This interaction between recombination rate, selection, and introgression can shape genome-wide patterns of divergence and generate heterogeneous landscapes of differentiation, complicating the inference of demographic history from genomic data (Edelman et al., 2019; Ravinet et al., 2017). However, these patterns have yet to be investigated in-depth in African Great Lakes cichlid radiations, despite there being much evidence for extensive admixture among taxa (Svardal et al., 2021). While these genome-wide insights have transformed our understanding of cichlid diversification, comparable whole-genome analyses of non-cichlid radiations in the African Great Lakes remain limited, leaving open the question of whether similar processes have shaped diversification in the other clades inhabiting these ecosystems.

In Lake Tanganyika, the open-water pelagic zone is highly depauperate in species compared to the species-rich littoral zone (Coulter, 1991b). It is dominated by two species of endemic sardines (*Limnothrissa miodon*, *Stolothrissa tanganicae*) and a four-species radiation of piscivores in the genus *Lates*. The LT *Lates* clade is the most species-rich lacustrine *Lates* radiation globally, and it is a highly unusual adaptive radiation of freshwater fish in that all four species are piscivorous predators as adults, with differentiation in habitat, life history, and sizeand only slight differences in trophic position (Coulter, 1991a; Ehrenfels et al., 2023). These four species occupy different ecological niches, with the small-bodied *Lates stappersii* (size at maturity 24cm, max length 45cm) occupying pelagic areas throughout its life (Ellis, 1978), while the three larger-bodied species inhabit inshore areas as juveniles and move into deeper and more open waters as adults (Coulter, 1991b). As adults, *L. microlepis* (size at maturity 50cm, max length 100cm) also lives near the surface in the pelagic zones, while *L. mariae* (size at maturity 46cm, max length 80cm) predominantly occupies deepwater benthic areas and *L. angustifrons* (size at maturity 57cm, max length 200cm) primarily occupies the demersal zone, with periodic movements into pelagic areas (Coulter, 1976, 1991b). Juveniles of *L. mariae* and *L. angustifrons* generally feed on shrimps as juveniles and transition to piscivory as they grow (Kondo and Abe, 1995), while *L. stappersii* feeds on shrimp and Tanganyikan sardine species (*Limnothrissa miodon* and *Stolothrissa tanganicae*) throughout its life stages (Mannini et al., 1999). Freshwater *Lates* are native and widespread in African rivers and have a patchy distribution among large lakes in East Africa; however, all of the species except *L. niloticus* are endemic to single lakes. The widespread *L. niloticus*, in contrast to these single-lake endemics, is native to both river and lake environments across central, western, and eastern Africa—and has been introduced into several lakes in East Africa, including Lake Victoria (Basiita et al., 2018; Mwanja et al., 2013, 2011)—with distinct mitochondrial haplotypes found in western versus eastern African locales (Basiita et al., 2018; Mwanja et al., 2013) and three distinct geographic lineages genome-wide (Koblmüller et al., 2021). These predators have large influences on their ecosystems, most clearly demonstrated by the introduction of *L. niloticus* into Lake Victoria, which famously led to ecological reorganization of the fish community and extinctions in the endemic pelagic cichlid community (Goudswaard et al., 2008; McGee et al., 2015; Seehausen, 1999).

Here, we use genome-wide analyses to investigate the evolutionary history of LT’s *Lates* species in relation to environmental change across the lake’s history. We find genomic evidence to suggest that global climatic cycles may have shaped the tempo and extent of this radiation via periodic bouts of admixture between *L. niloticus* and LT’s endemic *Lates* species, as well as recurrent opportunities for allopatric divergence. We further examine how recombination rate variation relates to the genomic distribution of introgression, and consider the potential role of linked selection in shaping patterns of admixture across the genome. Furthermore, we find evidence that *Lates* only recently arrived in Lake Tanganyika, which has implications for the rapidity of this radiation’s diversification and suggests that their arrival may have had significant, ecosystem-wide impacts on the lake’s already biodiverse fauna.

## Methods

### Study species and sampling

Our analyses focus on the widespread *L. niloticus* and the four *Lates* species endemic to Lake Tanganyika: *L. angustifrons, L. mariae, L. microlepis*, and *L. stappersii*. The *L. niloticus* specimens were obtained from tissues archived in the Cornell University Museum of Vertebrates (CUMV) and American Museum of Natural History (AMNH). A total of 152 tissue samples were sequenced using a genotyping-by-sequencing (GBS) protocol, as described in Rick et al. (2021), and 13 of those were additionally used to generate whole genome resequencing (WGS) data. The majority of the samples used for WGS came from sites near Kigoma, Tanzania, or Mahale Mountains National Park, Tanzania (Table S1). Samples for GBS sequencing were spread across the eastern shore of the lake (details in Rick et al., 2022), and these samples previously have shown some structure in the far southern portions of the lake in *L. mariae* and no obvious geographic population structure in *L. angustifrons*, *L. microlepis*, and *L. stappersii*. Reads for both data types were aligned to the chromosome-level *Lates calcarifer* genome assembly (Vij et al., 2016), as it is the most closely related high-quality reference genome available. WGS haplotypes were phased using WhatsHap (Martin et al., 2016) and population recombination rates in each species across each chromosome were estimated using the the recombinant neural network method implemented in ReLERNN (Adrion et al., 2020). More detailed sampling and bioinformatic methods can be found in the Supplemental Methods.

### Phylogenetic and diversification histories

We inferred phylogenetic relationships among taxa using both maximum-likelihood and gene tree methods, implemented in RAxML (Stamatakis, 2014) and ASTRAL-III (Zhang et al., 2018), from both GBS and WGS data. To investigate whether genomic regions exhibiting elevated levels of introgression contribute disproportionately to topological uncertainty, we conducted sliding window analysis of gene trees in TWISST2 (Martin, 2026; Martin and Van Belleghem, 2017). We estimated dates for divergence between taxa based on the relative cross-coalescence rates from the WGS data using MSMC2, using a generation time of 3 years and mutation rate of 3.5*x*10*^−^*^9^ per base pair (Malinsky et al., 2018) to convert coalescent units to years.

### Admixture inferences

We used Dsuite (Malinsky et al., 2021) to calculate genome-wide *D*-statistics using both WGS and GBS data for all three-taxon combinations of *L. niloticus, L. angustifrons, L. mariae, L. stappersii,* and *L. microlepis*, again always keeping *L. calcarifer* as the outgroup. To determine the extent of introgression and the likely location of introgression events on the *Lates* phylogeny, we used Dsuite to calculate *f* 4-ratio statistics (Patterson et al., 2012) and, from these, calculate *f* -branch statistics (Malinsky et al., 2018) constrained to the maximum likelihood topology inferred in RAxML. To localize signals of admixture across the genome, we conducted sliding window analyses of *f_d_*for those comparisons with a genome-wide significant *D* statistic. We calculated the *z*-score for each *f_d_* value, as well as its associated *p*-value, and identified significantly admixed regions as those with *z* > 3. We calculated length of introgression tracts for each comparison on each chromosome, and investigated relationships between introgression tract length and time since introgression, as well as between the proportion of a chromosome that was contained in significant introgression tracts and the length of that chromosome (Supplemental Methods).

### Demographic modeling

To test competing hypotheses for the evolutionary history and gene flow among our focal species, we inferred demographic history based on the site frequency spectra (SFS) summary of WGS sequences using the continuous-time sequential Markovian coalescent method implemented in fastsimcoal2 (Excoffier et al., 2013). We built 32 six-taxon models including the four Lake Tanganyika species, *L. niloticus*, and *L. calcarifer* (Fig. S2). The models all had the same divergence history matching the topology in the best RAxML species tree but differed in the number of admixture events between taxa. In all models, we constrained the split between *L. calcarifer* and the freshwater *Lates* to 11 million generations based on the oldest known *Lates* fossil found in Africa (*L. qatraniensis, ∼*33 Mya; Murray and Attia, 2004). We assessed model fit and performance via both likelihood distribution and Akaike information criterion (AIC) methods, following Meier et al. (2017). We used a non-parametric block bootstrapping approach (Meier et al., 2017) to generate 100 bootstrap replicates of parameter estimates for our best model. We then constructed two-taxon fastsimcoal2 models for all pairs of ingroup taxa (i.e., not including *L. calcarifer*) using the GBS data. For each pair, models with 0–3 admixture events (10 total models) were competed to infer the most likely number of admixture events between the given pair of taxa.

## Results

We generated genotyping-by-sequencing (GBS) and whole genome resequencing (WGS) data for *Lates* specimens from the four LT-endemic species and three *L. niloticus* museum specimens. For the focal LT *Lates* species, the average nucleotide diversity (*π*) was 0.040 (range of species estimates 0.032–0.056, with highest diversity observed in *L. stappersii* (Fig. S3). The between-species nucleotide divergence (*d_xy_*) among the LT species ranged from 0.0578 to 0.092, and divergence between the LT species and the widespread *L. niloticus* had a mean *d_xy_* = 0.119 (range 0.110–0.140). With both GBS and WGS, maximum likelihood concatenation (Stamatakis, 2014) and gene tree (Zhang et al., 2018) methods inferred a species tree with a monophyletic LT group and two sister species pairs within Lake Tanganyika (Fig. 1, 2a). Monophyly of the LT species is consistent with the phylogenetic hypothesis proposed by (Koblmüller et al., 2021), but intra-LT relationships differ. However, in both our results and those of Koblmüller et al. (2021), there are short internode distances during LT diversification. Our analyses show high gene tree incongruence (Fig. S4). With GBS, where we had sampling from multiple locations for *L. niloticus*, we additionally infer that *L. niloticus* is paraphyletic, with *L. niloticus* from the Congo basin sister to the LT clade, but West African individuals forming a distinct clade sister to the clade of Congo basin *L. niloticus* plus LT *Lates* (Fig. 1), mirroring the recent findings of (Koblmüller et al., 2021) and agreeing with our own analyses of mitochondrial haplotypes (Fig. S7).

Topology weightings for 100-SNP windows across the genome showed that the most common topology for all 24 of the *L. calcarifer* chromosomes matched the species tree (Fig. S6). However, only 2.63% of the genome supported this relationship (Fig. S5), while 9.4% supported the four topologies classified as “alternate” topologies (i.e., those in which one sister relationship agreed with the species tree topology and the LT taxa were monophyletic), and 72.6% supported the 90 possible topologies with *L. niloticus* nested within the LT clade.

### Evidence for ancient and recent admixture with *L. niloticus*

Short internodes in the *Lates* species tree suggest the possibility that these fish may have radiated quickly enough to be in the phylogenetic “anomaly zone”, where the majority of gene trees may reflect incomplete lineage sorting (ILS) rather than the true species tree. However, gene tree incongruence can also follow from admixture events within the clade’s history. We calculated *D*-statistics (Korneliussen et al., 2014; Malinsky et al., 2021) among sets of three taxa to distinguish among these possibilities. All comparisons with *L. niloticus* nested within the LT radiation had higher *D* than those with *L. niloticus* sister to two LT taxa, suggesting that all LT species are more closely related to one another than any one is to *L. niloticus*, providing further support for a monophyletic LT clade (Table S2). The remaining significant *D* estimates indicate excess allele sharing between *L. niloticus* and each of *L. angustifrons*, *L. microlepis*, and *L. stappersii*, although with weaker signal in *L. stappersii* (Table S3). The derivative Fbranch analysis (Malinsky et al., 2018, Fig. 2b) suggests that this excess allele sharing comes from an admixture event between *L. niloticus* and the common ancestor of *L. mariae* and *L. angustifrons*, with additional independent *L. niloticus*–*L. angustifrons* and *L.niloticus*–*L. microlepis* admixture events following intra-LT divergence into four species. Additionally, there is evidence for admixture between *L. microlepis* and each of *L. mariae* and *L. angustifrons*, and between *L. stappersii* and *L. mariae*. Using GBS data containing samples from West African *L. niloticus* populations, we split West Africa and Congo populations and find evidence that admixture is stronger between the Congo River lineage and LT species and weaker in the West African lineage (mean *D* = 0.213, *F*_4_-ratio = 0.139; Fig. S8a). Using an alternate phylogenetic hypothesis (from Koblmüller et al., 2021) in our Fbranch analysis suggests admixture between West African *L. niloticus* and the LT common ancestor that does not involve the Congo River lineage (Fig. S8b,c); while this would be difficult to imagine given current hydrological connections, the region has undergone much hydrological reorganization in the time since Lake Tanganyika formed (e.g., Beadle, 1974). In addition, it is possible that an unsampled or extinct lineage contributed to this observed pattern in our data (e.g., Tricou et al., 2022).

**Fig. 2.**
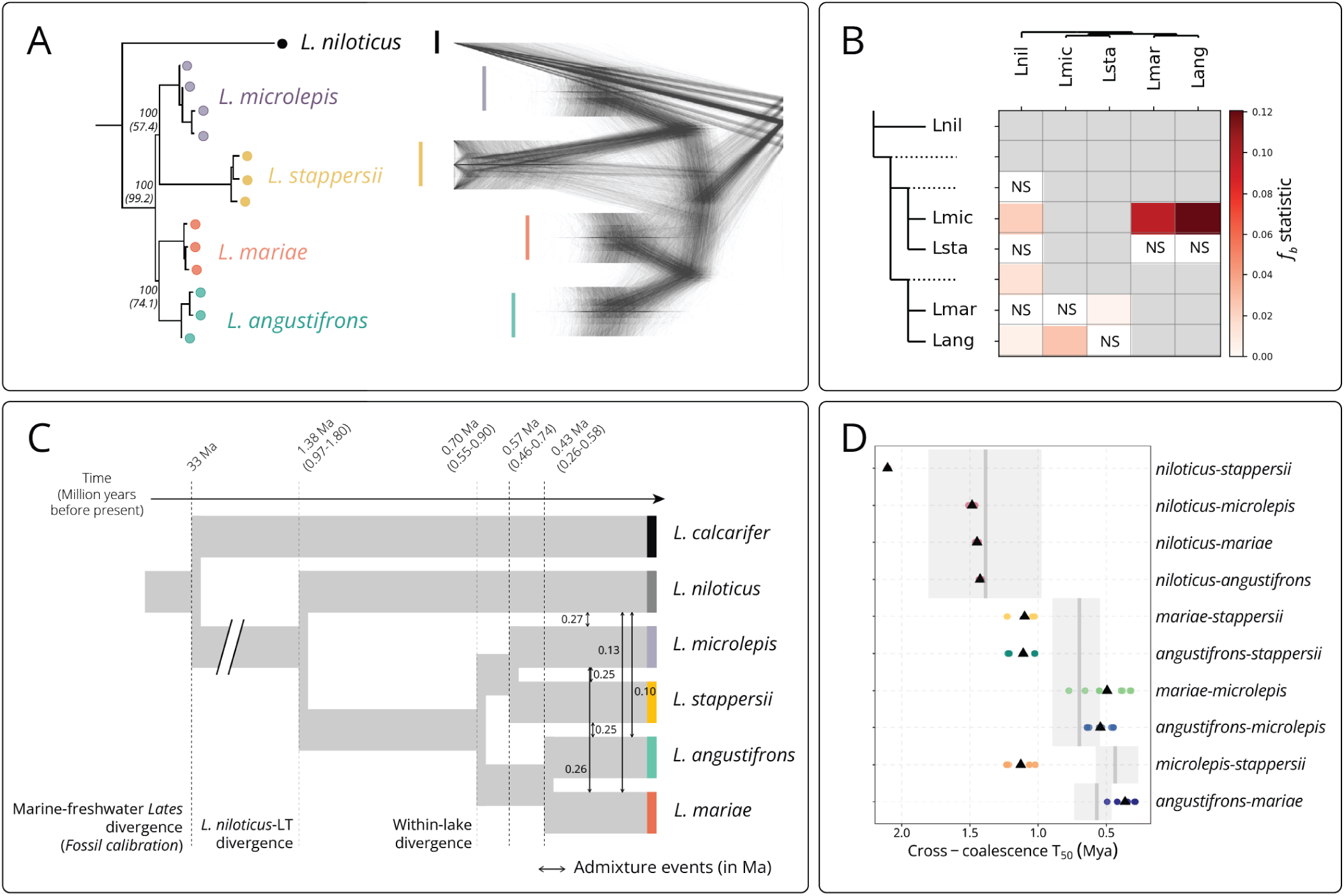
Multiple lines of evidence support a monophyletic Lake Tanganyika *Lates* clade with a relatively recent origin *∼*1–2 Mya and a history of admixture among species. (A) Species tree inference from whole genome sequences (left), rooted using *L. calcarifer* as the outgroup and including the *L. niloticus* sequence from the Congo River, demonstrates monophyly of the Lake Tanganyika clade (*L. microlepis, L. stappersii, L. mariae, L. angustifrons*). Confidence from bootstrap estimates (from RAxML) are shown for major splits, with quartet scores (from ASTRAL) shown in parentheses. Gene trees from 4000bp windows (right) show incongruence contributing to uncertainties in species tree inference. (B) The Fbranch statistics, conditioned on the species tree in (A), infer significant admixture events between many of the focal taxa. Darker red colors indicate a stronger signal of gene flow. Gray boxes indicate comparisons that could not be tested due to being sister taxa, while white boxes indicate non-significant comparisons. (C) Visualization of the best model from the 32 tested fastsimcoal2 demographic models, with the median bootstrap estimates for timing of divergence (dashed lines) and admixture (arrows) events labeled. Generation times have been converted to years using an average generation time of three years. For divergence events, we show estimates from the best model and 95% bootstrap intervals from nonparametric bootstrapping. (D) Reconstruction of divergence times using relative cross-coalescence rates for each pair of individuals in MSMC2 provide an estimate of divergence times as a comparison to estimates in (C). Divergence times are inferred as the point when the cross-coalescence rate reaches 50%, *T*_50_. Colored dots show the estimated *T*_50_ for each pair of individuals, while black triangles indicate the mean for each species pair. To provide a direct comparison to fastsimcoal2 results, estimates (dark gray lines) and bootstrap confidence intervals (light gray boxes) are shown from fastsimcoal2 inferences.

For each *L. niloticus*–LT *Lates* pair showing evidence for introgression from our Fbranch analysis, we identified regions of the genome involved in admixture by calculating the *f_d_* statistic (Martin et al., 2015) in 100-SNP sliding windows. The proportion of significant *f_d_* regions was similar across chromosomes (mean *f_d_*=0.51%, range=0.014–2.36%), although one chromosome (LG14) had an elevated proportion of admixed windows compared to the other chromosomes (Table S4; Fig. S9). Several *f_d_* peaks were shared across three-taxon comparisons (Fig. S10), while others were unique to specific species. The signal of admixture was elevated in regions with low recombination (Fig. 3b; Pearson’s *r*(10499) = *−*0.0818*, p <* 2.2*x*10*^−^*^16^), and both *f_d_* and recombination rate increased near chromosome ends (Fig. 3c,d). In addition, an elevated *f_d_*signal was associated with low topology weighting for the topology matching the RAxML tree (Fig. 3a; Pearson’s *r*(10499) = *−*0.0970*, p <* 2.2*x*10*^−^*^16^). Admixture proportions calculated using *f_d_*indicate that a higher proportion of the genome is involved in admixture between *L. angustifrons* and *L. niloticus* than between the other species and *L. niloticus* (Fig. 4b). These proportions varied among chromosomes, suggesting heterogeneity in the fate of introgressed regions across the genome, with shorter chromosomes generally having a higher proportion of their length showing elevated signal of admixture (Fig. 4d).

**Fig. 3.**
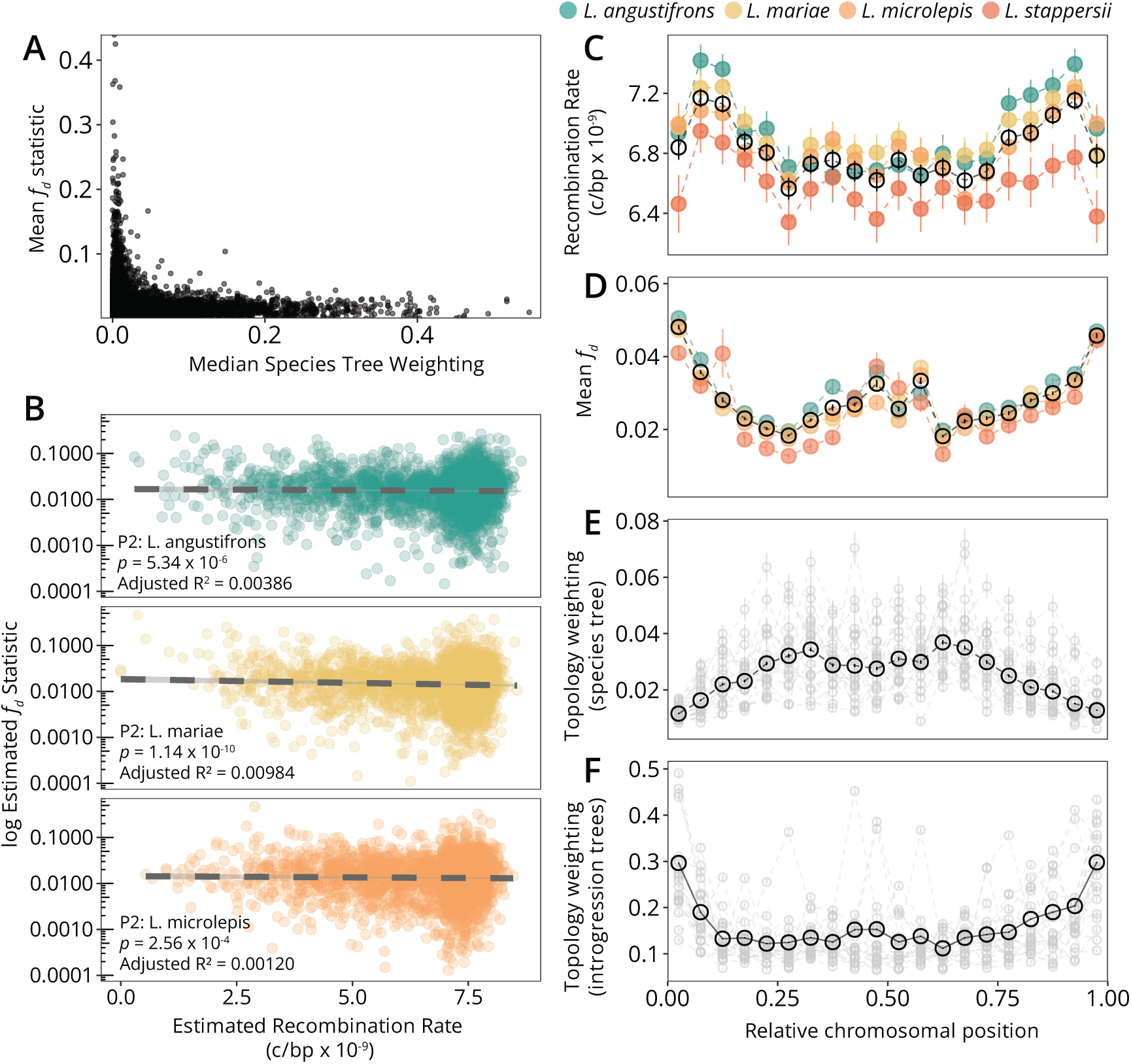
The signal of admixture varies across the *Lates* genome, with admixed regions contributing to gene tree incongruence, and regions with lower recombination rates holding higher admixture proportions. (A) Windows with higher *f_d_* values were most often those with a low median topology weighting for the species tree. (B) Estimated recombination rates are correlated with both *f_d_*, where regions with higher recombination rates tend to have lower *f_d_* estimates. Each point represents data binned to 5000bp regions for both statistics. The y-axis has been log-transformed for better visualization of the data. In recombination rates (C), *f_d_*estimates (D), and topology weighting estimates for the introgression trees (F), mean estimated values generally increase toward the ends of chromosomes, with a smaller peak near the middle of chromosomes, which is most pronounced in *f_d_*estimates. In examining the topology weighting support for the species tree (E), the complementary pattern is visible. For recombination rates and *f_d_*estimates, colored points show binned estimates for each species (grouped by P2 species for *f_d_*), while black open circles show binned means across all data. In topology weighting plots, gray circles show binned means for each chromosome, while black open circles show binned means across all data.

**Fig. 4.**
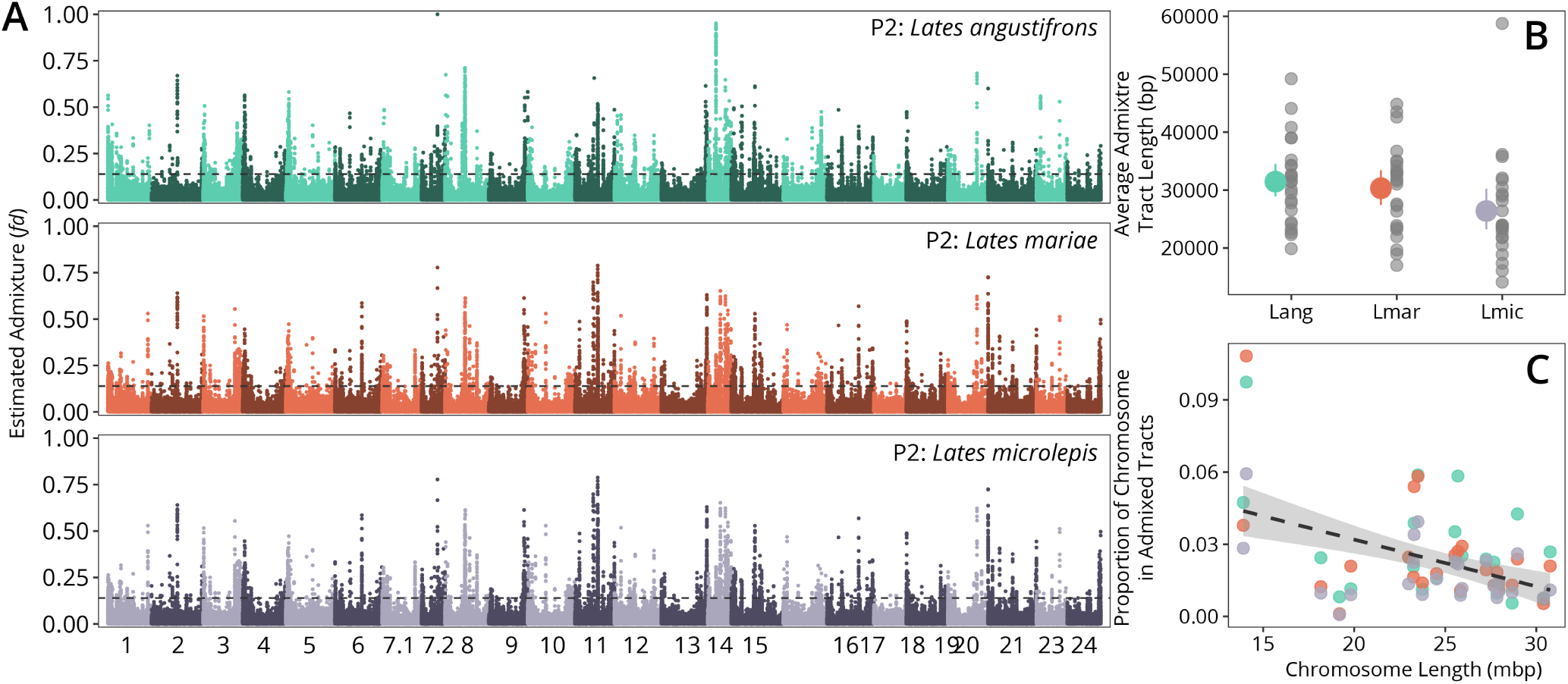
(A) Sliding window analysis of *f_d_*across the genome (P1: Other LT taxon, P2: the indicated species, P3: *Lates niloticus*) demonstrates variation in admixture signal across chromosomes, with some shared and some independent admixed regions among taxa. The horizontal dashed line indicates the *z*-score = 3 for *f_d_* estimates genome-wide, and each point is the maximum value estimated across comparisons for the given P2 taxon. (B) The average length of admixed windows varied among taxa, with the longest admixed regions found in *L. angustifrons* (Lang) and shorter average lengths in *L. mariae* (Lmar) and *L. microlepis* (Lmic). For each, gray points show the variation among chromosomes in mean admixture tract size. (C) A negative relationship was observed between total proportion of a chromosome that was contained in admixture tracts and the length of the chromosome. Points are colored to match the colors in (B), representing the P2 taxon.

### Divergence and admixture timing and magnitude

Analysis of cross-coalescence among species in MSMC2 (Schiffels and Durbin, 2014) estimated an average divergence time (i.e., time to 50% cross-coalescence rate) of 1.61 Mya (range 1.41–2.10 Mya; Fig. 2c, Table S5) between *L. niloticus* and LT *Lates* species. Divergence between *L. angustifrons* and *L. mariae* dated to *∼*0.49 Mya, between *L. microlepis* and *L. stappersii* dated to *∼*1.21 Mya, *L. angustifrons–L. microlepis* to 0.53 Mya, *L. mariae–L. microlepis* to 0.55 Mya, *L. angustifrons–L. stappersii* to 1.21 Mya, and *L. mariae–L. stappersii* to 1.23 Mya (Fig. 2c, Table S5).

Site frequency spectrum-based coalescent modeling of demographic history in fastsimcoal2 indicated bouts of ancient admixture between *L. niloticus* and the LT *Lates* species, as was indicated with the *f_d_* analyses. Model selection using both AIC and distributions of likelihood values (Fig. S14) indicated that our best demographic model involved six independent bouts of admixture among our focal taxa (Fig. 2c). We estimated the split between *L. niloticus* and the LT *Lates* clade at 1.38 Mya (95% bootstrap CI: 0.97–1.80 Mya) and between the two LT sister clades at 697 Kya (95% bootstrap CI: 545–895 Kya). The two Tanganyika clades then split with divergence times dating to 570 (460–735) Kya and 435 (264-577) Kya (Table S6, Fig. 2c).

Our best demographic model had an overlapping likelihood distribution with the next 10 best models (Fig. S14), and therefore we examined the distribution of estimated divergence dates across these top models. The mean divergence time estimate for these 10 models between *L. niloticus* and the LT *Lates* clade was 1.3 Mya (95%: 1.2–1.7 Mya). The other intra-LT divergence times similarly overlapped with those of our best model, suggesting that our choice of best model does not significantly bias our divergence time estimates (Fig. S15). While the MSMC-inferred divergence times do not overlap with those from our demographic model for some of the within-lake speciation events (Fig. 2d), this is possibly because MSMC inferences do not account for post-divergence gene flow among taxa, and we find much evidence for gene flow from other analyses.

Based on bootstrap estimates from our best model (Fig. S16), admixture events between *L. niloticus* and the LT *Lates* occurred in two distinct bouts. Around 0.27 Mya (95% CI: 0.02–0.56 Mya), admixture occurred between *L. niloticus* and *L. microlepis*. In this same period, we also infer admixture events between *L. microlepis* and each of *L. stappersii* (0.25 Mya; 95% CI: 0.06–0.45 Mya) and *L. mariae* (0.26 Mya; 95% CI: 0.06–0.46 Mya), as well as admixture between *L. stappersii* and *L. angustifrons* (0.25 Mya; 95% CI: 0.05–0.46 Mya; Table S6). Closer to the present, we infer that admixture occurred between *L. niloticus* and *L. mariae* around 0.13 Mya (95% CI: 0.01–0.43 Mya) and between *L. niloticus* and *L. angustifrons* around 0.10 Mya (95% CI: 0.007–0.36 Mya). While confidence intervals for all of these admixture events overlap, they also are quite wide, suggesting much uncertainty in the exact timing of gene flow between taxa. In these admixture bouts, population size-scaled migration rates were higher from *L. niloticus* into the LT taxa than from LT taxa into *L. niloticus*, while intra-LT admixture events had more evenly balanced bidirectional migration rates (Table S6). Using two-taxon models with the GBS data (for which we had more individuals sequenced per species), we further investigated whether there is evidence for multiple bouts of admixture between taxa and find that the best models for *L. niloticus* and each of the LT taxa had either two or three admixture events (Fig. S17, S18), while those between LT taxa inferred either one bout of post-divergence admixture (for *L. angustifrons–L. microlepis*, *L. mariae–L. microlepis*, and *L. mariae–L. stappersii* or two bouts of admixture (*L. microlepis–L. stappersii*; Fig. S19, S20). The one exception was *L. angustifrons–L. stappersii*, whose best model supported three bouts of post-divergence admixture (Fig. S19, S20), although these admixture events were inferred to have occurred within a narrow window of time (188-200 Kya).

## Discussion

Using a combination of reduced representation and whole genome data and multiple approaches to model the evolutionary history of Lake Tanganyika’s endemic *Lates* species, we find that the origin of this radiation is much younger than the age of the LT basin, with colonization and diversification likely facilitated by climate-driven increases in precipitation that reconnected the lake to the Congo River system (Flügel et al., 2015) and allowed this novel predator to enter the LT ecosystem. The finding of a mid-Pleistocene origin of the radiation (*∼*1–2 Mya) has two major implications. First, it suggests that these top predator species may have speciated in the face of an already species-rich cichlid fauna, implying that their colonization of LT would likely have had dramatic ecosystem consequences. Second, it implies that all four species evolved during time periods when the lake is thought to have undergone cycles of water level change that subdivided distinct basins, which could have facilitated divergence and allopatric speciation; this timing supports the hypothesis that such cycles acted as a “species pump” (sensu Rossiter, 1995) within the lake, although more precise dating for both divergence times and lake level changes are needed to explicitly test this. Finally, we detect multiple, distinct bouts of admixture within the lacustrine *Lates* species and between the riverine *L. niloticus* and lacustrine *Lates*, further suggesting that cyclic patterns of sympatry and allopatry among these species likely influenced their evolution.

### Ecosystem implications of a young *Lates* clade

Two coalescent-based demographic modeling approaches both infer the origin of the Tanganyikan *Lates* clade to *∼*1–2 Mya and the crown age of the lacustrine radiation to *∼*0.70 Mya (Fig. 2). This age estimate is similar to but slightly younger than the result obtained via distinct methods and data from Koblmüller et al. (2021), which estimated a crown age of 0.87–2.35 Mya for the LT *Lates* clade and the divergence from *L. niloticus* at 2.01–4.59 Mya. These divergence estimates are much more recent than those for other endemic and speciose fish clades in the lake: current evidence suggests that other clades colonized the lake either near the onset of its formation 9–12 Mya (e.g., the cichlid species flock and *Mastacembelus* eels; Brown et al., 2010; Irisarri et al., 2018; Ronco et al., 2021) or during the establishment of a truly lacustrine habitat and deepwater conditions 5–6 Mya (Cohen et al., 1993) (e.g., the sardine species, *Synodontis* catfishes, and *Chrysichthys* catfishes; Day et al., 2009; Milec et al., 2022; Peart et al., 2014), and radiated soon thereafter. Even with the uncertainty that our analyses have in both mutation rates and generation times, ranges of reasonable estimates for each of these still place the *Lates* radiation younger than each of these clades. These inferred dates imply that the *Lates* clade diversified in the face of an already species-rich fish community in LT. Given that other large East African lakes without *Lates*, such as Lake Malawi (Fig. S21), have more diverse pelagic cichlid communities, it is possible that the arrival and diversification of *Lates* into LT would have had major ecosystem effects, namely through the extinction or competitive exclusion of a diverse pelagic cichlid community due to predation or competition from *Lates*.

With this insight, the recent anthropogenic introduction of *L. niloticus* into Lake Victoria acts as a modern analog for understanding the events that likely occurred within LT in the mid-Pleistocene following *Lates* colonization. In Lake Victoria, introduced *L. niloticus* outcompeted (McGee et al., 2015) and preyed on native fishes (Witte et al., 2007), which altered community and food web structure within the lake over the course of a few generations. The introduction of *L. niloticus* into Lake Victoria interacted with changing limnological conditions (Hecky et al., 2010; van Zwieten et al., 2015) to decimate cichlid stocks, resulting in the disappearance of about 40% of the >500 endemic haplochromine cichlid species (Goudswaard et al., 2008; Witte et al., 1992), with extinction falling disproportionately among pelagic piscivorous species (Fig. S21; Goldschmidt et al., 1993; McGee et al., 2015; Witte et al., 2007). While numerical abundance of cichlids has subsequently recovered among pelagic zooplanktivores and detritivores to reach pre-Nile perch numbers (van Zwieten et al., 2015; Witte et al., 2007), species diversity remains much reduced (Fig. S21) and pelagic piscivores show no signs of recovering (van Zwieten et al., 2015). The disappearance of haplochromine species occurred concurrently with increases in the native cyprinid and shrimp populations (Goudswaard et al., 2008; Ogutu-Ohwayo, 1990), shifting both community composition and food web structure within the lake. Similar shifts in food web structure occurred following Nile perch introduction into Lakes Kyoga and Nabugabo (Ogutu-Ohwayo, 1990, 1993), indicating that this may be a general trend for the introduction of these predators into native fish communities.

LT has few pelagic cichlid species in comparison to similar lakes without *Lates*, such as Lake Malawi (Fig. S21). If the Lake Victoria *Lates* invasion acts as an accurate modern analog to the process of ecosystem change which *Lates* introduction causes, we expect that LT would have had a diverse pelagic cichlid community prior to *Lates* invasion that would have been decimated with their arrival and diversification. Intriguingly, this history should be available through proxies contained in the sedimentary record of the lake; although currently available sediment cores do not extend far enough back in time to examine dynamics at this timescale (Scholz et al., 2003), this hypothesis of dramatic ecosystem change upon *Lates* introduction is testable with long sediment cores in the future (e.g., Russell et al., 2020).

### Cyclic environmental conditions may have facilitated both divergence and admixture between taxa

In tropical Africa, major tectonic and climatic events—such as continental rifting, glacial–interglacial cycles, and precessional cycles—have played a large role in shaping faunal biogeography for both aquatic and terrestrial clades. In this way, geology and climate together have led to periodic reorganization of corridors and barriers to dispersal in the region since the early Miocene (Salzburger et al., 2014; Stewart, 2001, and references therein). Although rifting began in the LT region 9–12 Mya (Cohen et al., 1993) and the lake became a truly lacustrine environment 5–6 Mya, the lake environment has undergone considerable change in both bathymetry and connectivity to other waterways during this long history (Shaban et al., 2021). LT’s tie to the Congo River via the Lukuga River outflow is common in recent times (except during lowstands such as the Last Glacial Maximum; e.g., Cohen et al., 1997; Gasse et al., 1989); however it appears that this connection is only a recent phenomenon (*∼*1.5–2.5 Mya), based on geologic and phylogeographic evidence (Flügel et al., 2015), and thus the lake may have been isolated from the Congo basin for much of its history. The re-establishment of the Lukuga River influenced the divergence of chimpanzees and bonobos in the region (Flügel et al., 2015; Langergraber et al., 2012; Prüfer et al., 2012), and our results suggest that it similarly had a strong influence on divergence in *Lates* via facilitating colonization of the Lukuga River (Koblmüller et al., 2021) and subsequently LT around 1.2 Mya. However, it is important to note that the geologic and lake level history of LT are not precisely known, and many of the timing estimates in the geologic literature depend on biologic or phylogenetic inferences (e.g., Flügel et al., 2015). Mitochondrial data (Fig. S7) further support a rapid divergence between *L. niloticus* from the Congo River, *L. stappersii*, *L. microlepis*, and a common ancestor of *L. mariae* and *L. angustifrons*, with mtDNA haplotypes found in *L. niloticus* from the Congo River remaining much more similar to the LT haplotypes than to those found in *L. niloticus* from the Niger or Nile River basins. This provides further evidence for *L. niloticus* colonization of LT from the Congo River basin, rather than from the Nile River basin earlier in the history of LT.

Patterns of lake level fluctuations over the course of LT’s long history undoubtedly influenced the evolution of its fauna, and likely were influential in the evolution of high levels of endemism within the lake (Coulter, 1991c). Within LT, considerable fault-related lake floor features (e.g., the Kavala Island Ridge and Kalya Horst; Fig. 1) create barriers to movement of lacustrine taxa at lowstands. As a result, low water levels can separate previously panmictic populations for evolutionarily relevant lengths of time, and this has likely contributed to diversification in LT’s littoral species (Coulter, 1991c; Rossiter, 1995). Although the height of these ridges separating basins would have changed over geologic time, significant water level drops likely separated the lake’s three major basins hydrologically (>400m) or physically (>500m), which could have created barriers (e.g., for pelagic species) or conduits (e.g., for littoral species) for gene flow between populations in geographically distant parts of the lake (Fig. 1; Rossiter, 1995). Allopatric divergence in *Lates* may have been facilitated by periods of basin separation. While seismic reflection data confirm major water level fluctuations in LT (Russell et al., 2020; Shaban et al., 2021), we currently have few calibrations for anchoring the timeline of these events prior to *∼*100 Kya (Scholz et al., 2007), the oldest ages of sediment cores from the lake. Yet water level fluctuations in nearby Lake Malawi (Lyons et al., 2015) and correlations with global climate phenomena (Fig. S1; Cohen et al., 2007) indicate that LT also experienced large water level fluctuations during the mid- to late-Pleistocene, concurrent with our inferred divergence times among the four LT endemic *Lates* species and introgression events involving *L. niloticus*. While there is uncertainty in the timing and magnitude of lake level changes, as well as in the timing of *Lates* divergence, this indicates a possible role for these fluctuations as a “species pump” within the *Lates*, as has been suggested for cichlid species in LT and other lakes (Duftner et al., 2005; Ivory et al., 2016; Koblmüller et al., 2005; Rossiter, 1995; Sturmbauer et al., 2017).

The most recent megadrought conditions in the African Great Lakes region occurred *∼*80–220 Kya, as evidenced by sediment cores from Lakes Malawi (which was reduced to a *∼*125m deep saline, alkaline, well mixed lake; Cohen et al., 2007; Scholz et al., 2007; Streib et al., 2024) and Tanganyika (drop >400m; Burnett et al., 2011). The most recent admixture event that we detect, between *L. niloticus* and *L. angustifrons*, appears to have occurred toward the end of these known megadrought conditions (*∼*100 Kya, Fig. 2D), and thus may relate to a time of rising lake levels that reconnected the Lukuga River to LT. The group of admixture events prior to this, both within LT and between *L. niloticus* and *L. microlepis*, are dated to the late Pleistocene, just prior to this period of extreme drought (median estimates *∼*250–270 Kya). While our confidence intervals on admixture timing for these post-divergence admixture events are wide, this uncertainty may in fact suggest that there were multiple admixture events among these taxa around 250 Kya, a time period during which LT water levels were relatively high (Fig. S1; McGlue et al., 2008; Scholz et al., 2007; Tiercelin and Mondeguer, 1991), allowing for connectivity among populations of lacustrine fauna prior to later drought conditions. Our two-taxon demographic models are consistent with the hypothesis that multiple bouts of admixture occurred between *Lates* taxa within LT (Fig. S19), but further investigations will be needed to pinpoint the timing and number of admixture events, as our current analyses are limited by the resolution of data available for inferring such events.

### Variation in admixture across the genome

Our results show substantial variation in the fate of introgressed regions across the *Lates* genomes (Fig. 3). We see strong correspondence between admixture proportions (*F_d_*) and chromosomal position, tied to variation in recombination rates across chromosomes (Fig. 3B,D). Relatedly, there is a genome-wide slightly negative relationship between recombination rate and admixture proportion (Fig. 3C), demonstrating that alleles derived from introgression are more likely to be retained in regions of low recombination, the opposite of the pattern found in similar previous studies (e.g., Martin et al., 2019; Schumer et al., 2018) and what we would expect from theory for selection against deleterious introgressed alleles (Martin and Jiggins, 2017). However, there are other examples of negative recombination-introgression relationships, such as in a North American population of *Drosophila melanogaster* with admixed European and African ancestry (Pool, 2015). In this system, it was shown that positive selection can contribute to such a negative correlation, with the strength of the effect dependent on both the number of sites under selection and the strength of selection (Duranton and Pool, 2022). This suggests the relationship between recombination rate and admixture can be informative about selection in admixed populations. These results have implications for understanding how selection and genome structure interact to influence genetic diversity across species’ genomes, and how these forces therefore influence our inferences of species histories through phylogenetic approaches. In regions of the genome with low recombination, we observe the retention of more alleles derived from introgression, and thus infer topologies consistent with this history of introgression; in contrast, regions with low retention of introgressed alleles (with high recombination) recover the species tree topology more often (Fig. 3A). However, the effect observed here is relatively weak (adjusted *R*^2^ = 0.00368). We also find a relationship between admixture extent and chromosome length, such that shorter chromosomes have a greater proportion of their length showing significant signals of admixture (Fig. 4C). These results underscore the value and increasingly, necessity, of understanding genome structure and recombination rates in order to effectively interpret species histories from genomic data, and how this may differ in systems with different introgression histories.

## Summary

Together, our analyses suggest a dynamic evolutionary history of the *Lates* radiation within LT, marked by rapid, recent, environmentally linked diversification. The colonization of the lake by these top predators in the face of an already species-rich community likely would have dramatically altered the lake’s community and food web, as has been observed via the anthropogenic introduction of related fish in other lakes in the region. While a clearer record of past lake conditions accessible through scientific drilling will clarify just how wide-ranging the effects of *Lates* colonization and diversification were, we predict that the ecosystem consequences of this recent radiation would have been substantial, and would have left the biodiversity of LT irreversibly altered.Thus, the *Lates* of LT and the broader fish assemblage in this lake represent an intriguing system for studying the effects that predators can have on biodiversity, food webs, and community structure in novel ecosystems.

## Acknowledgements

All analyses of genomic data requiring high performance computing were accomplished with an allocation from the University of Wyoming’s Advanced Research Computing Center, on its Teton (https://doi.org/10.15786/M2FY47) and Beartooth (https://doi.org/10.15786/M2FY47) Computing Environments; GNU Parallel was used to parallelize computations (Tange, 2011). Special thanks go to Benedikt Ehrenfels for coordinating field campaigns, the late Mupape Mukuli for facilitating logistics during fieldwork, and to the crew of the MV Maman Benita for invaluable fieldwork assistance. We thank the whole team at the Tanzanian Fisheries Research Institute for their support and the Tanzanian Commission for Science and Technology (COSTECH) for the permits to collect fish from Lake Tanganyika. Live fish samples were collected under the approved University of Wyoming IACUC protocol #20160602CW00241-01. We thank members of the extended Wagner Lab at the University of Wyoming for discussions of analyses and comments on the manuscript. We thank Andy Cohen for valuable comments on the manuscript, and lifelong support and enthusiasm for science in Lake Tanganyika, which has inspired so many of us to investigate its beautiful biodiversity. Bathymetry data for Lake Tanganyika maps (Fig. 1) were shared by TCarta (Denver, CO). JAR was supported by fellowships from the Data Science Center, NASA Space Grant Consortium (NASA Grant #NNX15AI08H), and Office of Graduate Education at the University of Wyoming, and sequencing was accomplished with support from the University of Wyoming INBRE Data Science Core, which is supported by an Institutional Development Award (IDeA) from the National Institute of General Medical Sciences of the National Institutes of Health under Grant #2P20GM103432. JAR was also supported by an NSF Postdoctoral Fellowship in Biology (award DBI-2109825). This work was also supported by NSF grants DEB-1556963 and DEB-2224892 to CEW.

## Author contributions

JAR and CEW designed the research; JAR, JJ, BS, MMM, EAS, IAK, and CEW performed fieldwork to collect samples; JAR generated the data; JAR, JJ, and ALL analyzed the genomic data; BS generated and analyzed the mtDNA data; JAR, OS, and CEW framed and planned the manuscript; JAR and CEW wrote the manuscript with substantial input and feedback from all other authors.

## Data and code availability

All genomic data are archived in the NCBI Sequence Read Archive, under BioProjects PRJNA776855 (genotyping-by-sequencing) and PRJNA1207083 (whole genome sequencing, *note for review: these data are currently embargoed*). Metadata and scripts used in data processing and analysis are archived on Zenodo at [TBD], and scripts can additionally be found in the GitHub repository https://github.com/jessicarick/lates-wgs-scripts.

## References

1. Adrion, J. R., Galloway, J. G., and Kern, A. D. (2020). Predicting the Landscape of Recombination Using Deep Learning. Molecular Biology and Evolution, 37(6):1790–1808.

2. Alston, J., Maitland, B., Brito, B., Esmaeili, S., Ford, A., Hays, B., Jesmer, B., Molina, F., and Goheen, J. (2019). Reciprocity in restoration ecology: When might large carnivore reintroduction restore ecosystems? Biological Conservation, 234:82–89.

3. April, J., Hanner, R. H., Dion-Côté, A.-M., and Bernatchez, L. (2013). Glacial cycles as an allopatric speciation pump in north-eastern American freshwater fishes. Molecular Ecology, 22(2).

4. Aubry, K. B., Statham, M. J., Sacks, B. N., Perrine, J. D., and Wisely, S. M. (2009). Phylogeography of the North American red fox: vicariance in Pleistocene forest refugia. Molecular Ecology, 18(12).

5. Ayoub, N. A. and Riechert, S. E. (2004). Molecular evidence for Pleistocene glacial cycles driving diversification of a North American desert spider, Agelenopsis aperta. Molecular Ecology, 13(11).

6. Barnosky, A. D., Bell, C. J., Emslie, S. D., Goodwin, H. T., Mead, J. I., Repenning, C. A., Scott, E., and Shabel, A. B. (2004). Exceptional record of mid-Pleistocene vertebrates helps differentiate climatic from anthropogenic ecosystem perturbations. Technical report.

7. Basiita, R. K., Zenger, K. R., Mwanja, M. T., and Jerry, D. R. (2018). Gene flow and genetic structure in nile perch, lates niloticus, from African freshwater rivers and lakes. PLoS ONE, 13(7):1–21.

8. Beadle, L. C. (1974). Lake Tanganyika. In The inland waters of tropical Africa. An introduction to tropical limnology, chapter 16, pages 214–229. Longman, London/New York.

9. Brown, K. J., Rüber, L., Bills, R., and Day, J. J. (2010). Mastacembelid eels support Lake Tanganyika as an evolutionary hotspot of diversification. BMC Evolutionary Biology, 10(1):188.

10. Burnett, A. P., Soreghan, M. J., Scholz, C. A., and Brown, E. T. (2011). Tropical East African climate change and its relation to global climate: A record from Lake Tanganyika, Tropical East Africa, over the past 90+ kyr. Palaeogeography, Palaeoclimatology, Palaeoecology, 303(1-4).

11. Carpenter, S. R., Kitchell, J. F., and Hodgson, J. R. (1985). Cascading Trophic Interactions and Lake Productivity. BioScience, 35(10):634–639.

12. Claramunt, S. and Cracraft, J. (2015). A new time tree reveals Earth history’s imprint on the evolution of modern birds. Science Advances, 1(11).

13. Clarke, A. and Crame, J. A. (2010). Evolutionary dynamics at high latitudes: speciation and extinction in polar marine faunas. Philosophical Transactions of the Royal Society B: Biological Sciences, 365(1558).

14. Clement, A. C., Hall, A., and Broccoli, A. J. (2004). The importance of precessional signals in the tropical climate. Climate Dynamics, 22(4):327–341.

15. Cohen, A. S., Lezzar, K. E., Tiercelin A, J. J., and Soreghan, M. (1997). New palaeogeographic and lake-level reconstructions of Lake Tanganyika: Implications for tectonic, climatic and biological evolution in a rift lake. Basin Research, 9(2):107–132.

16. Cohen, A. S., Soreghan, M. J., and Scholz, C. A. (1993). Estimating the age of formation of lakes: an example from Lake Tanganyika, East African Rift system. Geology, 21(6):511–514.

17. Cohen, A. S., Stone, J. R., Beuning, K. R. M., Park, L. E., Reinthal, P. N., Dettman, D., Scholz, C. A., Johnson, T. C., King, J. W., Talbot, M. R., Brown, E. T., and Ivory, S. J. (2007). Ecological consequences of early Late Pleistocene megadroughts in tropical Africa. Proceedings of the National Academy of Sciences, 104(42):16422–16427.

18. Coulter, G. W. (1976). The biology of Lates species (Nile perch) in Lake Tanganyika, and the status of the pelagic fishery for Lates species and Luciolates stappersii (Blgr.). Journal of Fish Biology, 9(3):235–259.

19. Coulter, G. W. (1991a). Lake Tanganyika and its Life. Oxford University Press, London.

20. Coulter, G. W. (1991b). Pelagic Fish. In Coulter, G. W., editor, Lake Tanganyika and its Life, chapter 6, pages 111–138. Oxford University Press.

21. Coulter, G. W. (1991c). Zoogeography, affinities and evolution, with special regard to the fish. In Coulter, G. W., editor, Lake Tanganyika and its Life, chapter 10, pages 275–305. Oxford University Press, London.

22. Day, J., Bills, R., and Friel, J. (2009). Lacustrine radiations in African Synodontis catfish. Journal of Evolutionary Biology, 22(4):805–817.

23. DeChaine, E. G., Wendling, B. M., and Forester, B. R. (2014). Integrating environmental, molecular, and morphological data to unravel an ice-age radiation of arctic-alpine Campanula in western North America. Ecology and Evolution, 4(20).

24. DeMoncal, P. B. (1995). Plio-Pleistocene African Climate. Science, 270(5233):53–59.

25. Duftner, N., Koblmüller, S., and Sturmbauer, C. (2005). Evolutionary Relationships of the Limnochromini, a Tribe of Benthic Deepwater Cichlid Fish Endemic to Lake Tanganyika, East Africa. Journal of Molecular Evolution, 60(3).

26. Duranton, M. and Pool, J. E. (2022). Interactions Between Natural Selection and Recombination Shape the Genomic Landscape of Introgression. Molecular Biology and Evolution, 39(7):msac122.

27. Edelman, N. B., Frandsen, P. B., Miyagi, M., Clavijo, B., Davey, J., Dikow, R. B., García-Accinelli, G., Van Belleghem, S. M., Patterson, N., Neafsey, D. E., Challis, R., Kumar, S., Moreira, G. R. P., Salazar, C., Chouteau, M., Counterman, B. A., Papa, R., Blaxter, M., Reed, R. D., Dasmahapatra, K. K., Kronforst, M., Joron, M., Jiggins, C. D., McMillan, W. O., Di Palma, F., Blumberg, A. J., Wakeley, J., Jaffe, D., and Mallet, J. (2019). Genomic architecture and introgression shape a butterfly radiation. Science, 366(6465).

28. Ehrenfels, B., Junker, J., Namutebi, D., Callbeck, C. M., Dinkel, C., Kalangali, A., Kimirei, I. A., Mbonde, A. S., Mosille, J. B., Sweke, E. A., Schubert, C. J., Seehausen, O., Wagner, C. E., and Wehrli, B. (2023). Isotopic signatures induced by upwelling reveal regional fish stocks in Lake Tanganyika. PLOS ONE, 18(11):e0281828.

29. Ellis, C. M. A. (1978). Biology of Luciolates stappersi in Lake Tanganyika (Burundi). Transactions of the American Fisheries Society, 107(4):557–566.

30. Estes, J. A., Terborgh, J., Brashares, J. S., Power, M. E., Berger, J., Bond, W. J., Carpenter, S. R., Essington, T. E., Holt, R. D., Jackson, J. B. C., Marquis, R. J., Oksanen, L., Oksanen, T., Paine, R. T., Pikitch, E. K., Ripple, W. J., Sandin, S. a., Scheffer, M., Schoener, T. W., Shurin, J. B., Sinclair, A. R. E., Soulé, M. E., Virtanen, R., and Wardle, D. a. (2011). Trophic downgrading of planet Earth. Science, 333:301–306.

32. Excoffier, L., Dupanloup, I., Huerta-Sánchez, E., Sousa, V. C., and Foll, M. (2013). Robust Demographic Inference from Genomic and SNP Data. PLoS Genetics, 9(10).

33. Flügel, T. J., Eckardt Frank, F. D., and Cotterill, F. P. (2015). The present day drainage patterns of the Congo river system and their Neogene evolution. In Geology and Resource Potential of the Congo Basin, pages 315–337. Springer Berlin Heidelberg.

34. Gasse, F., Lédée, V., Massault, M., and Fontes, J.-C. (1989). Water-level fluctuations of Lake Tanganyika in phase with oceanic changes during the last glaciation and deglaciation. Nature, 342(6245):57–59.

35. Goldschmidt, T., Witte, F., and Wanink, J. (1993). Cascading Effects of the Introduced Nile Perch on the Detritivorous/ Phytoplanktivorous Species in the Sublittoral Areas of Lake Victoria Cascading Effects of the Introduced Nile Perch on the Detritivorous/Phytoplanktivorous Species in the Sublittoral Areas of Lake Victoria TIJS GOLDSCHMIDT. Technical Report 3.

36. Goudswaard, K., Witte, F., and Katunzi, E. F. (2008). The invasion of an introduced predator, Nile perch (Lates niloticus, L.) in Lake Victoria (East Africa): Chronology and causes. Environmental Biology of Fishes, 81(2):127–139.

37. Groombridge, B. and Jenkins, M. (1998). Freshwater biodiversity: a preliminary global assessment. Technical report, United Nationals Environmental Programme - World Conservation Montitoring Centre, Cambridge, UK.

38. He, Z., Li, X., Yang, M., Wang, X., Zhong, C., Duke, N. C., Wu, C.-I., and Shi, S. (2019). Speciation with gene flow via cycles of isolation and migration: insights from multiple mangrove taxa. National Science Review, 6(2).

39. Hecky, R. E., Mugidde, R., Ramlal, P. S., Talbot, M. R., and Kling, G. W. (2010). Multiple stressors cause rapid ecosystem change in Lake Victoria. Freshwater Biology, 55(SUPPL. 1):19–42.

40. Irisarri, I., Singh, P., Koblmüller, S., Torres-Dowdall, J., Henning, F., Franchini, P., Fischer, C., Lemmon, A. R., Lemmon, E. M., Thallinger, G. G., Sturmbauer, C., and Meyer, A. (2018). Phylogenomics uncovers early hybridization and adaptive loci shaping the radiation of Lake Tanganyika cichlid fishes. Nature Communications, 9(1).

41. Ivory, S. J., Blome, M. W., King, J. W., McGlue, M. M., Cole, J. E., and Cohen, A. S. (2016). Environmental change explains cichlid adaptive radiation at Lake Malawi over the past 1.2 million years. Proceedings of the National Academy of Sciences of the United States of America, 113(42):11895–11900.

42. Koblmüller, S., Duftner, N., Katongo, C., Phiri, H., and Sturmbauer, C. (2005). Ancient Divergence in Bathypelagic Lake Tanganyika Deepwater Cichlids: Mitochondrial Phylogeny of the Tribe Bathybatini. Journal of Molecular Evolution, 60(3):297–314.

43. Koblmüller, S., Schöggl, C. A., Lorber, C. J., Van Steenberge, M., Kmentová, N., Vanhove, M. P., and Zangl, L. (2021). African lates perches (Teleostei, Latidae, Lates): Paraphyly of Nile perch and recent colonization of Lake Tanganyika. Molecular Phylogenetics and Evolution, 160.

44. Koel, T. M., Tronstad, L. M., Arnold, J. L., Gunther, K. A., Smith, D. W., Syslo, J. M., and White, P. J. (2019). Predatory fish invasion induces within and across ecosystem effects in Yellowstone National Park. Science Advances, 5(3).

45. Kondo, T. and Abe, N. (1995). Habitat preference, food habits and growth of juveniles of Lates angustifrons and Lates mariae (Pisces:Centropomidae) in Lake Tanganyika. Ecological Research, 10(October 1988):275–280.

46. Korneliussen, T. S., Albrechtsen, A., and Nielsen, R. (2014). ANGSD: Analysis of next generation sequencing data. BMC Bioinformatics, 15:356.

47. Langergraber, K. E., Prufer, K., Rowney, C., Boesch, C., Crockford, C., Fawcett, K., Inoue, E., Inoue-Muruyama, M., Mitani, J. C., Muller, M. N., Robbins, M. M., Schubert, G., Stoinski, T. S., Viola, B., Watts, D., Wittig, R. M., Wrangham, R. W., Zuberbuhler, K., Paabo, S., and Vigilant, L. (2012). Generation times in wild chimpanzees and gorillas suggest earlier divergence times in great ape and human evolution. Proceedings of the National Academy of Sciences, 109(39).

48. Levin, N. E. (2015). Environment and Climate of Early Human Evolution. Annual Review of Earth and Planetary Sciences, 43(1):405–429.

49. Linck, E. and Battey, C. J. (2019). On the relative ease of speciation with periodic gene flow. bioRxiv, pages 1–33.

50. Lyons, R. P., Scholz, C. A., Cohen, A. S., King, J. W., Brown, E. T., Ivory, S. J., Johnson, T. C., Deino, A. L., Reinthal, P. N., Mcglue, M. M., and Blome, M. W. (2015). Continuous 1.3-million-year record of East African hydroclimate, and implications for patterns of evolution and biodiversity. Proceedings of the National Academy of Sciences, 112(51):2–7.

51. Malinsky, M., Matschiner, M., and Svardal, H. (2021). Dsuite - Fast D-statistics and related admixture evidence from VCF files. Molecular Ecology Resources, 21(2):584–595.

52. Malinsky, M., Svardal, H., Tyers, A. M., Miska, E. A., Genner, M. J., Turner, G. F., and Durbin, R. (2018). Whole-genome sequences of Malawi cichlids reveal multiple radiations interconnected by gene flow. Nature Ecology and Evolution, 2(12):1940–1955.

53. Mannini, P., Katonda, I., Kissaka, B., and Verburg, P. (1999). Feeding ecology of Lates stappersii in Lake Tanganyika. Hydrobiologia, 407(0):131–139.

54. Martin, M., Patterson, M., Garg, S., Fischer, S. O., Pisanti, N., Klau, G. W., Schöenhuth, A., and Marschall, T. (2016). WhatsHap: fast and accurate read-based phasing. bioRxiv.

55. Martin, S. H. (2026). A model-free method for genealogical inference without phasing and its application for topology weighting. Genetics, 232(1):iyaf181.

56. Martin, S. H., Davey, J. W., and Jiggins, C. D. (2015). Evaluating the use of ABBA-BABA statistics to locate introgressed loci. Molecular Biology and Evolution, 32(1):244–257.

57. Martin, S. H., Davey, J. W., Salazar, C., and Jiggins, C. D. (2019). Recombination rate variation shapes barriers to introgression across butterfly genomes. PLOS Biology, 17(2):e2006288.

58. Martin, S. H. and Jiggins, C. D. (2017). Interpreting the genomic landscape of introgression.

59. Martin, S. H. and Van Belleghem, S. M. (2017). Exploring Evolutionary Relationships Across the Genome Using Topology Weighting. Genetics, 206:429–438.

60. McGee, M. D., Borstein, S. R., Meier, J. I., Marques, D. A., Mwaiko, S., Taabu, A., Kishe, M. A., O’Meara, B., Bruggmann, R., Excoffier, L., and Seehausen, O. (2020). The ecological and genomic basis of explosive adaptive radiation. Nature, 586(7827):75–79.

61. McGee, M. D., Borstein, S. R., Neches, R. Y., Buescher, H. H., Seehausen, O., and Wainwright, P. C. (2015). A pharyngeal jaw evolutionary innovation facilitated extinction in Lake Victoria cichlids. Science, 350(6264).

62. McGlue, M. M., Lezzar, K. E., Cohen, A. S., Russell, J. M., Tiercelin, J. J., Felton, A. A., Mbede, E., and Nkotagu, H. H. (2008). Seismic records of late Pleistocene aridity in Lake Tanganyika, tropical East Africa. Journal of Paleolimnology, 40(2):635–653.

63. Meier, J. I., Marques, D. A., Mwaiko, S., Wagner, C. E., Excoffier, L., and Seehausen, O. (2017). Ancient hybridization fuels rapid cichlid fish adaptive radiations. Nature Communications, 8(1).

64. Meier, J. I., McGee, M. D., Marques, D. A., Mwaiko, S., Kishe, M., Wandera, S., Neumann, D., Mrosso, H., Chapman, L. J., Chapman, C. A., Kaufman, L., Taabu-Munyaho, A., Wagner, C. E., Bruggmann, R., Excoffier, L., and Seehausen, O. (2023). Cycles of fusion and fission enabled rapid parallel adaptive radiations in African cichlids. Science, 381(6665).

65. Milec, L. J. M., Vanhove, M. P. M., Bukinga, F. M., De Keyzer, E. L. R., Kapepula, V. L., Masilya, P. M., Mulimbwa, N., Wagner, C. E., and Raeymaekers, J. A. M. (2022). Complete mitochondrial genomes and updated divergence time of the two freshwater clupeids endemic to Lake Tanganyika (Africa) suggest intralacustrine speciation. BMC Ecology and Evolution, 22(1):127.

66. Mooney, H. A. and Cleland, E. E. (2001). The evolutionary impact of invasive species. Proceedings of the National Academy of Sciences, 98(10):5446–5451.

67. Moran, B. M., Payne, C., Langdon, Q., Powell, D. L., Brandvain, Y., and Schumer, M. (2021). The genomic consequences of hybridization. eLife, 10.

68. Murray, A. M. and Attia, Y. S. (2004). A new species of Lates (Teleostei: Perciformes) from the Lower Oligocene of Egypt. Journal of Vertebrate Paleontology, 24(2):299–308.

69. Mwanja, M. T., Muwanika, V., Masembe, C., Nyakaana, S., and Mwanja, W. W. (2013). Evolutionary history of nile perch lates sp. inferred from mitochondrial DNA variation analyses. Zoological Studies, 52(1):1–12.

70. Mwanja, M. T., Muwanika, V., Nyakaana, S., Masembe, C., Mbabazi, D., Rutasire, J., and Mwanja, W. W. (2011). Population morphological variation of Nile perch (Lates niloticus, L. 1758), of East African Lakes and their associated waters. African Journal of Environmental Science and Technology, 5(11):941–949.

71. Nevado, B., Contreras-Ortiz, N., Hughes, C., and Filatov, D. A. (2018). Pleistocene glacial cycles drive isolation, gene flow and speciation in the high-elevation Andes. New Phytologist, 219(2).

72. Ogutu-Ohwayo, R. (1990). Changes in the prey ingested and the variations in the Nile perch and other fish stocks of Lake Kyoga and the northern waters of Lake Victoria (Uganda). Journal of Fish Biology, 37(1).

73. Ogutu-Ohwayo, R. (1993). The Effects of Predation by Nile Perch, Lates niloticus L., on the Fish of Lake Nabugabo, with Suggestions for Conservation of Endangered Endemic Cichlids. Conservation Biology, 7(3).

74. Patterson, N., Moorjani, P., Luo, Y., Mallick, S., Rohland, N., Zhan, Y., Genschoreck, T., Webster, T., and Reich, D. (2012). Ancient admixture in human history. Genetics, 192(3):1065–1093.

75. Peart, C. R., Bills, R., Wilkinson, M., and Day, J. J. (2014). Nocturnal claroteine catfishes reveal dual colonisation but a single radiation in Lake Tanganyika. Molecular Phylogenetics and Evolution, 73:119–128.

76. Pool, J. E. (2015). The Mosaic Ancestry of the *Drosophila* Genetic Reference Panel and the *D. melanogaster* Reference Genome Reveals a Network of Epistatic Fitness Interactions. Molecular Biology and Evolution, page msv194.

77. Prüfer, K., Munch, K., Hellmann, I., Akagi, K., Miller, J. R., Walenz, B., Koren, S., Sutton, G., Kodira, C., Winer, R., Knight, J. R., Mullikin, J. C., Meader, S. J., Ponting, C. P., Lunter, G., Higashino, S., Hobolth, A., Dutheil, J., Karakoç, E., Alkan, C., Sajjadian, S., Catacchio, C. R., Ventura, M., Marques-Bonet, T., Eichler, E. E., André, C., Atencia, R., Mugisha, L., Junhold, J., Patterson, N., Siebauer, M., Good, J. M., Fischer, A., Ptak, S. E., Lachmann, M., Symer, D. E., Mailund, T., Schierup, M. H., Andrés, A. M., Kelso, J., and Pääbo, S. (2012). The bonobo genome compared with the chimpanzee and human genomes. Nature, 486(7404).

78. Ravinet, M., Faria, R., Butlin, R. K., Galindo, J., Bierne, N., Rafajlović, M., Noor, M. A. F., Mehlig, B., and Westram, A. M. (2017). Interpreting the genomic landscape of speciation: a road map for finding barriers to gene flow. Journal of Evolutionary Biology, 30(8):1450–1477.

79. Rick, J. A., Junker, J., Kimirei, I. A., Sweke, E. A., Dinkel, C., Mwaiko, S., Seehausen, O., and Wagner, C. E. (2021). The genetic population structure of Lake Tanganyika’s Lates 1 species flock, an endemic radiation of pelagic top predators 2 3. *bioRxiv*.

80. Rick, J. A., Junker, J., Kimirei, I. A., Sweke, E. A., Mosille, J. B., Dinkel, C., Mwaiko, S., Seehausen, O., and Wagner, C. E. (2022). The genetic population structure of Lake Tanganyika’s Lates species flock, an endemic radiation of pelagic top predators. Journal of Heredity, (10.1093/jhered/esab072).

81. Ronco, F., Matschiner, M., Böhne, A., Boila, A., Büscher, H. H., El Taher, A., Indermaur, A., Malinsky, M., Ricci, V., Kahmen, A., Jentoft, S., and Salzburger, W. (2021). Drivers and dynamics of a massive adaptive radiation in cichlid fishes. Nature, 589(7840):76–81.

82. Rossiter, A. (1995). The Cichlid Fish Assemblages of Lake Tanganyika: Ecology, Behaviour and Evolution of its Species Flocks.

83. Russell, J. M., Barker, P., Cohen, A., Ivory, S., Kimirei, I., Lane, C., Leng, M., Maganza, N., McGlue, M., Msaky, E., Noren, A., Park Boush, L., Salzburger, W., Scholz, C., Tiedemann, R., and Nuru, S. (2020). ICDP workshop on the Lake Tanganyika Scientific Drilling Project: a late Miocene–present record of climate, rifting, and ecosystem evolution from the world’s oldest tropical lake. Scientific Drilling, 27:53–60.

84. Salzburger, W., Van Bocxlaer, B., and Cohen, A. S. (2014). Ecology and Evolution of the African Great Lakes and Their Faunas. Annual Review of Ecology, Evolution, and Systematics, 45(1):519–545.

85. Schiffels, S. and Durbin, R. (2014). Inferring human population size and separation history from multiple genome sequences. Nature Genetics, 46(8):919–25.

86. Schmitz, O. J., Krivan, V., and Ovadia, O. (2004). Trophic cascades: The primacy of trait-mediated indirect interactions.

87. Scholz, C. A., Johnson, T. C., Cohen, A. S., King, J. W., Peck, J. A., Overpeck, J. T., Talbot, M. R., Brown, E. T., Kalindekafe, L., Amoako, P. Y. O., Lyons, R. P., Shanahan, T. M., Castañeda, I. S., Heil, C. W., Forman, S. L., McHargue, L. R., Beuning, K. R. M., Gomez, J., and Pierson, J. (2007). East African megadroughts between 135 and 75 thousand years ago and bearing on early-moderns human origins. Proceedings of the National Academy of Sciences, 104(42):16416–16421.

88. Scholz, C. A., King, J. W., Ellis, G. S., Swart, P. K., Stager, J. C., and Colman, S. M. (2003). Paleolimnology of Lake Tanganyika, East Africa, over the past 100 k yr. Journal of Paleolimnology, 30:139–150.

89. Schumer, M., Xu, C., Powell, D. L., Durvasula, A., Skov, L., Holland, C., Blazier, J. C., and Sankararaman, S. (2018). Natural selection interacts with recombination to shape the evolution of hybrid genomes. Science, 660(May):656–660.

90. Seehausen, O. (1999). A reconsideration o fthe ecological composition of the cichlid species flock in Lake Victoria before and after the Nile perch boom. In van Densen, W. and Morris, M., editors, Fish and fisheries of lakes and reservoirs in Southeast Asia and Africa, pages 281–293. Westbury, West Yorkshire.

91. Seehausen, O. and Wagner, C. E. (2014). Speciation in Freshwater Fishes. Annual Review of Ecology, Evolution, and Systematics, 45:621–651.

92. Shaban, S. N., Scholz, C. A., Muirhead, J. D., and Wood, D. A. (2021). The stratigraphic evolution of the Lake Tanganyika Rift, East Africa: Facies distributions and paleo-environmental implications. Palaeogeography, Palaeoclimatology, Palaeoecology, 575.

93. Stamatakis, A. (2014). RAxML version 8: A tool for phylogenetic analysis and post-analysis of large phylogenies. Bioinformatics, 30(9):1312–1313.

94. Stewart, K. M. (2001). The freshwater fish of Neogene Africa (Miocene-Pleistocene): systematics and biogeography. Fish and Fisheries, 2(3).

95. Streib, L. C., Armitage, S. J., and Scholz, C. A. (2024). Using luminescence dating to constrain lake sediment records: A new age model for the 1.38 Ma lake Malawi drill core, Eastern Africa. Quaternary Science Reviews, 334:108691.

96. Sturmbauer, C., Börger, C., Van Steenberge, M., and Koblmüller, S. (2017). A separate lowstand lake at the northern edge of Lake Tanganyika? Evidence from phylogeographic patterns in the cichlid genus Tropheus. Hydrobiologia, 791(1):51–68.

97. Svardal, H., Quah, F. X., Malinsky, M., Ngatunga, B. P., Miska, E. A., Salzburger, W., Genner, M. J., Turner, G. F., and Durbin, R. (2020). Ancestral Hybridization Facilitated Species Diversification in the Lake Malawi Cichlid Fish Adaptive Radiation. Molecular Biology and Evolution, 37(4).

98. Svardal, H., Salzburger, W., and Malinsky, M. (2021). Genetic Variation and Hybridization in Evolutionary Radiations of Cichlid Fishes. Annual Review of Animal Biosciences, 9(1):55–79.

99. Tange, O. (2011). GNU Parallel - The Command-Line Power Tool. The USENIX Magazine, pages 42–47.

100. Tiercelin, J. J. and Mondeguer, A. (1991). The geology of the Tanganyika Trough. In Coulter, G. W., editor, Lake Tanganyika and its Life, page 7–48. Oxford University Press.

101. Tierney, J. E., Russell, J. M., Huang, Y., Damste, J. S. S., Hopmans, E. C., and Cohen, A. S. (2008a). Northern Hemisphere Controls on Tropical Southeast African Climate During the Past 60,000 Years. Science, 322(5899).

102. Tierney, J. E., Russell, J. M., Huang, Y., Sinninghe Damsté, J. S., Hopmans, E. C., and Cohen, A. S. (2008b). Northern hemisphere controls on tropical southeast African climate during the past 60,000 years. Science, 322(5899):252–255.

103. Trauth, M. H., Maslin, M. A., Deino, A. L., Junginger, A., Lesoloyia, M., Odada, E. O., Olago, D. O., Olaka, L. A., Strecker, M. R., and Tiedemann, R. (2010). Human evolution in a variable environment: the amplifier lakes of Eastern Africa. Quaternary Science Reviews, 29(23-24):2981–2988.

104. Trauth, M. H., Maslin, M. A., Deino, A. L., Strecker, M. R., Bergner, A. G., and Dühnforth, M. (2007). High- and low-latitude forcing of Plio-Pleistocene East African climate and human evolution. Journal of Human Evolution, 53(5):475–486.

105. Tricou, T., Tannier, E., and de Vienne, D. M. (2022). Ghost Lineages Highly Influence the Interpretation of Introgression Tests. Systematic Biology, 71(5):1147–1158.

106. van Zwieten, P., Kolding, J., Plank, M., Hecky, R. E., Bridgeman, T., MacIntyre, S., Seehausen, O., and Silsbe, G. (2015). The Nile perch invasion in Lake Victoria: cause or consequence of the haplochromine decline? Canadian Journal of Fisheries and Aquatic Sciences, 73(4):622–643.

107. Verschuren, D., Sinninghe Damsté, J. S., Moernaut, J., Kristen, I., Blaauw, M., Fagot, M., and Haug, G. H. (2009). Half-precessional dynamics of monsoon rainfall near the East African Equator. Nature, 462(7273).

108. Vij, S., Kuhl, H., Kuznetsova, I. S., Komissarov, A., Yurchenko, A. A., Van Heusden, P., Singh, S., Thevasagayam, N. M., Prakki, S. R. S., Purushothaman, K., Saju, J. M., Jiang, J., Mbandi, S. K., Jonas, M., Hin Yan Tong, A., Mwangi, S., Lau, D., Ngoh, S. Y., Liew, W. C., Shen, X., Hon, L. S., Drake, J. P., Boitano, M., Hall, R., Chin, C. S., Lachumanan, R., Korlach, J., Trifonov, V., Kabilov, M., Tupikin, A., Green, D., Moxon, S., Garvin, T., Sedlazeck, F. J., Vurture, G. W., Gopalapillai, G., Kumar Katneni, V., Noble, T. H., Scaria, V., Sivasubbu, S., Jerry, D. R., O’Brien, S. J., Schatz, M. C., Dalmay, T., Turner, S. W., Lok, S., Christoffels, A., and Orbán, L. (2016). Chromosomal-Level Assembly of the Asian Seabass Genome Using Long Sequence Reads and Multi-layered Scaffolding. PLoS Genetics, 12(4):1–35.

109. Wagner, C. E., Harmon, L. J., and Seehausen, O. (2012). Ecological opportunity and sexual selection together predict adaptive radiation. Nature, 487(7407):366–369.

110. Wainright, C. A., Muhlfeld, C. C., Elser, J. J., Bourret, S. L., and Devlin, S. P. (2021). Species invasion progressively disrupts the trophic structure of native food webs. Proceedings of the National Academy of Sciences, 118(45):e2102179118.

111. Weir, J. T., Haddrath, O., Robertson, H. A., Colbourne, R. M., and Baker, A. J. (2016). Explosive ice age diversification of kiwi. Proceedings of the National Academy of Sciences, 113(38):E5580–E5587.

112. Weir, J. T. and Schluter, D. (2004). Ice sheets promote speciation in boreal birds. Proceedings of the Royal Society of London. Series B: Biological Sciences, 271(1551).

113. Witte, F., Goldschmidt, T., Wanink, J., Van Oijen, M., Goudswaard, K., Witte-Maas’, E., and Bouton’, N. (1992). The destruction of an endemic species flock : quantitative data on the decline of the haplochromine cichlids of Lake Victoria. Technical report.

114. Witte, F., Wanink, J. H., Kishe-Machumu, M., Mkumbo, O. C., Goudswaard, P. C., and Seehausen, O. (2007). Differential decline and recovery of haplochromine trophic groups in the Mwanza Gulf of Lake Victoria. Aquatic Ecosystem Health & Management, 10(4).

115. Zhang, C., Rabiee, M., Sayyari, E., and Mirarab, S. (2018). ASTRAL-III: Polynomial time species tree reconstruction from partially resolved gene trees. BMC Bioinformatics, 19(Suppl 6):15–30.

