## Supplementary Material for "Recent rapid radiation of apex predators suggests dramatic biodiversity turnover in an ancient lake"

### Supplementary Methods

**Sampling and DNA sequencing.** Between 2001 and 2018, tissue samples (fin clips) were collected from *Lates* along the Tanzanian shore of Lake Tanganyika, as described in Rick et al. (2021). Fish were predominantly collected opportunistically from fishermen at fishing grounds and landing sites. Fish were identified in the field by sampling personnel following United Nations Food and Agricultural Organization (FAO) identification guidelines (Eccles, 1992) and guidance from local fishermen. The species can be difficult to distinguish morphologically, particularly in the juvenile stage, and field-based identifications were checked using genetic identification, as described in Rick et al. (2021).

In addition to our own collections, we obtained tissues from two *L. niloticus* collected in Cameroon from the Cornell University Museum of Vertebrates (CUMV Catalog No. 93501 and 90829) and one *L. niloticus* collected in the Democratic Republic of the Congo from the American Museum of Natural History (AMNH Catalog No. 216682). We additionally downloaded the chromosome-level reference genome assembly of one *L. calcarifer* (barramundi) individual (v3; Vij et al., 2016) and the corresponding 100bp PE `fastq` reads from NCBI (SRR3140997). This genome was used as a reference for assembly, and the reads from this individual were included as an outgroup for phylogenetic analyses.

Fin clips were stored in 95% ethanol or RNAlater until extraction. We extracted DNA from fin clips using Qiagen DNeasy Blood and Tissue Kits (Qiagen, Inc.) following the standard protocol, with the addition of an RNAse A incubation step.

***Genotyping-by-sequencing.*** Samples were sequenced using a genotyping-by-sequencing (GBS) protocol, as described in Rick et al. (2021). Briefly, we prepared libraries following protocols outlined in Parchman et al. (2012), using MseI and EcoRI restriction enzymes. Digested and barcoded fragments were amplified via PCR and then pooled for sequencing in two libraries. The prepared libraries were size-selected for 200–350bp fragments using Blue Pippin (Sage Science, MA). Each library contained ~192 individuals, and select individuals were duplicated across libraries to check for library compatibility.

We processed raw sequencing reads by first filtering common contaminants and unwanted sequences (PhiX, E. coli, and leftover barcodes, primers, and adapters from library preparation) from our data using `bowtie2` (v2.3.4.1; Langmead and Salzberg, 2012). We then matched sequence barcodes to individual fish and assembled all data to the *L. calcarifer* reference genome (v3; Vij et al., 2016) using `bwa mem` (v0.7.17; Li and Durbin, 2009) with default settings. Following alignment, we discarded any individuals with <50,000 high-quality reads assembled to the reference genome. We then identified variable sites in the assembly using `samtools` and `bcftools` (v1.8; Li et al., 2009), omitting indels and keeping only high-quality biallelic variant sites (`QUAL >20` and `GQ >9`). We filtered SNPs by minor allele frequency (`-maf 0.01`) and amount of missing data (`-max-missing 0.5`) using `vcftools` (v0.1.14; Danecek et al., 2011), in addition to only calling SNPs at sites with a minimum read depth of 10.

***Whole genome resequencing.*** In addition to the GBS dataset, we generated a set of whole genome resequencing data for 3 *L. stappersii*, 3 *L. mariae*, 3 *L. angustifrons*, 3 *L. microlepis*, and 1 *L.*

*niloticus* individuals. Tissues for these individuals were sent to HudsonAlpha Discovery (Huntsville, AL) for high molecular weight DNA extraction, PCR-free library preparation, multiplexing, and 150bp paired-end sequencing. For library preparation, samples were sonicated on the Covaris LE220 using a protocol designed to achieve a target insert size of 550 bp. The samples were then prepared using the KAPA HyperPrep whole genome protocol and dual indices. Nine of the samples were sequenced across 3 lanes of Illumina HiSeq X (150bp PE), and four were sequenced on 10% each of an Illumina NovaSeq S4 flowcell (150bp PE), resulting in ∼40x and ∼100x coverage at sequenced sites, respectively.

Short reads from whole genome sequencing were prepared in a similar manner to the GBS reads. First, read quality was checked using `FastQC` (v0.11.19; https://www.bioinformatics.babraham.ac.uk/projects/fastqc/). Then, common contaminants and unwanted sequences were removed from the data using `bowtie2`, followed by assembling the remaining reads to the *L. calcarifer* reference genome using `bwa mem` with default settings. We then created an `mpileup` with all 13 individuals using `samtools` and created three separate filtered datasets using `vcftools`, appropriate for the different types of analyses: (1) a whole genome dataset, where we filtered for missing data but kept both variable and invariant sites; (2) a full SNP dataset, where we filtered for biallelic sites, <50% missing data (`-max-missing 0.5`), and minor allele frequency >0.01 (`-maf 0.01`); and (3) a linkage-filtered SNP dataset, where we filtered for biallelic sites, <20% missing data, and minor allele frequency >0.01. We additionally filtered all datasets for sites with a minimum read depth ≥30 (roughly 1.5x the interquartile range of read depths across sites).

***Mitochondrial DNA sequencing.*** A subset of samples were selected for mitochondrial sequencing. We amplified the mitochondrial 16s region using primers 16sL2510 (CGC CTG TTT ATC AAA AAC AT) and 16sH3058 (CCG GTC TGA ACT CAG ATC ACG T), as well as internal sequencing primers 16s135F (GCA ATA GAV AWA GTA CCG CAA GGG) and 16s1543R (AAC TGA CCT GGA TTR CTC CGG TC) (Li et al., 2011). The PCR products were visualized on agarose gels to check the quality and size of amplification before sequencing. Sequencing was carried out by the Cornell University Biotechnology Resource Center (Ithaca, NY). Standard chromatographic curves of forward and reverse sequences were imported into the program `Geneious` (https://www.geneious.com) and manually aligned and edited. Consensus sequences were then combined with 16s *Lates* sequences downloaded from NCBI GenBank (https://www.ncbi.nlm.nih.gov/genbank/), and aligned in `Geneious` using MAFFT sequence alignment with default parameters for DNA. We imported alignments were into `PopART` (Leigh and Bryant, 2015) and constructed a median-joining haplotype network (Bandelt et al., 1999) using an epsilon value of 0 to visualize haplotype relationships.

**Genetic species affinities.** Because we did not have definitive identification to species for every specimen in our GBS analyses, we used hierarchical Bayesian clustering in the program `entropy` (Gompert et al., 2014; Shastry et al., 2021) to determine species identity for each specimen, as detailed in Rick et al. (2021). For the full set of Lake Tanganyika *Lates* GBS individuals filtered for 50% missing data and a minor allele frequency of 0.01, we ran `entropy` for K=4, and used the results to assign individuals to the group for which group membership ($q$) was >0.6. We assigned each of these inferred groups to a species using individuals for which we had definitive species assignments based on morphology.

**Phylogenetic relationships.** We used multiple approaches to infer phylogenetic relationships among the taxa included in this study. First, we estimated a maximum-likelihood (ML) phylogenetic tree from the concatenated GBS data using the hybrid, MPI/PThreads version of `RAxML v8.1.17`

(Stamatakis, 2014). We concatenated all variant sites that had <50% missing data and a minor allele frequency >0.01, and used the Lewis correction for invariant sites with the ASC_GTRGAMMA model of molecular evolution (Lewis 2001 ) within `RAxML` to construct the maximum likelihood phylogeny, including *L. calcarifer* as an outgroup. We additionally estimated the maximum-likelihood phylogeny from the concatenated WGS dataset containing all SNPs with <50% missing data and minor allele frequency >0.05, again using the ASC_GTRGAMMA model and Lewis ascertainment bias correction. For each dataset, 100 bootstrap replicates were used to infer nodal support.

We estimated second phylogenetic tree using the multi-species coalescent with the program `SNAPP` (Bryant et al., 2012) using the whole genome data. For `SNAPP` analyses, we started with the linkage-filtered WGS data set and thinned randomly to one SNP per 10,000bp. We tuned operators and priors in `SNAPP` to produce effective sample sizes of at least 200 for all parameters and ran each chain for 10 million steps while sampling every 1,000 steps following a burn-in of 500,000 steps. The first 20% of samples were treated as burn-in and removed before individual runs were combined in `logcombiner`. Stationarity and chain convergence were assessed using the program `Tracer` (v1.7.1, Rambaut et al., 2018).

We further generated a phylogenetic tree from individual gene trees using `ASTRAL-III` (v5.7.1, Zhang et al., 2018). `ASTRAL` finds the species tree that has the maximum number of shared induced quartet trees with the set of gene trees, and is statistically consistent under the multi-species coalescent. Gene trees were inferred using `RAxML` for 4,000bp windows across the genome (118,707 windows), again using the ASC_GTRGAMMA model of molecular evolution and the Lewis correction for invariant sites. We inferred the species tree in `ASTRAL` using the bootstrap gene trees, and calculated local posterior probabilities (Sayyari and Mirarab, 2016) and quartet support for each node.

**Diversity and divergence estimates.** From the whole genome data, we generated an "all sites" dataset containing both variant and invariant sites, retaining those sites with <20% missing data and genotype calls with a minimum read depth >10, resulting in a dataset with 25,680,832 sites. We then used `pixy` (?; v2.0.0) to estimate nucleotide diversity ($\pi$), Watterson's $\theta$, and Tajima's $D$ for each species in 10kb windows. We then estimated Weir–Cockerham $F_{ST}$ and $d_{xy}$ in 10kb windows for each pair of species using `pixy`. Using these windowed estimates, we calculated genome-wide estimates for each species ($\pi$, $\theta$) or each species pair ($F_{ST}$, $d_{xy}$).

**Estimating $N_e$ and divergence timing.** To estimate effective population sizes ($N_e$) through time and approximate dates for divergence between taxa, we used `MSMC2` (Schiffels and Durbin, 2014). We generated input files for individual samples using `samtools` and custom scripts from the `msmc-tools` repository (https://github.com/stschiff/msmc-tools), filtering for a minimum read depth of 10 and minimum mapping quality of 20. In addition, a mappability mask was generated for the *L. calcarifer* genome using the `SNPable regions` program (http://lh3lh3.users.sourceforge.net/snpable.shtml) to select only regions to which short reads could be uniquely assembled using 35bp $k$-mers and a stringency of 50% ($\sim$89.6 % of the genome). Variant sites were phased using `WhatsHap` (v1.0; Martin et al., 2016) and input files generated using scripts from the `msmc-tools` repository. Input files were generated for each sample individually (for $N_e$ estimation), for each species, and for each pair of individuals (for cross-coalescence). For individuals and species, we used a time pattern of $1*3+3*2+30*1+3*2+1*3$ and a maximum of 100 expectation maximization iterations. For cross-coalescence analyses, we reduced the time patterning to $1*3+3*2+20*1+3*2+1*3$ to increase the number of SNPs in each time segment. For runs that had problems with overfitting (11 of 78 comparisons), we further reduced the time patterning to $1*3+3*2+20*1+3*2+1*3$.

Coalescent-scaled units were scaled to biological units using $\mu = 3.5x10^{-9}$ (estimate from Lake Malawi cichlids; Malinsky et al., 2018) per generation and a mean generation time of 3 years [estimated from data in Froese and Pauly (2021); see *Calculating generation time* section below]. To estimate uncertainty in our $N_e$ estimates, we performed 50 bootstrap replicates for each individual and each species. We calculated the divergence time between a pair of taxa as the mean $T_{50}$, the time at which the cross-coalescence rate reaches 50%, for all combinations of individuals belonging to those species.

**Identifying and characterizing introgression.** Analysis of gene flow must take ILS into account because both processes generate gene trees that are incongruent with the species tree. The $D$-statistic (or ABBA-BABA statistic) is a useful and widely applied parsimony-like method for detecting gene flow despite the existence of ILS (Green et al., 2010). In a comparison of taxa (((P1,P2),P3),O), a positive value represents an excess of "ABBA" alleles, indicating excess allele sharing between P2 and P3; a negative value represents an excess of "BABA" alleles, indicating excess allele sharing between P1 and P3, compared to the null expectation under ILS only. We used `Dsuite` (v0.4; Malinsky et al., 2021) to calculate genome-wide $D$-statistics for all three-taxon combinations of *L. niloticus, L. angustifrons, L. mariae, L. stappersii,* and *L. microlepis*, again always keeping *L. calcarifer* as the outgroup (O). Significant values of $D$ can represent either the wrong tree or admixture between those taxa indicated to have an excess of shared alleles; these two hypotheses can be distinguished by testing all possible combinations of taxa.

To determine the extent of introgression and the likely location of introgression events on the *Lates* phylogeny, we used `Dsuite` to calculate $f4$-ratio statistics (Patterson et al., 2012) and, from these, calculate $f$-branch statistics (Malinsky et al., 2018). The $f$-branch metric, or $f_b(C)$, is a summary of $f_4$-ratio scores that capture excess allele sharing between a species $C$ and a branch $b$ of a given tree. In this way, the $f$-branch statistic deals with the possibility of correlated $f_4$ statistics related to a single gene flow event and also is able to infer the directionality of introgression. For $f$-branch calculations, we calculated statistics constrained to (1) the maximum likelihood topology from `RAxML` in our analyses, and (2) the species tree from Koblmüller et al. (2021). We calculated these statistics for both the GBS and WGS variant-only SNP datasets (both filtered for <50% missing data and minor allele frequency >0.01), separating the two *L. niloticus* individuals into separate "West Africa" and "Congo" lineages in the GBS dataset, based on collection locations and the recent suggestion that these two lineages of *L. niloticus* may be paraphyletic with respect to the Lake Tanganyika radiation (Koblmüller et al., 2021). We additionally calculated $f$-branch statistics for each chromosome separately for the WGS dataset constrained to the maximum likelihood topology from our `RAxML` analyses to quantify broad-scale differences in rates of introgression across the genome.

To further investigate differential patterns of admixture signal across the genome, we calculated sliding window analyses of $f_d$. The $f_d$ statistic is related to the $D$-statistic in that it tests for excess allele sharing among taxa but is designed for use in smaller regions of the genome and is used to identify regions of excess allele sharing contributing to the overall $D$-statistic observed (Martin et al., 2015). To quantify the proportion of the genome involved in admixture, we calculated $f_d$ using `Dsuite Dinvestigate` for 100-informative SNP sliding windows with 20-SNP step size, iterating over all trios involving two LT taxa as P1 and P2, with *L. niloticus* as P3 where (((P1,P2),P3),O). The $f_d$ statistic gives meaningless values if $D$ is negative, and thus $f_d$ values for windows with negative $D$ were converted to missing data, as recommended by the authors. We calculated the $z$-score for each $f_d$ value, as well as its associated $p$-value, and identified significantly admixed regions as those with $z > 2.33$. We identified continuous admixed blocks as those where sequential windows exhibited an elevated signal of admixture, and calculated the length of these blocks. We did this by using a seed-and-expand method, where blocks with $z > 2.33$ were used as seed locations, and then windows around these were merged iteratively into continuous blocks until a maximum of 3 non-significant ($z < 1.5$) blocks was reached. Admixed tracts were defined as those containing at least 5 consecutive windows due to the overlapping nature of our windowed $f_d$ estimation. We then compared the distribution of admixed blocks among species comparisons and among chromosomes, and calculated the total admixed tracts, mean admixed block size and mean proportion of the sequenced genome that exhibited a significantly elevated admixture signal for each chromosome and genome-wide, averaging across trios with the same P2 taxon.

To investigate whether those regions of the genome exhibiting elevated levels of introgression contribute disproportionately to topological uncertainty, we conducted sliding window analysis of gene trees in `TWISST2` (Martin, 2026). We used our WGS VCF (filtered to retain sites with <50% missing data, minimum minor allele frequency of 0.01, and minimum depth of 30) as input to `sticcs prep`, identifying *L. calcarifer* as our outgroup taxon. The `sticcs` program is a model-free method for inferring the series of ancestral recombination graphs (ARGs) along the genome that does not require phased haplotypes (Martin, 2026). With the output from `sticcs`, we ran `TWISST2` on this "sticcstack" using the recommended settings (i.e., `max_subtrees=512`), and then topology weightings were imported into `R` for plotting.

To compare $f_d$ windows and topology weightings to recombination rates across the genome, we estimated population recombination rates for each of the four LT species using the recombinant neural network method implemented in `ReLERNN` (Adrion et al., 2020). We first created a genome mask file using the `mop` utility available from https://github.com/RILAB/mop, using the default settings and quality thresholds (`-Q` and `-q`) of 20, and a minimum mean depth across all individuals (`-M`) of 3. We then ran the Simulate, Train, Predict, and BSCorrect modules of `ReLERNN` using the default settings except for Simulate (nTrain 100000, nVali 2000, nTest 2000) and BSCorrect (nSlice 200, nReps 1000). We set the mutation rate to $3.5x10^{-9}$, generation time to 3 years, and the upper bound for the Rho-Theta ratio to 2. When analyzing these data, we filtered out any windows with fewer than 100 sites (nSites <100) and any estimated corrected recombination rates more than two standard deviations above the overall mean ($> 9.33x10^{-9}$) as being implausibly high compared to the genome-wide average.

**Demographic modeling.** To test competing hypotheses for the evolutionary history and gene flow among our focal species, we explicitly modeled demographic history using `fastsimcoal2` (v2603, Excoffier et al., 2013). Analyses in `fastsimcoal2` use a continuous-time sequential Markovian coalescent approximation to estimate demographic parameters from the site frequency spectrum (SFS). We used the WGS dataset including all sites with genotypes for all individuals (no missing data) with read depth >30 and <1000, and generated the SFS using `easySFS` (https://github.com/isaacovercast/easySFS) with no down projection. We built both six-taxon models including the four Lake Tanganyika species, *L. niloticus*, and *L. calcarifer*, and two-taxon models for all combinations of ingroup taxa (i.e., *L. angustifrons*, *L. mariae*, *L. microlepis*, *L. stappersii*, and *L. niloticus*).

***Six taxon models.*** We tested a total of 32 six-taxon demographic models, all having the same pattern of divergence history conditioned on the topology of the best `RAxML` tree but differing from one another by the number of admixture events between the taxa (Fig. S2). For each model, `fastsimcoal2` was used to estimate the expected multi-SFS generated using 400 independent runs, each involving 200,000 coalescent simulations. We performed parameter estimation using the maximum composite likelihood from the SFS (`-M`), a minimum SFS count threshold of 10 (`-C 10`), and 50 expectation-maximization cycles (`-L 50`). For each set of 400 runs, the run that led to the highest maximum likelihood was selected as the best. In all models, we constrained the split between *L. calcarifer* and the freshwater *Lates* to 11 million generations based on the oldest known *Lates* fossil found in Africa (*L. qatraniensis* from the Jebel Qatrani Formation of the Fayum Depression in Egypt, ∼33 Mya; Murray and Attia, 2004).

We assessed model fit and performance via two approaches, following Meier et al. (2017). First, we compared each model's best run using the Akaike information criterion (AIC), adjusted to penalize parameters by a factor of 200 due to the large likelihoods generated in six-taxon models (Johri et al., 2021). Second, to determine whether models are truly different or if they differ simply due to stochasticity in the likelihood approximation, we ran each model with its best parameter values 300 times using 1,000,000 coalescent simulations. The likelihoods differ between these runs because `fastsimcoal` approximates the likelihood using simulations, rather than calculating the likelihood exactly. If the ranges of likelihoods of two models overlap, it suggests that they provide an equally good fit to the observed data. This additional method of model selection circumvents some of the problems with AIC, as AIC can overestimate support for the most likely model in situations where SNPs are not independent, and we did not filter our data explicitly for linkage.

To obtain confidence intervals around parameter estimates of the best fit model identified with AIC and the likelihood distributions, we used a non-parametric block bootstrapping approach, following Meier et al. (2017). Briefly, 100 bootstrap SNP datasets were created by randomly sampling 10,000-SNP blocks from the original dataset with replacement to form datasets of roughly equivalent size to the original, followed by conversion to an SFS using `easySFS`. For each bootstrap SFS, `fastsimcoal2` was then used to estimate the expected multi-SFS from 200 independent runs using the same settings as the original best fit model, with 100,000 coalescent simulations per run and using the best fit model's parameter estimates as starting values. The best run from each of these 100 bootstrap datasets was then selected and parameter estimates compiled to generate distributions and 95% bootstrap confidence intervals for each parameter in the model.

***Alternate fossil calibrations.*** For the best model in our six-taxon `fastsimcoal2` model set, we tested the influence of our fossil calibration on the dates inferred for divergence times between *Lates* taxa by re-running this model with an alternate fossil calibration scheme, following Koblmüller et al. (2021). For the alternate fossil calibration, we constrained the split between *L. calcarifer* and the freshwater *Lates* taxa to 5.3 million generations, based on the oldest *L. niloticus* fossil known (Koblmüller et al., 2021; Otero and Gayet, 2001), and kept all other parameterizations the same. We again used `fastsimcoal2` to estimate the expected multi-SFS generated from 400 independent runs, each consisting of 200,000 coalescent simulations per estimate, generated by a maximum of 50 expectation-maximization cycles. Parameter estimation was performed using maximum composite likelihood from the SFS (`-M`) and a minimum SFS count threshold of 10 was used for likelihood estimation (`-C 10`). From the set of 400 runs, the run that led to the highest maximum likelihood was selected as the best. For the best run, we again obtained confidence intervals around parameter estimates via a non-parametric block bootstrapping approach with 100 bootstrap replicates.

***Two taxon models.*** To further investigate the admixture events identified in our best six-taxon `fastsimcoal` model, we parameterized and competed a series of two-taxon models for each two-species pair of ingroup taxa (i.e., *L. niloticus*, *L. angustifrons*, *L. mariae*, *L. microlepis*, *L. stappersii*). These two-taxon models involved divergence between the two species, followed by no admixture events ("noadmix"), one admixture event ("1admix"), two admixture events ("2admix"), or three admixture events ("3admix"). In addition, we tested hypotheses of divergence with gene flow between taxa, where continuous gene flow occurred from divergence to some point in the past, followed by either no instantaneous admixture events, one admixture event, or two admixture events. For each of the evolutionary hypotheses involving admixture events, we had one model with unidirectional gene flow (i.e., only from one species into the other) and one model with bidirectional gene flow, where the migration rates were allowed to vary in each direction (thus allowing for bidirectional but uneven rates of gene flow). All of these models included invariant sites and the mutation rate was fixed at $3.5x10^{-9}$ to allow for absolute estimates of parameters. Files containing model specifications (i.e., `tpl` and `est` files) can be found on GitHub at http://github.com/jessicarick/lates-admixture.

We estimated the site frequency spectrum for each two-species pair from a combination of the genotyping-by-sequencing and whole genome sequencing data. To prepare these data, we used `bcftools mpileup` and `bcftools call` to generate a `VCF` containing all called sites for all individuals from all species from individual `BAM` files. We then subsetted this `VCF` for each pair of species, retaining only the sequences belonging to the two species. We then filtered the sites in each subsetted `VCF` using `vcftools`, choosing to retain those sites with a minimum quality >20, minimum read depth >10, and a maximum of 30% missing data. We used `easySFS -preview` to calculate the best number of individuals to retain for down-projection from each `VCF`, choosing the projection values that would maximize both the number of sites and the number of individuals retained.

We estimated the expected two-dimensional SFS from 300 independent runs, each consisting of 200,000 coalescent simulations per run, generated by a maximum of 50 expectation-maximization cycles. Parameter estimation was performed using maximum composite likelihood from the SFS and a minimum SFS count threshold of 10 was used for likelihood estimation. For each set of 400 runs, the run that led to the highest maximum likelihood was selected as the best. As with the six-taxon models, we used a combination of AIC and likelihood distributions (re-estimated 300 times using 100,000 coalescent simulations) to evaluate model fit and performance, selecting the best model for each species pair as that with the lowest AIC and highest median likelihood.

***Calculating generation time.*** As no reliable estimates of generation time exist for all of the species we are studying, we calculated the mean generation time using data from FishBase (Froese and Pauly, 2021) and the calculations outlined there, which are based on the von Bertalanffy Growth Function (Von Bertalanffy, 1938):

$$L_t = L_\infty(1 - e^{-K(t-t_0)}) \qquad [1]$$

Values of $K$ (the growth coefficent), $M$ (the rate of natural mortality), and $L_\infty$ (the asymptotic length) were pulled from FishBase for each species, as available. As suggested in Froese and Pauly (2021), we used $t_{opt}$, the time at which a fish reaches $L_{opt}$, as an approximation of generation time. For species without a published estimate for $M$, we calculated $M$ from $T$ (the annual mean water temperature) as $M = 10^{0.566-0.718*log(L_\infty)} + 0.02T$. The $L_{opt}$ can then be estimated from $K$ and $M$ as $L_{opt} = L_\infty(\frac{3}{3+\frac{M}{K}})$ (Beverton, 1992). The value for $t_0$ was calculated using the following equation (Pauly, 1979):

$$log(-t_0) = -0.3922 - 0.2752log(L_\infty) - 1.038log(K) \qquad [2]$$

We then calculated the mean $t_{opt}$ for the six species (mean $t_{opt} = 2.96$), and rounded this estimate to 3 for our generation time in `fastsimcoal2` and `MSMC2` analyses. The longest generation time was 3.58 (*L. niloticus*), while the shortest was 2.38 (*L. stappersii*). Calculated generation times were relatively similar to the ages at sexual maturity reported in the literature, where available (Coulter, 1991; Froese and Pauly, 2021).

***Diversity contrasts across lakes.*** Lake Malawi and Lake Tanganyika "open-water" species were identified from the habitat classification in **?** and assigned to feeding guilds from the diet information in that dataset. We included any species that had "open water" as any part of its habitat classification. For Lake Victoria, we used Witte et al. (1992) as a data source, taking all species found at trawl stations F-I (10-14 m) as representing the offshore community. We checked checked trophic categories against the guild classifications for species used in McGee et al. (2015). Species still recorded in 1987/90 by Witte et al. (1992), after the *L. niloticus* expansion, were classed as persisting. This classification is generous, since it counts a species caught in any gear or at any station. It should be noted that these diversity contrasts are imperfect, as the species data were compiled in very different ways: a single set of trawl surveys in Lake Victoria compared to full assembled species lists from Lake Malawi and Lake Victoria. However, the general patterns we can learn from this are highly illustrative.

### Supplementary Results

Genotyping-by-sequencing (GBS) resulted in short-read DNA sequence data for 150 individual *Lates* from a broad geographic area along the Tanzanian shore of Lake Tanganyika (Fig. 1), as well as three *L. niloticus* museum specimens (collected in Cameroon and from the Congo River), all of which were assembled to the chromosomal-level *Lates calcarifer* draft genome (V3, Vij et al., 2016). The Nile perch (*L. niloticus*) is widespread across the major lakes and river basins of tropical sub-Saharan Africa, and is hypothesized to be the colonizer of early LT that then diverged into the four LT-endemic *Lates*. We obtained 258,651,838 raw reads from the sequenced GBS libraries, which were filtered and assembled to the *Lates calcarifer* draft genome (Version 3; Vij et al., 2016), resulting in an average of 1.0 million reads assembled per individual and 161 individuals retained after filtering for low coverage ($<$ 20,000 reads). These individuals include 158 Lake Tanganyika *Lates* (31 *L. angustifrons*, 37 *L. mariae*, 18 *L. microlepis* and 72 *L. stappersii*), 2 *L. niloticus* from Cameroon, and 1 *L. calcarifer* from southeast Asia, whose FASTQ reads were downloaded from the NCBI Sequence Read Archive (SRA run SRR3140997). Whole genome sequencing (WGS) data were obtained for a subset of 3 individuals from each Tanganyika-endemic species and 1 *L. niloticus* from the Congo River. In addition, we included one *L. calcarifer* individual as an outgroup in phylogenetic analyses for both GBS and WGS datasets (Vij et al., 2016). The median genome coverage at sequenced sites was 14x per individual for GBS and 63.6x per individual for WGS, and assembly statistics did not differ systematically among taxa.

In contrast to the other phylogenetic reconstruction methods, `SNAPP` converged on a tree in which *L. niloticus* nested within the LT species, making the LT-endemic clade paraphyletic; however, species tree methods that do not explicitly account for admixture are known to be sensitive to and biased by admixture among species (Leaché et al., 2014). After removing putatively admixed regions, we find that the `SNAPP` phylogeny matches the topology from `RAxML`, and that we once again infer a monophyletic LT clade.

In compiling data on the comparative cichlid diversity in the pelagic of Lakes Malawi, Victoria and Tanganyika, we found that open-water cichlid assemblages differed strongly among lakes (Fig. S21). Lake Malawi, which has no *Lates*, had 28 species classified as "open-water", of which 22 were piscivores. Lake Tanganyika, with native *Lates*, had 8, of which 5 were piscivores. In Lake Victoria, the offshore assemblage before *Lates* comprised 56 taxa, 14 of them piscivores. Of these, only 11 were still caught in 1987/90, and none of the 11 was a piscivore.

### References

Adrion, J. R., Galloway, J. G., and Kern, A. D. (2020). Predicting the Landscape of Recombination Using Deep Learning. *Molecular Biology and Evolution*, 37(6):1790–1808.

Bandelt, H. J., Forster, P., and Rohl, A. (1999). Median-joining networks for inferring intraspecific phylogenies. *Molecular Biology and Evolution*, 16(1):37–48.

Beverton, R. J. (1992). Patterns of reproductive strategy parameters in some marine teleost fishes. *Journal of Fish Biology*, 41.

Bryant, D., Bouckaert, R., Felsenstein, J., Rosenberg, N. A., and Roychoudhury, A. (2012). Inferring species trees directly from biallelic genetic markers: Bypassing gene trees in a full coalescent analysis. *Molecular Biology and Evolution*, 29(8):1917–1932.

Coulter, G. W. (1991). Pelagic Fish. In Coulter, G. W., editor, *Lake Tanganyika and its Life*, chapter 6, pages 111–138. Oxford University Press.

Danecek, P., Auton, A., Abecasis, G., Albers, C. A., Banks, E., DePristo, M. A., Handsaker, R. E., Lunter, G., Marth, G. T., Sherry, S. T., McVean, G., and Durbin, R. (2011). The variant call format and VCFtools. *Bioinformatics*, 27(15):2156–2158.

Eccles, D. H. (1992). FAO Species identification sheets for fishery purposes. Field guide to the freshwater fishes of Tanzania. Technical report, Food and Agriculture Organization of the United Nations, Rome.

Excoffier, L., Dupanloup, I., Huerta-Sánchez, E., Sousa, V. C., and Foll, M. (2013). Robust Demographic Inference from Genomic and SNP Data. *PLoS Genetics*, 9(10).

Froese, R. and Pauly, D. (2021). FishBase.

Gompert, Z., Lucas, L. K., Buerkle, C. A., Forister, M. L., Fordyce, J. A., and Nice, C. C. (2014). Admixture and the organization of genetic diversity in a butterfly species complex revealed through common and rare genetic variants. *Molecular Ecology*, 23(18):4555–4573.

Green, R. E., Krause, J., Briggs, A. W., Maricic, T., Stenzel, U., Kircher, M., Patterson, N., Li, H., Zhai, W., Fritz, M. H. Y., Hansen, N. F., Durand, E. Y., Malaspinas, A. S., Jensen, J. D., Marques-Bonet, T., Alkan, C., Prufer, K., Meyer, M., Burbano, H. A., Good, J. M., Schultz, R., Aximu-Petri, A., Butthof, A., Hober, B., Hoffner, B., Siegemund, M., Weihmann, A., Nusbaum, C., Lander, E. S., Russ, C., Novod, N., Affourtit, J., Egholm, M., Verna, C., Rudan, P., Brajkovic, D., Kucan, Z., Gusic, I., Doronichev, V. B., Golovanova, L. V., Lalueza-Fox, C., de la Rasilla, M., Fortea, J., Rosas, A., Schmitz, R. W., Johnson, P. L. F., Eichler, E. E., Falush, D., Birney, E., Mullikin, J. C., Slatkin, M., Nielsen, R., Kelso, J., Lachmann, M., Reich, D., and Paabo, S. (2010). A Draft Sequence of the Neandertal Genome. *Science*, 328(5979).

Ivory, S. J., Blome, M. W., King, J. W., McGlue, M. M., Cole, J. E., and Cohen, A. S. (2016). Environmental change explains cichlid adaptive radiation at Lake Malawi over the past 1.2 million years. *Proceedings of the National Academy of Sciences of the United States of America*, 113(42):11895–11900.

Johri, P., Riall, K., Becher, H., Excoffier, L., Charlesworth, B., and Jensen, J. D. (2021). The Impact of Purifying and Background Selection on the Inference of Population History: Problems and Prospects. *Molecular Biology and Evolution*, 38(7).

Koblmüller, S., Schöggl, C. A., Lorber, C. J., Van Steenberge, M., Kmentová, N., Vanhove, M. P., and Zangl, L. (2021). African lates perches (Teleostei, Latidae, Lates): Paraphyly of Nile perch and recent colonization of Lake Tanganyika. *Molecular Phylogenetics and Evolution*, 160.

Korneliussen, T. S., Albrechtsen, A., and Nielsen, R. (2014). ANGSD: Analysis of next generation sequencing data. *BMC Bioinformatics*, 15:356.

Langmead, B. and Salzberg, S. L. (2012). Fast gapped-read alignment with Bowtie 2. *Nature Methods*, 9(4):357–359.

Leaché, A. D., Harris, R. B., Rannala, B., and Yang, Z. (2014). The Influence of Gene Flow on Species Tree Estimation: A Simulation Study. *Systematic Biology*, 63(1).

Leigh, J. W. and Bryant, D. (2015). POPART: full-feature software for haplotype network construction. *Methods in Ecology and Evolution*, 6(9):1110–1116.

Li, C., Ricardo, B.-R., Leo Smith, W., and Ortí, G. (2011). Monophyly and interrelationships of Snook and Barramundi (Centropomidae sensu Greenwood) and five new markers for fish phylogenetics. *Molecular Phylogenetics and Evolution*, 60(3):463–471.

Li, H. and Durbin, R. (2009). Fast and accurate short-read alignment with Burrows-Wheeler transform. *Bioinformatics*, 25(14):1754–1760.

Li, H., Handsaker, B., Wysoker, A., Fennell, T., Ruan, J., Homer, N., Marth, G., Abecasis, G., and Durbin, R. (2009). The Sequence Alignment/Map format and SAMtools. *Bioinformatics*, 25(16):2078–2079.

Lisiecki, L. E. and Raymo, M. E. (2005). A Pliocene-Pleistocene stack of 57 globally distributed benthic $\delta$ 18O records. *Paleoceanography*, 20(1).

Malinsky, M., Matschiner, M., and Svardal, H. (2021). Dsuite - Fast D-statistics and related admixture evidence from VCF files. *Molecular Ecology Resources*, 21(2):584–595.

Malinsky, M., Svardal, H., Tyers, A. M., Miska, E. A., Genner, M. J., Turner, G. F., and Durbin, R. (2018). Whole-genome sequences of Malawi cichlids reveal multiple radiations interconnected by gene flow. *Nature Ecology and Evolution*, 2(12):1940–1955.

Martin, M., Patterson, M., Garg, S., Fischer, S. O., Pisanti, N., Klau, G. W., Schöenhuth, A., and Marschall, T. (2016). WhatsHap: fast and accurate read-based phasing. *bioRxiv*.

Martin, S. H. (2026). A model-free method for genealogical inference without phasing and its application for topology weighting. *Genetics*, 232(1):iyaf181.

Martin, S. H., Davey, J. W., and Jiggins, C. D. (2015). Evaluating the use of ABBA-BABA statistics to locate introgressed loci. *Molecular Biology and Evolution*, 32(1):244–257.

McGee, M. D., Borstein, S. R., Neches, R. Y., Buescher, H. H., Seehausen, O., and Wainwright, P. C. (2015). A pharyngeal jaw evolutionary innovation facilitated extinction in Lake Victoria cichlids. *Science*, 350(6264).

Meier, J. I., Marques, D. A., Mwaiko, S., Wagner, C. E., Excoffier, L., and Seehausen, O. (2017). Ancient hybridization fuels rapid cichlid fish adaptive radiations. *Nature Communications*, 8(1).

Murray, A. M. and Attia, Y. S. (2004). A new species of Lates (Teleostei: Perciformes) from the Lower Oligocene of Egypt. *Journal of Vertebrate Paleontology*, 24(2):299–308.

Otero, O. and Gayet, M. (2001). Palaeoichthyofaunas from the Lower Oligocene and Miocene of the Arabian Plate: palaeoecological and palaeobiogeographical implications. *Palaeogeography, Palaeoclimatology, Palaeoecology*, 165(1-2):141–169.

Parchman, T. L., Gompert, Z., Mudge, J., Schilkey, F. D., Benkman, C. W., and Buerkle, C. A. (2012). Genome-wide association genetics of an adaptive trait in lodgepole pine. *Molecular Ecology*, 21(12):2991–3005.

Patterson, N., Moorjani, P., Luo, Y., Mallick, S., Rohland, N., Zhan, Y., Genschoreck, T., Webster, T., and Reich, D. (2012). Ancient admixture in human history. *Genetics*, 192(3):1065–1093.

Pauly, D. (1979). Gill size and temperature as governing factors in fish growth: a generalization of von Bertalanffy´s growth formula.

Rambaut, A., Drummond, A. J., Xie, D., Baele, G., and Suchard, M. A. (2018). Posterior summarization in Bayesian phylogenetics using Tracer 1.7. *Systematic Biology*, 67(5).

Rick, J. A., Junker, J., Kimirei, I. A., Sweke, E. A., Dinkel, C., Mwaiko, S., Seehausen, O., and Wagner, C. E. (2021). The genetic population structure of Lake Tanganyika's Lates 1 species flock, an endemic radiation of pelagic top predators 2 3. *bioRxiv*.

Sayyari, E. and Mirarab, S. (2016). Fast Coalescent-Based Computation of Local Branch Support from Quartet Frequencies. *Molecular Biology and Evolution*, 33(7):1654–1668.

Schiffels, S. and Durbin, R. (2014). Inferring human population size and separation history from multiple genome sequences. *Nature Genetics*, 46(8):919–25.

Scholz, C. A., Johnson, T. C., Cohen, A. S., King, J. W., Peck, J. A., Overpeck, J. T., Talbot, M. R., Brown, E. T., Kalindekafe, L., Amoako, P. Y. O., Lyons, R. P., Shanahan, T. M., Castañeda, I. S., Heil, C. W., Forman, S. L., McHargue, L. R., Beuning, K. R. M., Gomez, J., and Pierson, J. (2007). East African megadroughts between 135 and 75 thousand years ago and bearing on early-moderns human origins. *Proceedings of the National Academy of Sciences*, 104(42):16416–16421.

Shastry, V., Adams, P. E., Lindtke, D., Mandeville, E. G., Parchman, T. L., Gompert, Z., and Buerkle, C. A. (2021). Model-based genotype and ancestry estimation for potential hybrids with mixed-ploidy. *Molecular Ecology Resources*, 21(5):1434–1451.

Soraggi, S., Wiuf, C., and Albrechtsen, A. (2018). Powerful Inference with the D-Statistic on Low-Coverage Whole-Genome Data. *G3 Genes|Genomes|Genetics*, 8(2):551–566.

Stamatakis, A. (2014). RAxML version 8: A tool for phylogenetic analysis and post-analysis of large phylogenies. *Bioinformatics*, 30(9):1312–1313.

Tierney, J. E., Russell, J. M., Huang, Y., Sinninghe Damsté, J. S., Hopmans, E. C., and Cohen, A. S. (2008). Northern hemisphere controls on tropical southeast African climate during the past 60,000 years. *Science*, 322(5899):252–255.

Vij, S., Kuhl, H., Kuznetsova, I. S., Komissarov, A., Yurchenko, A. A., Van Heusden, P., Singh, S., Thevasagayam, N. M., Prakki, S. R. S., Purushothaman, K., Saju, J. M., Jiang, J., Mbandi, S. K., Jonas, M., Hin Yan Tong, A., Mwangi, S., Lau, D., Ngoh, S. Y., Liew, W. C., Shen, X., Hon, L. S., Drake, J. P., Boitano, M., Hall, R., Chin, C. S., Lachumanan, R., Korlach, J., Trifonov, V., Kabilov, M., Tupikin, A., Green, D., Moxon, S., Garvin, T., Sedlazeck, F. J., Vurture, G. W., Gopalapillai, G., Kumar Katneni, V., Noble, T. H., Scaria, V., Sivasubbu, S., Jerry, D. R., O'Brien, S. J., Schatz, M. C., Dalmay, T., Turner, S. W., Lok, S., Christoffels, A., and Orbán, L. (2016). Chromosomal-Level Assembly of the Asian Seabass Genome Using Long Sequence Reads and Multi-layered Scaffolding. *PLoS Genetics*, 12(4):1–35.

Von Bertalanffy, L. (1938). A quantitative theory of organic growth (inquiries on growth laws. II). *Human Biology*, 10(2).

Witte, F., Goldschmidt, T., Wanink, J., Van Oijen, M., Goudswaard, K., Witte-Maas', E., and Bouton', N. (1992). The destruction of an endemic species flock : quantitative data on the decline of the haplochromine cichlids of Lake Victoria. Technical report.

Zhang, C., Rabiee, M., Sayyari, E., and Mirarab, S. (2018). ASTRAL-III: Polynomial time species tree reconstruction from partially resolved gene trees. *BMC Bioinformatics*, 19(Suppl 6):15–30.

### Supplementary Tables and Figures

**Table S1.** Metadata for individuals included in analyses of whole genome resequencing data. [See separate file: Supplementary_Table_1_WGS_Metadata.xlsx].

**Table S2.** Whole genome $D$-statistics for all possible three-taxon combinations, using *Lates calcarifer* as the outgroup. Values were calculated in ANGSD (Korneliussen et al., 2014; Soraggi et al., 2018) using the whole genome sequencing data and are sorted from highest to lowest $z$-score, where a positive $D$ and $z$-score indicate excess allele sharing P2 $<–>$ P3 and a negative $D$ and $z$-score indicate excess allele sharing P1 $<–>$ P3. Columns indicate the three taxa in the comparison (P1, P2, P3), the outgroup taxon (P4), the $D$-statistic (D), the block jackknife estimate of $D$ (JK-D), the variance of the block jackknife estimate of $D$ [V(JK-D)], the $z$-score associated with $D$ (Z), the $p$-value associated with the z-score (pvalue), the number of ABBA sites (nABBA), the number of BABA sites (nBABA), and the number of blocks used in the calculation of $D$ for each comparison. The following taxon abbreviations are used in the table: Lmar (*L. mariae*), Lmic (*L. microlepis*), Lsta (*L. stappersii*), Lcal (*L. calcarifer*), Lnil (*L. niloticus*), Lang (*L. angustifrons*).

| P1 | P2 | P3 | P4 | D | JK-D | V(JK-D) | Z | pvalue | nABBA | nBABA | nBlocks |
|---|---|---|---|---|---|---|---|---|---|---|---|
| Lmar | Lmic | Lsta | Lcal | 0.0866 | 0.0866 | 0.000002 | 70.09 | 0 | 394819.771 | 331908.969 | 5878 |
| Lmar | Lmic | Lnil | Lcal | 0.0425 | 0.0425 | 0.000001 | 41.35 | 0 | 311563.154 | 286149.783 | 5878 |
| Lmar | Lsta | Lmic | Lcal | 0.0241 | 0.0241 | 0.000002 | 17.75 | 0 | 394819.771 | 376244.369 | 5878 |
| Lang | Lmic | Lsta | Lcal | 0.0057 | 0.0057 | 0.000002 | 4.29 | 0.000018 | 340242.189 | 336390.058 | 5878 |
| Lmar | Lsta | Lnil | Lcal | 0.0031 | 0.0031 | 0.000002 | 2.49 | 0.0129 | 314364.87 | 312397.34 | 5878 |
| Lmar | Lmic | Lang | Lcal | -0.0170 | -0.0170 | 0.000001 | -14.74 | 0 | 375933.156 | 388953.499 | 5878 |
| Lmic | Lsta | Lnil | Lcal | -0.0375 | -0.0375 | 0.000001 | -31.31 | 0 | 292217.736 | 314959.675 | 5878 |
| Lang | Lsta | Lmic | Lcal | -0.0751 | -0.0751 | 0.000002 | -49.41 | 0 | 340242.189 | 395463.921 | 5878 |
| Lmic | Lsta | Lmar | Lcal | -0.0626 | -0.0626 | 0.000001 | -54.34 | 0 | 331908.969 | 376244.369 | 5878 |
| Lmic | Lsta | Lang | Lcal | -0.0807 | -0.0807 | 0.000001 | -67.13 | 0 | 336390.058 | 395463.921 | 5878 |
| Lmar | Lsta | Lang | Lcal | -0.0927 | -0.0927 | 0.000002 | -74.54 | 0 | 352858.195 | 424974.049 | 5878 |
| Lang | Lmic | Lnil | Lcal | -0.0851 | -0.0851 | 0.000001 | -76.00 | 0 | 254073.084 | 301341.484 | 5878 |
| Lang | Lmar | Lsta | Lcal | -0.0910 | -0.0910 | 0.000001 | -84.06 | 0 | 294003.421 | 352858.195 | 5878 |
| Lang | Lsta | Lnil | Lcal | -0.1153 | -0.1153 | 0.000002 | -88.76 | 0 | 258719.328 | 326142.244 | 5878 |
| Lang | Lmar | Lmic | Lcal | -0.1136 | -0.1136 | 0.000001 | -101.93 | 0 | 299249.637 | 375933.156 | 5878 |
| Lang | Lmic | Lmar | Lcal | -0.1303 | -0.1303 | 0.000001 | -109.69 | 0 | 299249.637 | 388953.499 | 5878 |
| Lang | Lmar | Lnil | Lcal | -0.1295 | -0.1295 | 0.000001 | -121.26 | 0 | 242830.608 | 315083.583 | 5877 |
| Lang | Lsta | Lmar | Lcal | -0.1822 | -0.1822 | 0.000002 | -146.06 | 0 | 294003.421 | 424974.049 | 5878 |
| Lmar | Lnil | Lsta | Lcal | -0.3811 | -0.3811 | 0.000004 | -198.70 | 0 | 314364.87 | 701597.593 | 5878 |
| Lsta | Lnil | Lang | Lcal | -0.3657 | -0.3657 | 0.000003 | -214.64 | 0 | 326142.244 | 702140.923 | 5878 |
| Lmar | Lnil | Lmic | Lcal | -0.3967 | -0.3967 | 0.000003 | -221.53 | 0 | 311563.154 | 721216.779 | 5878 |
| Lmar | Lnil | Lang | Lcal | -0.4165 | -0.4165 | 0.000003 | -230.76 | 0 | 315083.583 | 764835.701 | 5878 |
| Lsta | Lnil | Lmic | Lcal | -0.4036 | -0.4036 | 0.000003 | -233.02 | 0 | 314959.675 | 741204.149 | 5878 |
| Lsta | Lnil | Lmar | Lcal | -0.3838 | -0.3838 | 0.000003 | -233.20 | 0 | 312397.34 | 701597.593 | 5878 |
| Lmic | Lnil | Lsta | Lcal | -0.4345 | -0.4345 | 0.000003 | -237.38 | 0 | 292217.736 | 741204.149 | 5878 |
| Lmic | Lnil | Lang | Lcal | -0.4199 | -0.4199 | 0.000003 | -247.47 | 0 | 301341.484 | 737583.101 | 5878 |
| Lang | Lnil | Lsta | Lcal | -0.4615 | -0.4615 | 0.000003 | -248.25 | 0 | 258719.328 | 702140.923 | 5878 |
| Lmic | Lnil | Lmar | Lcal | -0.4319 | -0.4319 | 0.000003 | -262.79 | 0 | 286149.783 | 721216.779 | 5878 |
| Lang | Lnil | Lmic | Lcal | -0.4876 | -0.4876 | 0.000003 | -278.25 | 0 | 254073.084 | 737583.101 | 5878 |
| Lang | Lnil | Lmar | Lcal | -0.5180 | -0.5180 | 0.000003 | -308.56 | 0 | 242830.608 | 764835.701 | 5878 |

**Table S3.** $D$-statistics for the five-taxon *Lates* phylogeny, using *L. calcarifer* as the outgroup, with $D$-statistic comparisons constrained to the species tree topology from `RAxML` [i.e., comparisons were arrange so that ((P1,P2),P3) arrangements match the species tree]. Values were calculated using `Dsuite` (Malinsky et al., 2021) using the whole genome sequencing data. Columns indicate the three taxa in the comparison (P1, P2, P3), the $D$-statistic (D), the $z$-score associated with $D$ (Z-score), the $p$-value associated with the $z$-score (p-value), the estimated admixture fraction (f4-ratio), the number of concordant BBAA sites (BBAA), the number of ABBA sites (ABBA), and the number of BABA sites (BABA). In addition, the $p$-values from two different clustering tests are shown; a significant value for clustering provides evidence in support of the clustering of ABBA sites along the genome, which is expected when the signal is related to true introgression versus a false positive signal driven by substitution rate variation along the genome [see discussion in ?]. The following taxon abbreviations are used in the table: Lmar (*L. mariae*), Lang (*L. angustifrons*), Lmic (*L. microlepis*), Lnil (*L. niloticus*), Lsta (*L. stappersii*).

| P1 | P2 | P3 | D | Z-score | p-value | f4-ratio | Clustering (Sensitive) | Clustering (Robust) | BBAA | ABBA | BABA |
|---|---|---|---|---|---|---|---|---|---|---|---|
| Lmar | Lang | Lmic | 0.0247 | 18.90 | 2.3e-16 | 0.0241 | 0.00651 | 0.167 | 288504 | 253501 | 241278 |
| Lmar | Lang | Lnil | 0.0170 | 17.93 | 2.3e-16 | 0.00455 | 8.62e-07 | 0.238 | 624773 | 225704 | 218143 |
| Lang | Lmar | Lsta | 0.00720 | 6.10 | 1.05936e-09 | 0.00369 | 0.000252 | 0.380 | 327971 | 243590 | 240105 |
| Lmic | Lang | Lnil | 0.0273 | 28.93 | 2.3e-16 | 0.00741 | 0.000101 | 0.244 | 596404 | 232337 | 219971 |
| Lsta | Lmic | Lang | 0.0975 | 70.61 | 2.3e-16 | 0.1060 | 2.58e-13 | 1.32e-05 | 278461 | 297622 | 244759 |
| Lsta | Lang | Lnil | 0.0959 | 78.35 | 2.3e-16 | 0.0276 | 4.83e-11 | 0.632 | 580004 | 268800 | 221774 |
| Lmic | Lmar | Lnil | 0.0105 | 11.72 | 2.3e-16 | 0.0029 | 4.70e-06 | 0.654 | 591248 | 231844 | 227039 |
| Lsta | Lmic | Lmar | 0.0686 | 55.70 | 2.3e-16 | 0.0816 | 6.96e-09 | 0.013 | 282629 | 289566 | 252412 |
| Lsta | Lmar | Lnil | 0.0799 | 68.16 | 2.3e-16 | 0.0232 | 2.24e-10 | 0.293 | 588899 | 266650 | 227184 |
| Lsta | Lmic | Lnil | 0.0706 | 63.14 | 2.3e-16 | 0.0203 | 8.98e-10 | 0.0222 | 620019 | 262747 | 228087 |

**Table S4.** Summary of significant $f_d$ windows for each trio tested in `Dsuite Dinvestigate`. Windows were defined as significant when the $z$-score was >2.33, and combined with neighboring windows that had $z > 1.5$, allowing for up to 3 non-significant windows per block. For each trio and each chromosome, we calculated the average length of these continuous blocks, the total length of the blocks, and the average $f_d$ among significant windows. Trio names match those in Fig. S10.

| Trio | Chromosome | Mean window length (bp) | Total window length (bp) | Mean $f_d$ |
|---|---|---|---|---|
| Lmar_Lang_Lnil (P1 = *L. mariae*, P2 = *L. angustifrons*, P3 = *L. niloticus*) | | | | |
| | ASB_LG1 | 36737 | 514320 | 0.167 |
| | ASB_LG10 | 42167 | 126501 | 0.144 |
| | ASB_LG11 | 14810 | 148096 | 0.178 |
| | ASB_LG12 | 27019 | 189132 | 0.193 |
| | ASB_LG13 | 26635 | 266348 | 0.216 |
| | ASB_LG14 | 38240 | 458886 | 0.229 |
| | ASB_LG15 | 26401 | 264011 | 0.176 |
| | ASB_LG16_LG22 | 22102 | 66305 | 0.213 |
| | ASB_LG17 | 33640 | 168201 | 0.159 |
| | ASB_LG18 | 29146 | 58293 | 0.152 |
| | ASB_LG19 | 19496 | 77983 | 0.122 |
| | ASB_LG2 | 13269 | 13269 | 0.120 |
| | ASB_LG20 | 11010 | 33029 | 0.148 |
| | ASB_LG21 | 33142 | 66284 | 0.121 |
| | ASB_LG23 | 45207 | 135620 | 0.267 |
| | ASB_LG24 | 39760 | 79521 | 0.118 |
| | ASB_LG3 | 23286 | 349291 | 0.173 |
| | ASB_LG4 | 38672 | 154688 | 0.139 |
| | ASB_LG5 | 37908 | 379081 | 0.138 |
| | ASB_LG6 | 9117 | 45586 | 0.138 |
| | ASB_LG7_1 | 33894 | 203361 | 0.168 |
| | ASB_LG7_2 | 32183 | 128731 | 0.155 |
| | ASB_LG8 | 18189 | 163705 | 0.264 |
| | ASB_LG9 | 14472 | 72360 | 0.142 |
| Lmic_Lang_Lnil (P1 = *L. mariae*, P2 = *L. microlepis*, P3 = *L. niloticus*) | | | | |
| | ASB_LG1 | 42944 | 472383 | 0.164 |
| | ASB_LG10 | 23030 | 92118 | 0.155 |
| | ASB_LG11 | 26803 | 402045 | 0.196 |
| | ASB_LG12 | 20878 | 146145 | 0.171 |
| | ASB_LG13 | 16501 | 181516 | 0.171 |
| | ASB_LG14 | 32133 | 481994 | 0.214 |
| | ASB_LG15 | 32050 | 288453 | 0.182 |
| | ASB_LG16_LG22 | 15134 | 105935 | 0.168 |
| | ASB_LG17 | 38594 | 308752 | 0.146 |
| | ASB_LG18 | 52274 | 52274 | 0.168 |
| | ASB_LG19 | 12698 | 25397 | 0.136 |
| | ASB_LG2 | 31062 | 124249 | 0.228 |
| | ASB_LG20 | 36117 | 72234 | 0.231 |
| | ASB_LG21 | 6779 | 6779 | 0.225 |
| | ASB_LG23 | 55783 | 167348 | 0.182 |
| | ASB_LG24 | 34907 | 69814 | 0.176 |
| | ASB_LG3 | 27744 | 499386 | 0.156 |
| | ASB_LG4 | 33268 | 332679 | 0.142 |
| | ASB_LG5 | 31079 | 341864 | 0.194 |
| | ASB_LG6 | 35893 | 215360 | 0.167 |
| | ASB_LG7_1 | 21918 | 109592 | 0.167 |
| | ASB_LG7_2 | 28176 | 140878 | 0.181 |
| | ASB_LG8 | 38032 | 228193 | 0.286 |
| | ASB_LG9 | 25773 | 128867 | 0.164 |
| Lsta_Lang_Lnil (P1 = *L. stappersii*, P2 = *L. angustifrons*, P3 = *L. niloticus*) | | | | |
| | ASB_LG1 | 42131 | 589830 | 0.200 |
| | ASB_LG10 | 27378 | 82134 | 0.206 |
| | ASB_LG11 | 26556 | 398334 | 0.225 |
| | ASB_LG12 | 18572 | 74286 | 0.193 |
| | ASB_LG13 | 26136 | 313626 | 0.212 |
| | ASB_LG14 | 37513 | 675230 | 0.238 |
| | ASB_LG15 | 29348 | 410871 | 0.165 |

| | | | |
|---|---|---|---|
| ASB_LG16_LG22 | 27098 | 243886 | 0.176 |
| ASB_LG17 | 37201 | 334807 | 0.169 |
| ASB_LG18 | 30464 | 60928 | 0.149 |
| ASB_LG19 | 30352 | 364229 | 0.182 |
| ASB_LG2 | 25721 | 102885 | 0.268 |
| ASB_LG20 | 32148 | 192885 | 0.189 |
| ASB_LG21 | 22928 | 183422 | 0.163 |
| ASB_LG23 | 46676 | 140027 | 0.182 |
| ASB_LG24 | 33384 | 133536 | 0.158 |
| ASB_LG3 | 32311 | 743157 | 0.182 |
| ASB_LG4 | 40872 | 490460 | 0.189 |
| ASB_LG5 | 33077 | 562313 | 0.211 |
| ASB_LG6 | 21275 | 212752 | 0.174 |
| ASB_LG7_1 | 33189 | 265513 | 0.188 |
| ASB_LG7_2 | 31279 | 406632 | 0.198 |
| ASB_LG8 | 41114 | 287801 | 0.238 |
| ASB_LG9 | 57316 | 458524 | 0.194 |
| Lang_Lmar_Lnil (P1 = *L. angustifrons*, P2 = *L. mariae*, P3 = *L. niloticus*) | | | |
| ASB_LG1 | 21510 | 21510 | NaN |
| ASB_LG10 | 42084 | 84168 | 0.295 |
| ASB_LG11 | 32944 | 329438 | 0.289 |
| ASB_LG12 | 28528 | 114113 | 0.155 |
| ASB_LG13 | 18286 | 146289 | 0.192 |
| ASB_LG14 | 38842 | 427264 | 0.203 |
| ASB_LG15 | 21402 | 149817 | 0.153 |
| ASB_LG16_LG22 | 29032 | 58065 | 0.157 |
| ASB_LG17 | 23498 | 70493 | 0.253 |
| ASB_LG19 | 13190 | 65951 | 0.190 |
| ASB_LG2 | 26514 | 26514 | 0.133 |
| ASB_LG20 | 41462 | 82925 | 0.196 |
| ASB_LG21 | 19836 | 99182 | 0.249 |
| ASB_LG24 | 21322 | 85286 | 0.258 |
| ASB_LG3 | 42637 | 426372 | 0.160 |
| ASB_LG4 | 34024 | 68049 | 0.203 |
| ASB_LG5 | 19682 | 98408 | 0.153 |
| ASB_LG6 | 27109 | 108436 | 0.172 |
| ASB_LG7_1 | 13420 | 93943 | 0.166 |
| ASB_LG7_2 | 14493 | 57972 | 0.124 |
| ASB_LG8 | 21491 | 128946 | 0.191 |
| ASB_LG9 | 14498 | 28996 | 0.229 |
| Lmic_Lmar_Lnil (P1 = *L. microlepis*, P2 = *L. mariae*, P3 = *L. niloticus*) | | | |
| ASB_LG1 | 48572 | 291435 | 0.158 |
| ASB_LG10 | 29903 | 89710 | 0.143 |
| ASB_LG11 | 37210 | 372105 | 0.265 |
| ASB_LG12 | 49834 | 199335 | 0.184 |
| ASB_LG13 | 12263 | 122633 | 0.198 |
| ASB_LG14 | 26057 | 364800 | 0.214 |
| ASB_LG15 | 23480 | 258285 | 0.169 |
| ASB_LG16_LG22 | 17416 | 69665 | 0.154 |
| ASB_LG17 | 27208 | 163250 | 0.165 |
| ASB_LG19 | 12938 | 77626 | 0.116 |
| ASB_LG2 | 32131 | 128523 | 0.206 |
| ASB_LG20 | 32461 | 64922 | 0.248 |
| ASB_LG21 | 19506 | 117037 | 0.214 |
| ASB_LG23 | 48649 | 145947 | 0.152 |
| ASB_LG24 | 27460 | 137299 | 0.189 |
| ASB_LG3 | 29242 | 409394 | 0.142 |
| ASB_LG4 | 21392 | 171138 | 0.143 |
| ASB_LG5 | 30398 | 243183 | 0.187 |
| ASB_LG6 | 38085 | 304679 | 0.142 |
| ASB_LG7_1 | 18317 | 128219 | 0.147 |
| ASB_LG7_2 | 28463 | 142317 | 0.162 |
| ASB_LG8 | 40791 | 326331 | 0.259 |
| ASB_LG9 | 17761 | 106566 | 0.182 |
| Lsta_Lmar_Lnil (P1 = *L. stappersii*, P2 = *L. mariae*, P3 = *L. niloticus*) | | | |
| ASB_LG1 | 39248 | 470973 | 0.161 |

| | | | |
|---|---|---|---|
| ASB_LG10 | 27116 | 189815 | 0.154 |
| ASB_LG11 | 34664 | 554631 | 0.191 |
| ASB_LG12 | 56984 | 284919 | 0.163 |
| ASB_LG13 | 20239 | 303588 | 0.191 |
| ASB_LG14 | 34171 | 785927 | 0.214 |
| ASB_LG15 | 22803 | 296445 | 0.163 |
| ASB_LG16_LG22 | 36323 | 217937 | 0.172 |
| ASB_LG17 | 28453 | 227621 | 0.139 |
| ASB_LG18 | 21962 | 21962 | 0.128 |
| ASB_LG19 | 39152 | 352369 | 0.182 |
| ASB_LG2 | 22350 | 111750 | 0.225 |
| ASB_LG20 | 32902 | 230316 | 0.165 |
| ASB_LG21 | 21798 | 217982 | 0.210 |
| ASB_LG23 | 39098 | 78197 | 0.233 |
| ASB_LG24 | 39213 | 235280 | 0.166 |
| ASB_LG3 | 34835 | 766379 | 0.179 |
| ASB_LG4 | 33655 | 538479 | 0.163 |
| ASB_LG5 | 38885 | 427735 | 0.188 |
| ASB_LG6 | 23048 | 207436 | 0.152 |
| ASB_LG7_1 | 21688 | 238567 | 0.169 |
| ASB_LG7_2 | 28715 | 373299 | 0.174 |
| ASB_LG8 | 27766 | 333195 | 0.211 |
| ASB_LG9 | 44565 | 445649 | 0.173 |
| Lang_Lmic_Lnil (P1 = *L. angustifrons*, P2 = *L. microlepis*, P3 = *L. niloticus*) | | | |
| ASB_LG1 | 12948 | 38845 | 0.223 |
| ASB_LG10 | 27084 | 162501 | 0.226 |
| ASB_LG11 | 21289 | 212886 | 0.256 |
| ASB_LG13 | 14636 | 102454 | 0.213 |
| ASB_LG14 | 43092 | 344738 | 0.185 |
| ASB_LG15 | 12795 | 63976 | 0.199 |
| ASB_LG17 | 27839 | 55678 | 0.392 |
| ASB_LG19 | 13504 | 94528 | 0.183 |
| ASB_LG2 | 46876 | 93753 | 0.310 |
| ASB_LG20 | 43769 | 87538 | 0.245 |
| ASB_LG21 | 13627 | 54507 | 0.318 |
| ASB_LG24 | 24073 | 24073 | 0.273 |
| ASB_LG3 | 25091 | 225815 | 0.201 |
| ASB_LG4 | 14324 | 85941 | 0.170 |
| ASB_LG5 | 28062 | 84186 | 0.253 |
| ASB_LG6 | 62898 | 62898 | NaN |
| ASB_LG7_1 | 18731 | 74925 | 0.212 |
| ASB_LG7_2 | 8688 | 26064 | 0.146 |
| ASB_LG8 | 19837 | 99183 | 0.204 |
| ASB_LG9 | 20845 | 41690 | 0.146 |
| Lmar_Lmic_Lnil (P1 = *L. mariae*, P2 = *L. microlepis*, P3 = *L. niloticus*) | | | |
| ASB_LG1 | 25895 | 181266 | 0.222 |
| ASB_LG10 | 19879 | 99397 | 0.163 |
| ASB_LG11 | 17452 | 174522 | 0.230 |
| ASB_LG12 | 14168 | 85007 | 0.163 |
| ASB_LG13 | 20741 | 165929 | 0.190 |
| ASB_LG14 | 25404 | 203236 | 0.217 |
| ASB_LG15 | 15376 | 107632 | 0.201 |
| ASB_LG16_LG22 | 10314 | 30943 | 0.164 |
| ASB_LG17 | 38408 | 115225 | 0.227 |
| ASB_LG19 | 18679 | 112072 | 0.180 |
| ASB_LG2 | 25097 | 50194 | 0.303 |
| ASB_LG21 | 14033 | 28066 | 0.170 |
| ASB_LG23 | 50820 | 50820 | 0.288 |
| ASB_LG24 | 29445 | 29445 | 0.130 |
| ASB_LG3 | 15614 | 140528 | 0.192 |
| ASB_LG4 | 27294 | 109177 | 0.168 |
| ASB_LG5 | 24477 | 171340 | 0.236 |
| ASB_LG6 | 26341 | 79022 | 0.248 |
| ASB_LG7_1 | 21227 | 148588 | 0.185 |
| ASB_LG7_2 | 21326 | 42651 | 0.109 |
| ASB_LG8 | 19904 | 199044 | 0.192 |

| | | | |
|---|---|---|---|
| ASB_LG9 | 13541 | 54165 | 0.131 |
| Lsta_Lmic_Lnil (P1 = *L. stappersii*, P2 = *L. microlepis*, P3 = *L. niloticus*) | | | |
| ASB_LG1 | 25301 | 404818 | 0.175 |
| ASB_LG10 | 20989 | 146923 | 0.172 |
| ASB_LG11 | 34671 | 624076 | 0.190 |
| ASB_LG12 | 26323 | 210584 | 0.150 |
| ASB_LG13 | 24629 | 369439 | 0.169 |
| ASB_LG14 | 28549 | 456778 | 0.223 |
| ASB_LG15 | 19503 | 214536 | 0.167 |
| ASB_LG16_LG22 | 24718 | 247181 | 0.159 |
| ASB_LG17 | 28705 | 200936 | 0.136 |
| ASB_LG18 | 14112 | 14112 | 0.138 |
| ASB_LG19 | 25471 | 305656 | 0.170 |
| ASB_LG2 | 14961 | 89766 | 0.217 |
| ASB_LG20 | 32331 | 129324 | 0.158 |
| ASB_LG21 | 25376 | 279140 | 0.154 |
| ASB_LG23 | 62726 | 125452 | 0.165 |
| ASB_LG24 | 40493 | 202465 | 0.145 |
| ASB_LG3 | 31644 | 632884 | 0.182 |
| ASB_LG4 | 34054 | 442708 | 0.177 |
| ASB_LG5 | 31646 | 537988 | 0.187 |
| ASB_LG6 | 22932 | 160524 | 0.170 |
| ASB_LG7_1 | 25867 | 388004 | 0.160 |
| ASB_LG7_2 | 33390 | 333899 | 0.170 |
| ASB_LG8 | 14530 | 87177 | 0.176 |
| ASB_LG9 | 29304 | 234428 | 0.165 |
| Lmar_Lsta_Lnil (P1 = *L. mariae*, P2 = *L. stappersii*, P3 = *L. niloticus*) | | | |
| ASB_LG1 | 13831 | 55324 | 0.197 |
| ASB_LG10 | 12325 | 36975 | 0.193 |
| ASB_LG11 | 28440 | 142200 | 0.236 |
| ASB_LG12 | 32900 | 32900 | 0.281 |
| ASB_LG13 | 14438 | 43313 | 0.228 |
| ASB_LG14 | 20237 | 121422 | 0.226 |
| ASB_LG15 | 10346 | 62073 | 0.257 |
| ASB_LG16_LG22 | 8990 | 17980 | 0.300 |
| ASB_LG17 | 24091 | 48182 | 0.418 |
| ASB_LG19 | 14377 | 43131 | 0.219 |
| ASB_LG2 | 31313 | 31313 | 0.391 |
| ASB_LG21 | 12578 | 12578 | 0.149 |
| ASB_LG23 | 59000 | 59000 | 0.758 |
| ASB_LG3 | 8527 | 51161 | 0.223 |
| ASB_LG4 | 14827 | 29654 | 0.226 |
| ASB_LG5 | 23215 | 116077 | 0.249 |
| ASB_LG6 | 14185 | 14185 | 0.355 |
| ASB_LG7_1 | 19605 | 19605 | 0.375 |
| ASB_LG8 | 14992 | 89952 | 0.296 |
| ASB_LG9 | 20973 | 20973 | 0.237 |
| Lmic_Lsta_Lnil (P1 = *L. microlepis*, P2 = *L. stappersii*, P3 = *L. niloticus*) | | | |
| ASB_LG1 | 54944 | 109888 | 0.157 |
| ASB_LG11 | 34333 | 205997 | 0.277 |
| ASB_LG12 | 19417 | 58250 | 0.216 |
| ASB_LG13 | 20782 | 83129 | 0.242 |
| ASB_LG14 | 28666 | 114665 | 0.341 |
| ASB_LG15 | 17588 | 105529 | 0.209 |
| ASB_LG16_LG22 | 7666 | 15331 | 0.308 |
| ASB_LG17 | 64122 | 64122 | 0.190 |
| ASB_LG2 | 24323 | 24323 | 0.160 |
| ASB_LG21 | 13620 | 13620 | 0.158 |
| ASB_LG23 | 53783 | 53783 | 0.741 |
| ASB_LG3 | 5014 | 5014 | 0.228 |
| ASB_LG4 | 11891 | 35674 | 0.231 |
| ASB_LG5 | 25180 | 75541 | 0.291 |
| ASB_LG6 | 9342 | 18685 | 0.218 |
| ASB_LG7_1 | 15050 | 30099 | 0.207 |
| ASB_LG7_2 | 18617 | 37234 | 0.118 |
| ASB_LG8 | 9244 | 36977 | 0.294 |

| | | | |
|---|---|---|---|
| ASB_LG9 | 31424 | 62848 | 0.208 |

**Table S5.** Average divergence time estimates from cross-coalescence rates estimated using `MSMC2`. Reported divergence times are the average left and right time boundaries for the segment where the relative cross-coalescence rate for pairwise comparisons between individuals of the given species reached 50%. Estimates have been converted to years using a mutation rate of $3.5x10^{-9}$ and generation time of three years.

| Species Pair | Left Time Boundary (yr) | Right Time Boundary (yr) |
|---|---|---|
| **LT-LT** | **901,772** | **1,076,889** |
| Lang-Lmar | 636,979 | 757,300 |
| Lang-Lmic | 641,040 | 762,628 |
| Lmic-Lmar | 707,685 | 841,615 |
| Lang-Lsta | 1,122,649 | 1,334,336 |
| Lmic-Lsta | 1,146,387 | 1,391,770 |
| Lmar-Lsta | 1,155,890 | 1,373,685 |
| **Lnil-LT** | **1,108,439** | **1,337,488** |
| Lnil-Lang | 978,686 | 1,200,767 |
| Lnil-Lmar | 988,401 | 1,210,916 |
| Lnil-Lmic | 1,080,626 | 1,287,231 |
| Lnil-Lsta | 1,386,043 | 1,651,037 |

**Table S6.** Parameter estimates from nonparametric bootstrapping of the best model in `fastsimcoal2`, with parameters as indicated on model diagram to the right. Estimates are generated from the maximum likelihood estimates from 200 bootstrap datasets. Estimates of times are in units of generations, not years, and population sizes are in units of $2N$. The average generation time for this group of taxa is three years. Migration rate parameters are labeled with the two populations involved [i.e., MIG12 is admixture between (1) *L. niloticus* and (2) *L. mariae*, and MIG21 is the migration rate in the opposite direction]. Timing for admixture events matches the same labeling pattern (i.e., TADM12 is the timing of admixture events MIG12 and MIG21).

| Parameter | Mean | Median | 2.5% Quantile | 97.5% Quantile |
|---|---|---|---|---|
| **Migration Rates** | | | | |
| MIG12 | 8.240000e-04 | 1.850000e-05 | 1.000000e-07 | 6.594000e-03 |
| MIG14 | 3.617000e-04 | 1.810000e-05 | 1.000000e-07 | 1.911600e-03 |
| MIG15 | 6.928000e-04 | 1.450000e-05 | 1.000000e-07 | 4.555400e-03 |
| MIG21 | 1.713380e-02 | 1.206000e-04 | 1.000000e-07 | 1.177663e-01 |
| MIG23 | 1.325300e-03 | 8.140000e-05 | 2.000000e-07 | 1.153420e-02 |
| MIG25 | 1.474700e-03 | 2.120000e-05 | 0.000000e+00 | 1.547330e-02 |
| MIG32 | 7.788000e-04 | 4.330000e-05 | 1.000000e-07 | 6.560200e-03 |
| MIG34 | 1.217200e-03 | 1.870000e-05 | 1.000000e-07 | 8.497000e-03 |
| MIG41 | 5.438450e-02 | 2.569500e-02 | 1.000000e-07 | 1.444328e-01 |
| MIG43 | 9.051000e-04 | 7.570000e-05 | 1.000000e-07 | 8.156200e-03 |
| MIG51 | 4.599470e-02 | 6.652300e-03 | 4.000000e-07 | 1.391530e-01 |
| MIG52 | 9.979000e-04 | 3.680000e-05 | 1.000000e-07 | 7.642800e-03 |
| **Effective Population Sizes** | | | | |
| POP0 | 3.137472e+05 | 3.117270e+05 | 2.131975e+05 | 4.143505e+05 |
| POP1 | 9.084603e+04 | 8.352500e+04 | 4.543850e+04 | 1.700480e+05 |
| POP2 | 2.851361e+05 | 2.850700e+05 | 2.066840e+05 | 3.595100e+05 |
| POP3 | 3.340721e+05 | 3.270370e+05 | 2.453715e+05 | 4.589210e+05 |
| POP4 | 2.540657e+05 | 2.523180e+05 | 1.405875e+05 | 3.713435e+05 |
| POP5 | 3.364123e+05 | 3.204830e+05 | 2.086800e+05 | 5.199625e+05 |
| **Admixture Events (generations ago)** | | | | |
| TADM12 | 9.104819e+04 | 8.901600e+04 | 6.654500e+03 | 1.870345e+05 |
| TADM13 | 1.078896e+06 | 1.083528e+06 | 3.071305e+05 | 1.709435e+06 |
| TADM14 | 4.646818e+04 | 3.167300e+04 | 2.331000e+03 | 1.205125e+05 |
| TADM15 | 5.638590e+04 | 4.407900e+04 | 4.141500e+03 | 1.444000e+05 |
| TADM23 | 8.563990e+04 | 8.441200e+04 | 2.111250e+04 | 1.502070e+05 |
| TADM25 | 8.713465e+04 | 8.628600e+04 | 1.895400e+04 | 1.524090e+05 |
| **Divergence Events (generations ago)** | | | | |
| TADM34 | 8.035712e+04 | 8.208100e+04 | 1.492550e+04 | 1.564315e+05 |
| TDIVLT1LT2 | 1.943398e+05 | 1.899580e+05 | 1.532045e+05 | 2.451165e+05 |
| TDIVLT3LT4 | 1.468291e+05 | 1.449590e+05 | 8.805700e+04 | 1.924345e+05 |
| TDIVLTLT | 2.309192e+05 | 2.321800e+05 | 1.819790e+05 | 2.984175e+05 |
| TDIVNILLT | 4.603517e+05 | 4.615850e+05 | 3.248185e+05 | 6.015390e+05 |

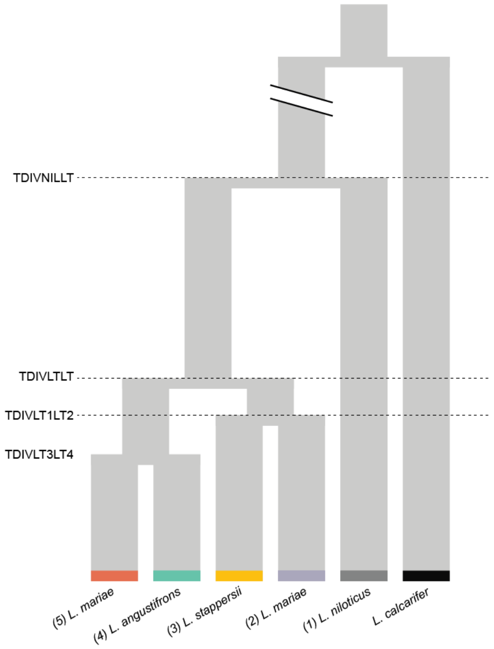

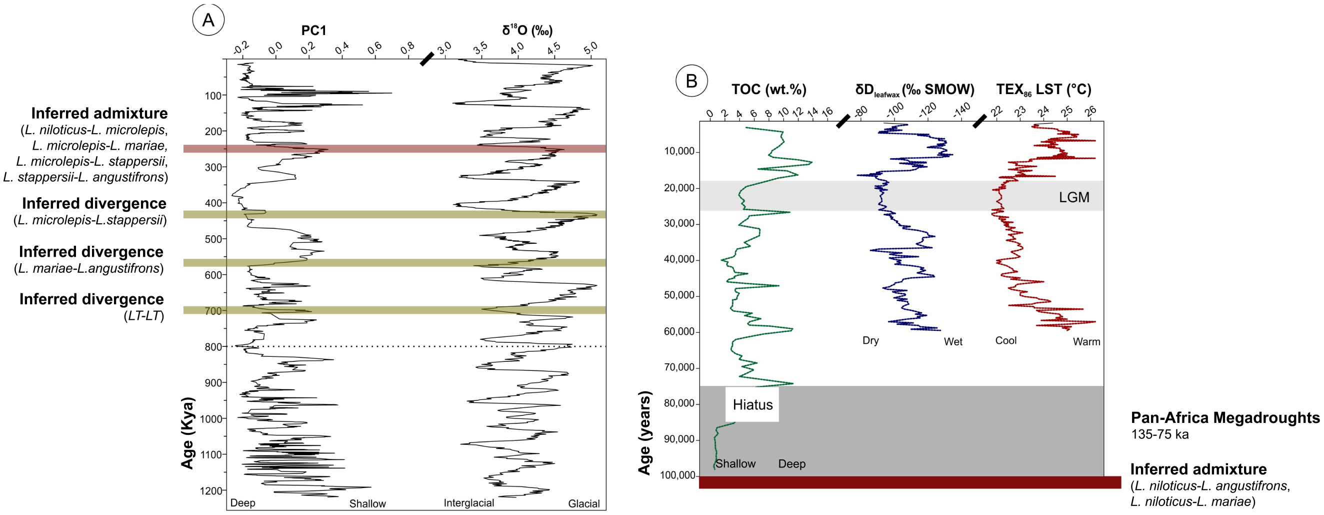


**Fig. S1.** Sedimentary records of Quaternary environmental change for eastern Africa. (A) Lake level reconstruction derived from paleoecological data (pollen, ostracodes, mollusks, charcoal) from central Lake Malawi (Ivory et al., 2016). PC1 is the first principal component of core data containing fossil remnants (biogenic) and mineral bearing (mineralogical) sediments from the MAL05-1B core from Lake Malawi, presented in Ivory et al. (2016), which correlates with deep versus shallow lake conditions. Amplitude and frequency of inferred lake level oscillations shift at approximately 800 Kya (dotted line), near the same time that global benthic oxygen isotope data suggest that glacial-interglacial cycles transitioned from a 41 kyr to a 100 kyr periodicity (e.g., Lisiecki and Raymo, 2005). (B) Climate and lake level proxy reconstructions from deepwater sediments from Lake Tanganyika. Water levels declined several hundered meters during the Last Glacial Maximum (LGM; Tierney et al., 2008) and the Pan-African Megadrought intervals ($\sim$135–75 ka; Scholz et al., 2007); both episodes (grey shading) generated hiatuses observable on seismic profiles, owing to subaerial exposure of the lake floor. These regressions lead to persistent hydrological closure of Lake Tanganyika, disconnecting the lake from the Congo Basin as water level elevation dropped below the elevation at the Lukuga River outlet on the western margin (Fig. 1).

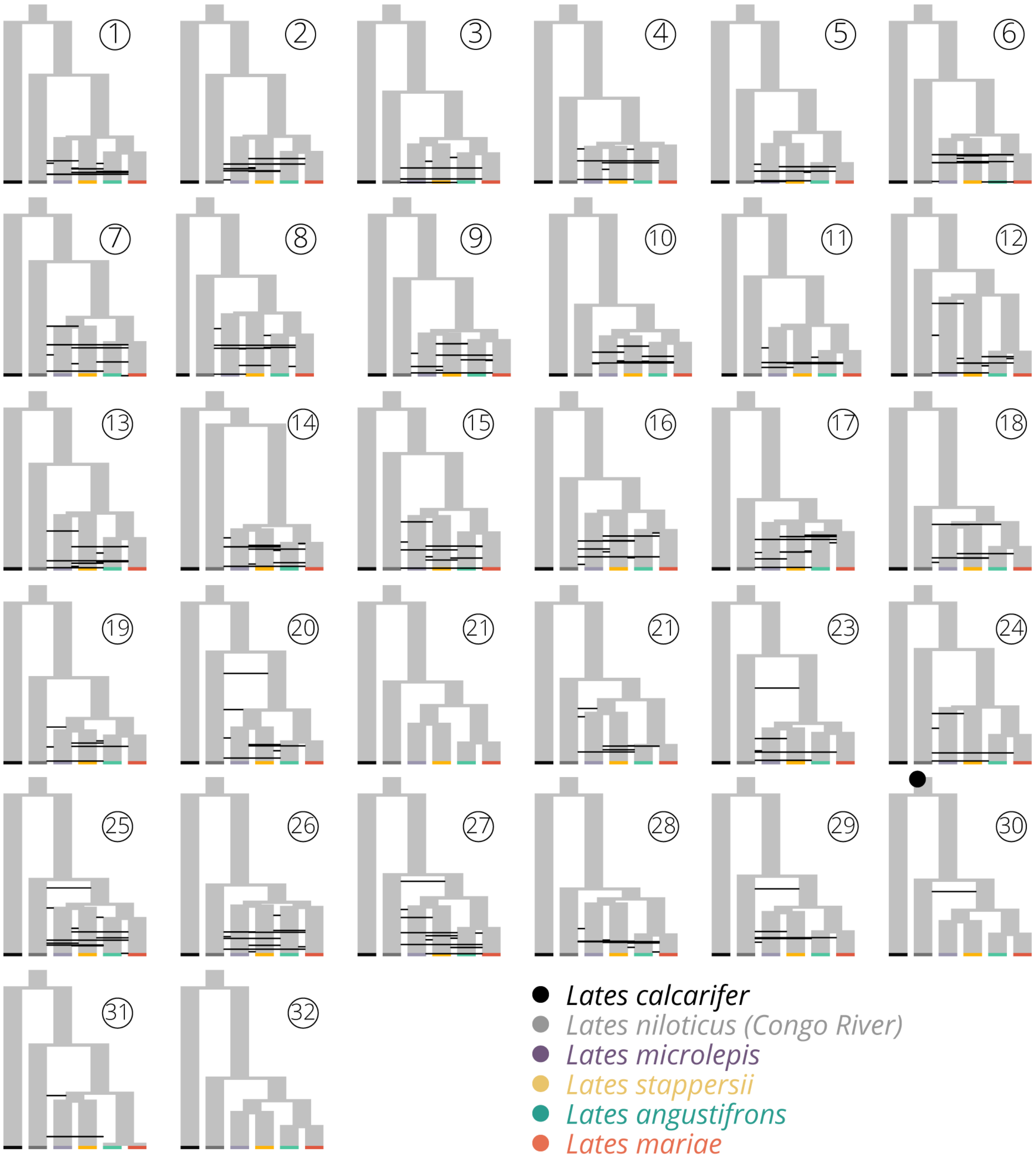


**Fig. S2.** Diagrams of models tested in `fastsimcoal2` analyses. All topologies were constrained to the `RAxML` species tree topology, with the split between *L. calcarifer* and the freshwater *Lates* constrained using the oldest known *Lates* fossil found in Africa (*L. qatraniensis* from the Jebel Qatrani Formation of the Fayum Depression in Egypt, ∼33 MYA; Murray & Attia 2004). Lines indicate admixture events parameterized in each model. All model specification files can be found at http://github.com/jessicarick/lates-fastsimcoal-scripts.

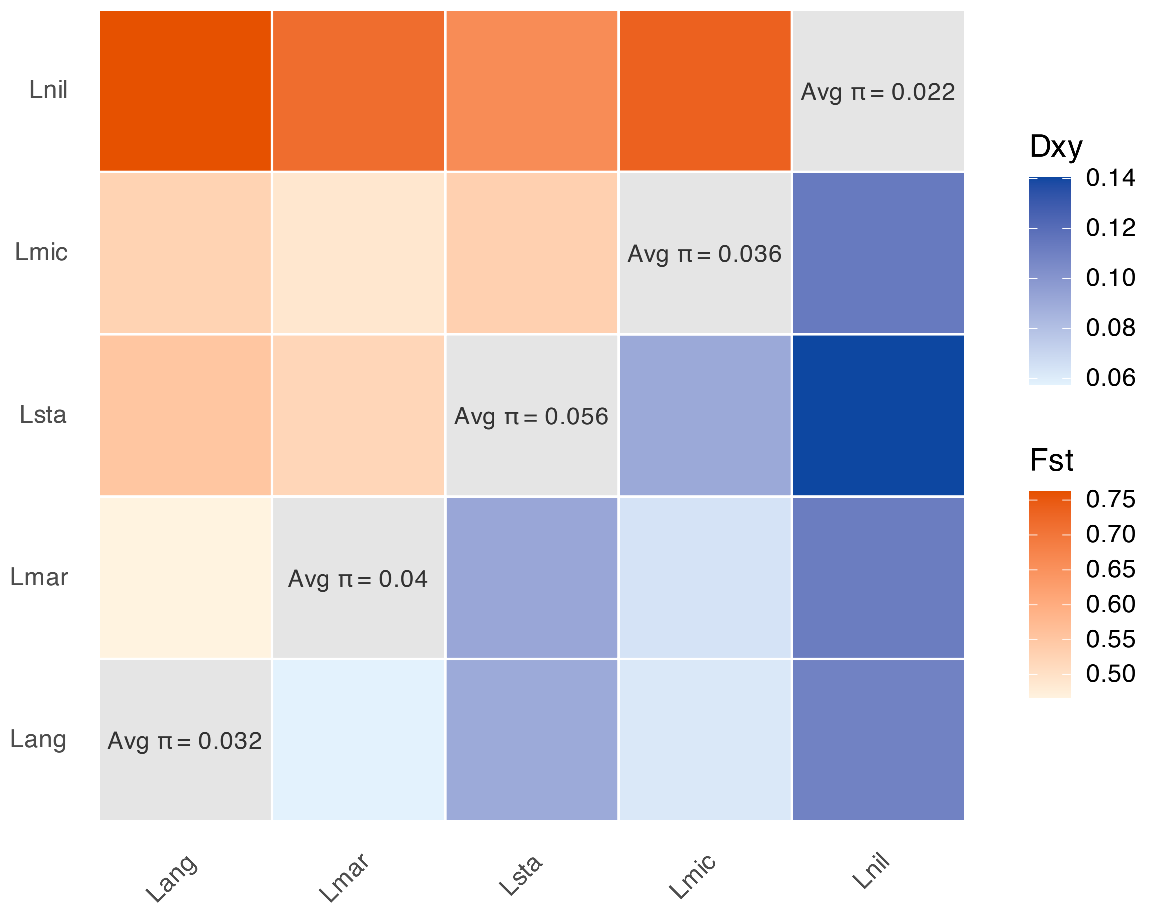


**Fig. S3.** Estimates of genome-wide divergence between species from whole genome resequencing data, including $d_{xy}$ (below diagonal, in blue) and $F_{ST}$ (above diagonal, in red), estimated using `pixy` (**?**). Numbers along the diagonal show mean genome-wide estimates of nucleotide diversity ($\pi$) for each species. The following taxon abbreviations are used in the figure: Lang (*L. angustifrons*), Lmar (*L. mariae*), Lsta (*L. stappersii*), Lmic (*L. microlepis*), Lnil (*L. niloticus*).

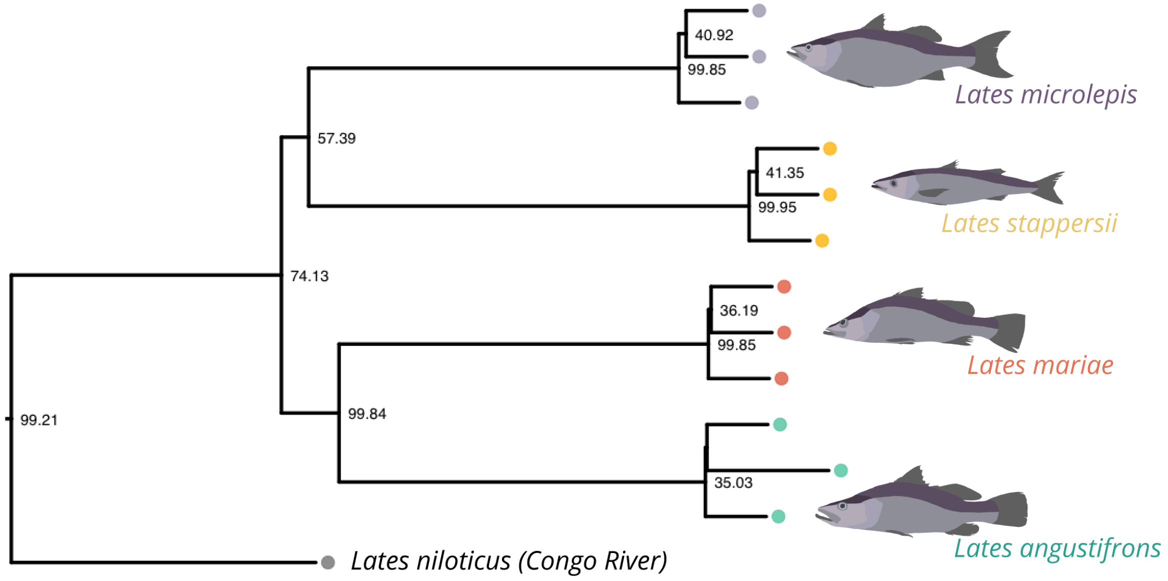


**Fig. S4.** Phylogeny inferred from quartet-based multispecies coalescent species tree reconstruction in `ASTRAL-III` using whole genome data. Node annotations indicate normalized quartet scores (i.e., the percentage of quartets in the 4,000bp gene trees that agree with a split). The tree was rooted using *Lates calcarifer* as an outgroup.

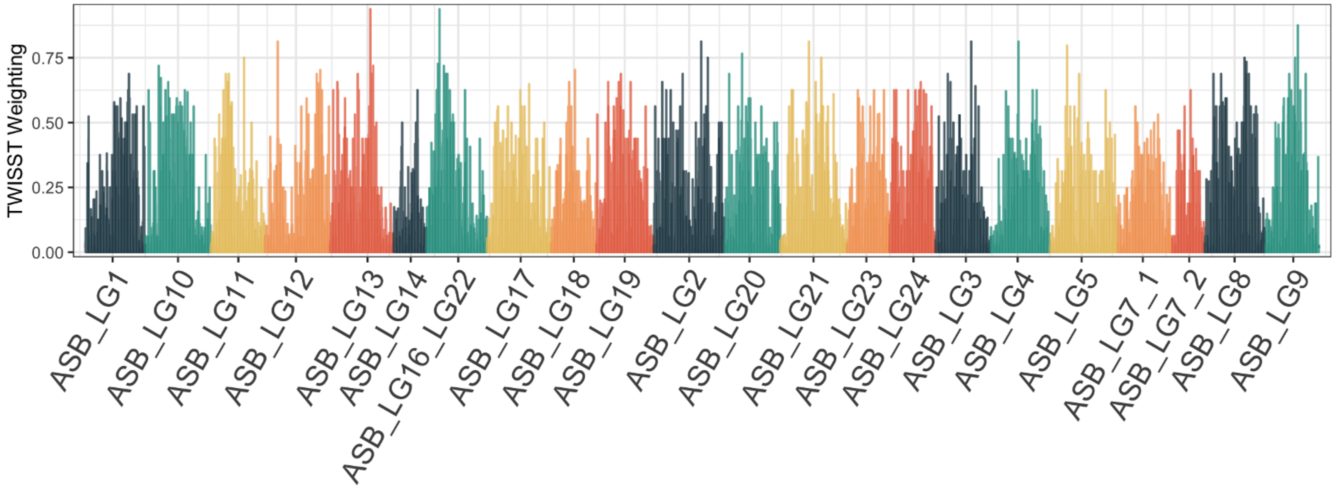


**Fig. S5.** Manhattan plot of topology weightings from `TWISST2`, which have been smoothed in 10kb windows. Weightings shown are the sum for all topologies with the correct sister taxa relationships with Lake Tanganyika and a monophyletic Lake Tanganyika *Lates* clade.

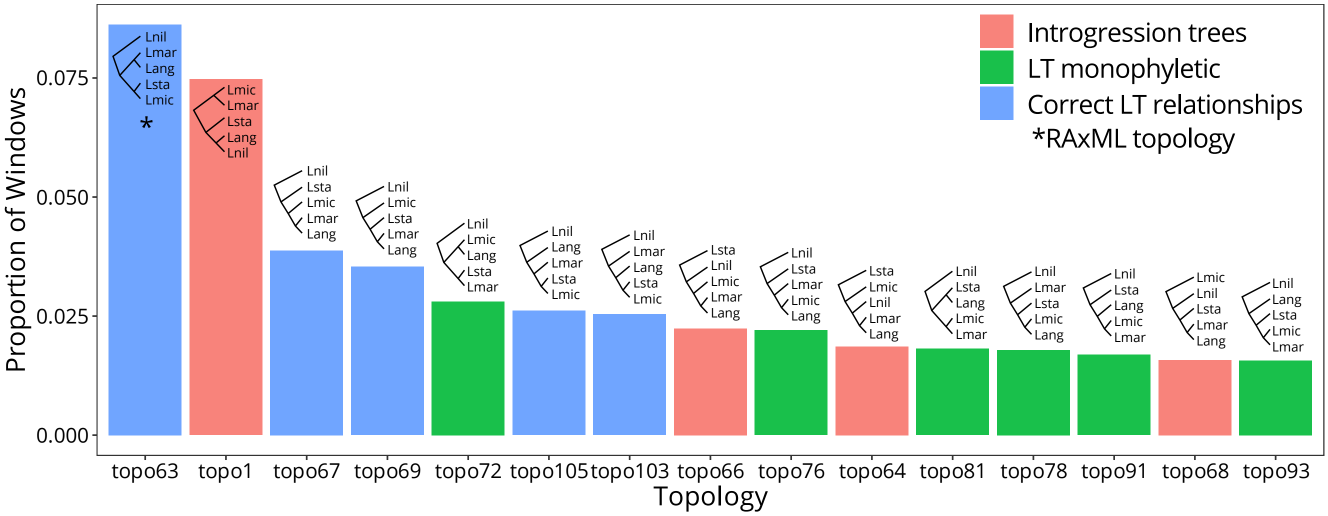


**Fig. S6.** Summary of topology weighting findings from `TWISST2`, with the proportion of windows supporting the top 15 topologies (out of 105 total possibilities) shown, sorted by the proportion of the windows across the genome that support the given topology. Topologies have been categorized by their general features, including whether they have the correct sister relationships within the Lake Tanganyika taxa (blue), whether they have incorrect sister relationships but a monophyletic Lake Tanganyika clade (green), or if they have *L. niloticus* nested within the Lake Tanganyika clade (red). The topology supported by our RAxML findings was the one with the most windows supporting it and is indicated with a star.

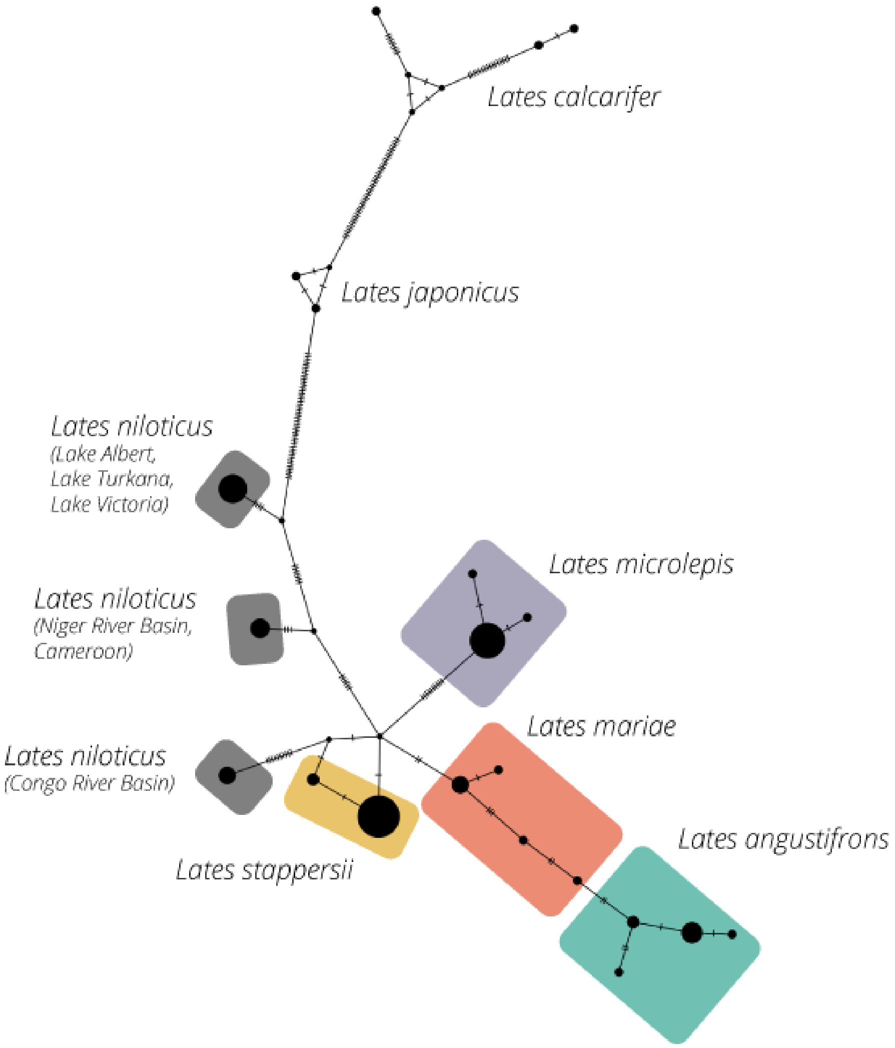


**Fig. S7.** Median-joining haplotype network inferred from n=87 individual *Lates* fishes' 16s mitochondrial sequencing data generated in this paper and combined with sequences from Koblmüller et al. (2021) and Li et al. (2011). Each circle represents a distinct haplotype, with tick marks indicating mutations between haplotypes and circle size representing the number of individuals in the data set with the given haplotype.

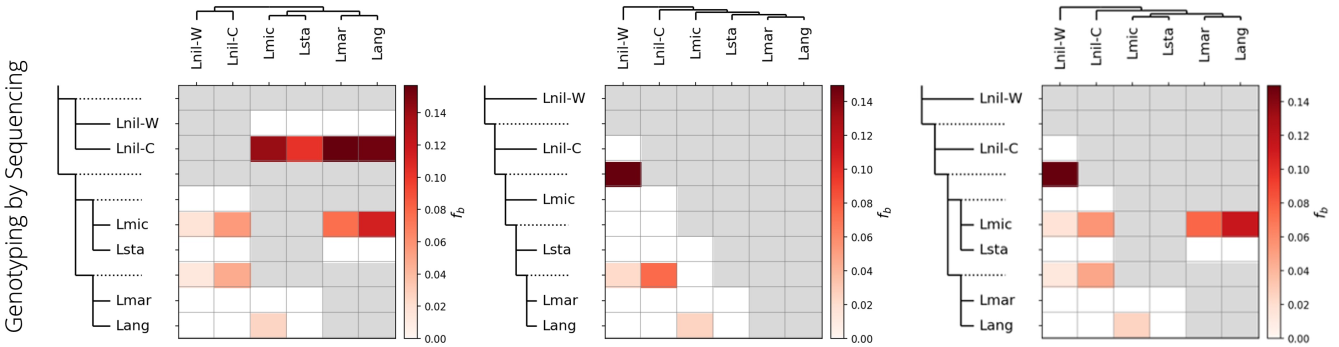


**Fig. S8.** A comparison of Fbranch results for the GBS dataset, constrained to three alternate phylogenetic hypotheses for the relationship among *Lates* species: (a) *L. niloticus* as a monophyletic group, with two sister groups within Lake Tanganyika; (b) a paraphyletic *L. niloticus*, with pectinate toplogy within Lake Tanganyika (sensu Koblmüller et al., 2021); and (c) a paraphyletic *L. niloticus*, with two sister groups within Lake Tanganyika (as seen in our GBS phylogeny, Fig 1). The following taxon abbreviations are used in the figure: Lang (*L. angustifrons*), Lmar (*L. mariae*), Lmic (*L. microlepis*), Lsta (*L. stappersii*), Lnil-W (*L. niloticus* from West Africa), Lnil-C (*L. niloticus* from the Congo River basin).

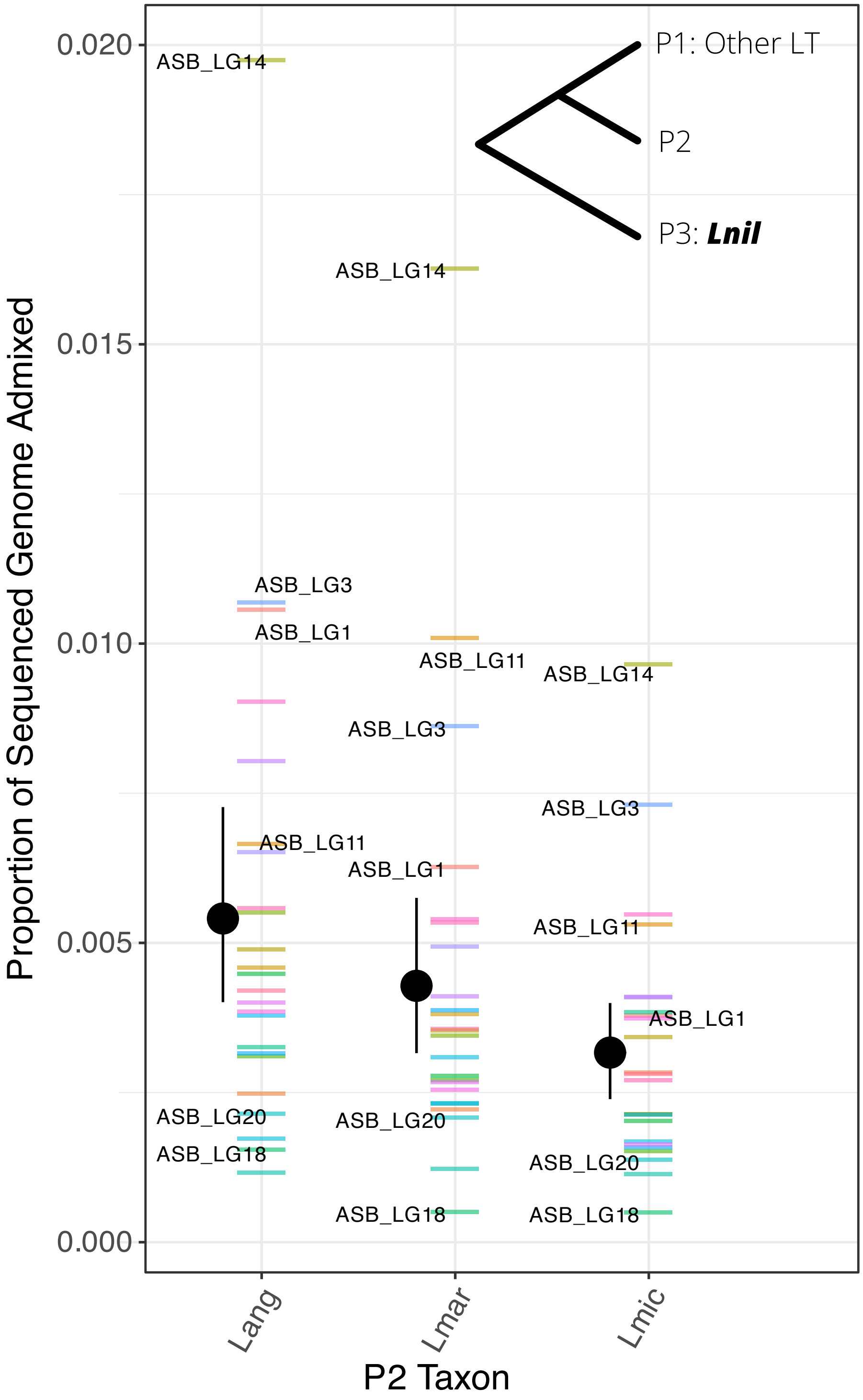


**Fig. S9.** Proportion of genome with an elevated signal of admixture ($z_{fd} > 2.33$, corresponding to $p < 0.01$) for each choice of P2 taxon, averaged across iterations using the other Lake Tanganyika (LT) species as P1 taxa. Lines show the proportion for each chromosome, while black points and bars indicate the mean and 95% confidence interval for the overall genome-wide proportion for the given P2 taxon. To calculate the proportion, windows were combined into blocks if adjacent windows had elevated $f_d$ signals, and the sum total length of significant blocks for a chromosome was divided by the total sequenced length of the chromosome. Estimates of $f_d$ were calculated in 100bp windows across the genome in `Dsuite` (Malinsky et al., 2021), for each trio comprised of two LT *Lates* species and *L. niloticus* that showed an elevated $D$-statistic in genome-wide analyses. Values of $f_d$ for windows with $D < 0$ have been converted to missing data, as recommended. A few of the chromosomes with highest and lowest proportions have been annotated. The following taxon abbreviations are used in the figure: Lang (*L. angustifrons*), Lmar (*L. mariae*), Lmic (*L. microlepis*), Lnil (*L. niloticus*).

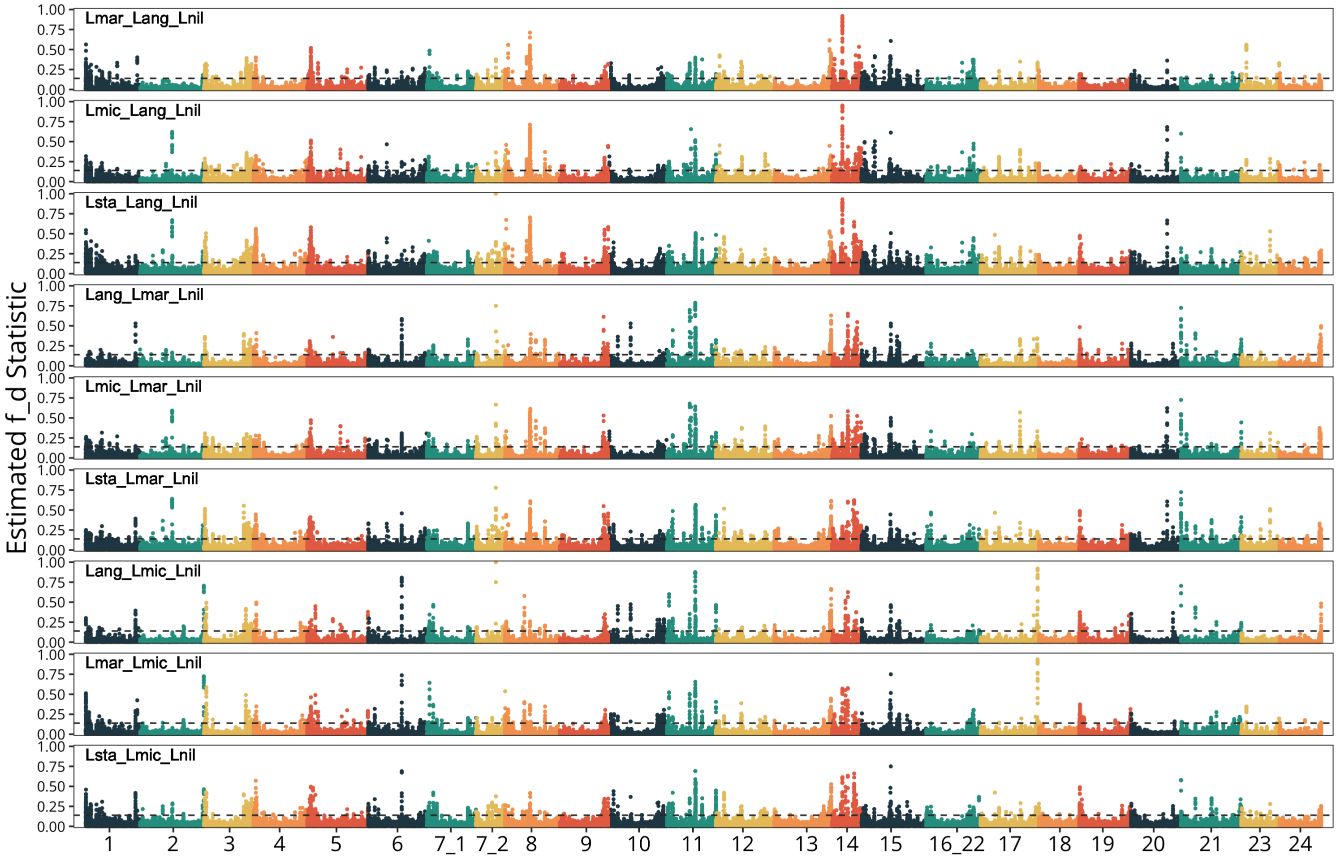


**Fig. S10.** Signals of admixture ($f_d$) in 100bp windows across the genome, as calculated in `Dsuite` (Malinsky et al., 2021), for each trio comprised of two Lake Tanganyika *Lates* species and *L. niloticus*. Dashed lines indicate $p = 0.01$. Values of $f_d$ for windows with $D < 0$ have been corrected to NA, as recommended.

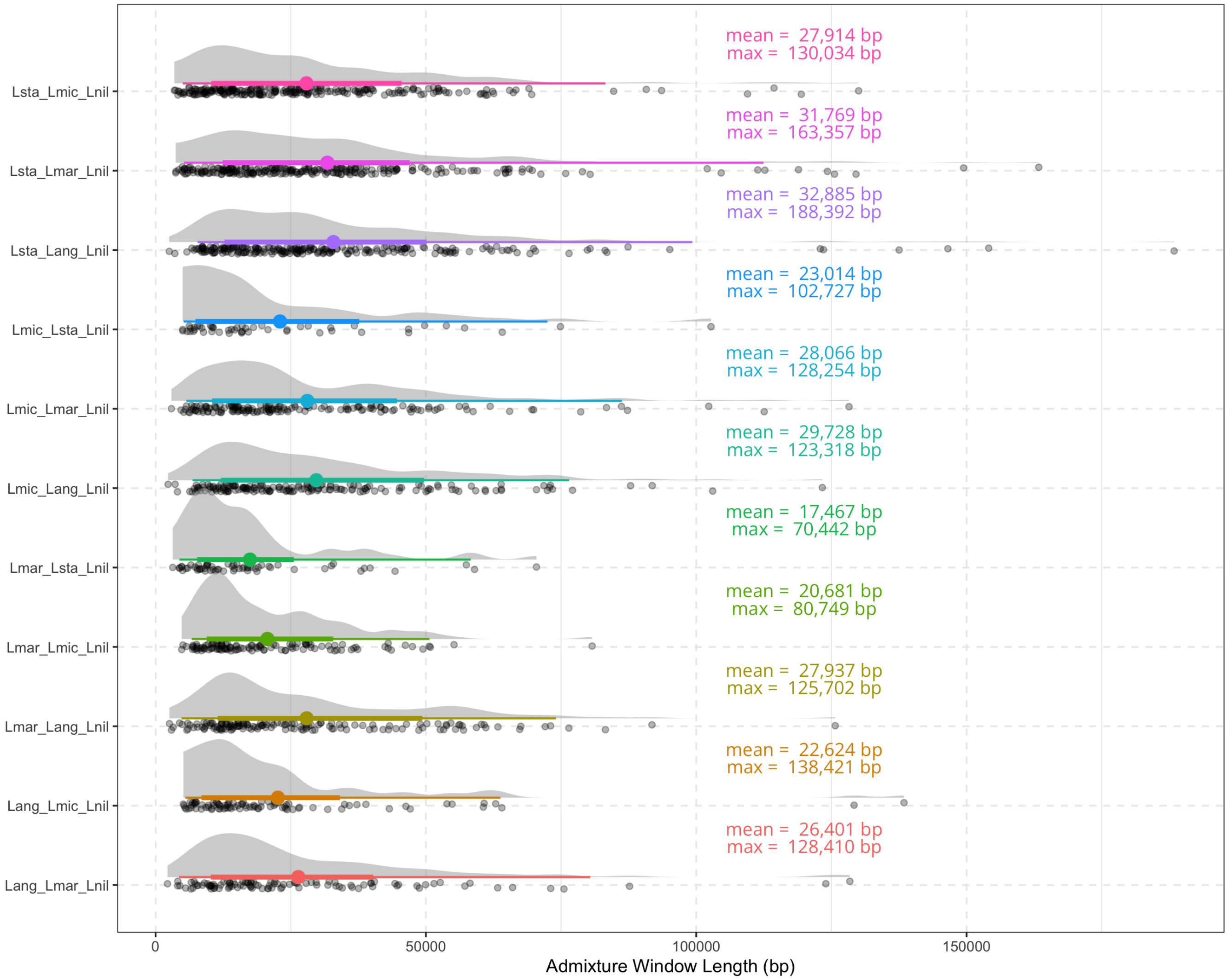


**Fig. S11.** Distribution of block sizes for windows with a significantly elevated signal of admixture for each three-taxon comparisons. Points show the overall median block size, while bars indicate the 2.5% and 97.5% quantiles. To calculate block sizes, windows were seeded using a threshold of $z > 2.33$ and merged if adjacent windows had elevated $f_d$ signals using a more tolerant threshold of $z > 1.5$ and allowing for up to 3 windows within a range to be lower than this threshold. Estimates of $(f_d)$ were calculated in 100 SNP windows with 20 SNP step size across the genome in `Dsuite` (Malinsky et al., 2021), for each trio comprised of two Lake Tanganyika *Lates* species and *L. niloticus*. Values of $f_d$ for windows with $D < 0$ have been converted to missing data, as recommended. The following taxon abbreviations are used in the figure: Lang (*Lates angustifrons*), Lmar (*L. mariae*), Lmic (*L. microlepis*), Lsta (*L. stappersii*), Lnil (*L. niloticus*).

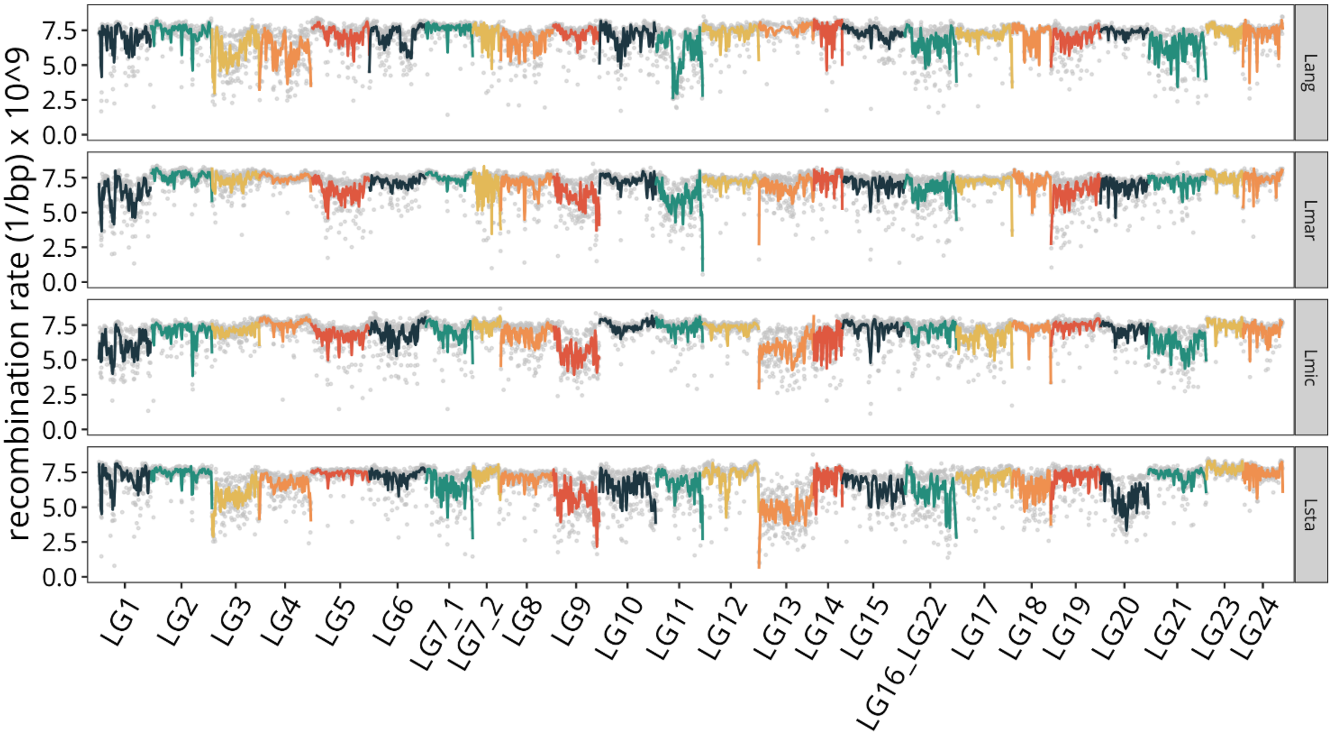


**Fig. S12.** Recombination rates across the genome for each of the four Lake Tanganyika *Lates* species, as estimated using `ReLERNN`. Points show estimates for windows with >100 sites, and lines show smoothing using loess regression.

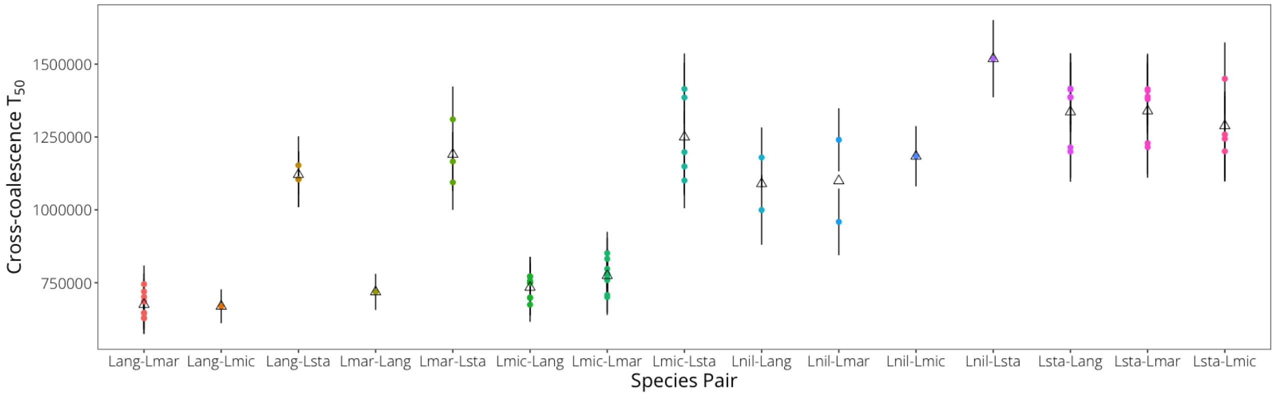


**Fig. S13.** Divergence time estimates from relative cross-coalescence rates inferred using `MSMC2`, with individual sequence pairs categorized by species pair. Divergence times were estimated as the time (in years) at which the relative cross-coalescence rate reached 50%. Each colored point indicates the estimate for a single pair of individuals, and black triangles show the mean for pairs in the given species comparison. The following taxon abbreviations are used in the figure: Lang (*Lates angustifrons*), Lmar (*L. mariae*), Lmic (*L. microlepis*), Lsta (*L. stappersii*), Lnil (*L. niloticus*).

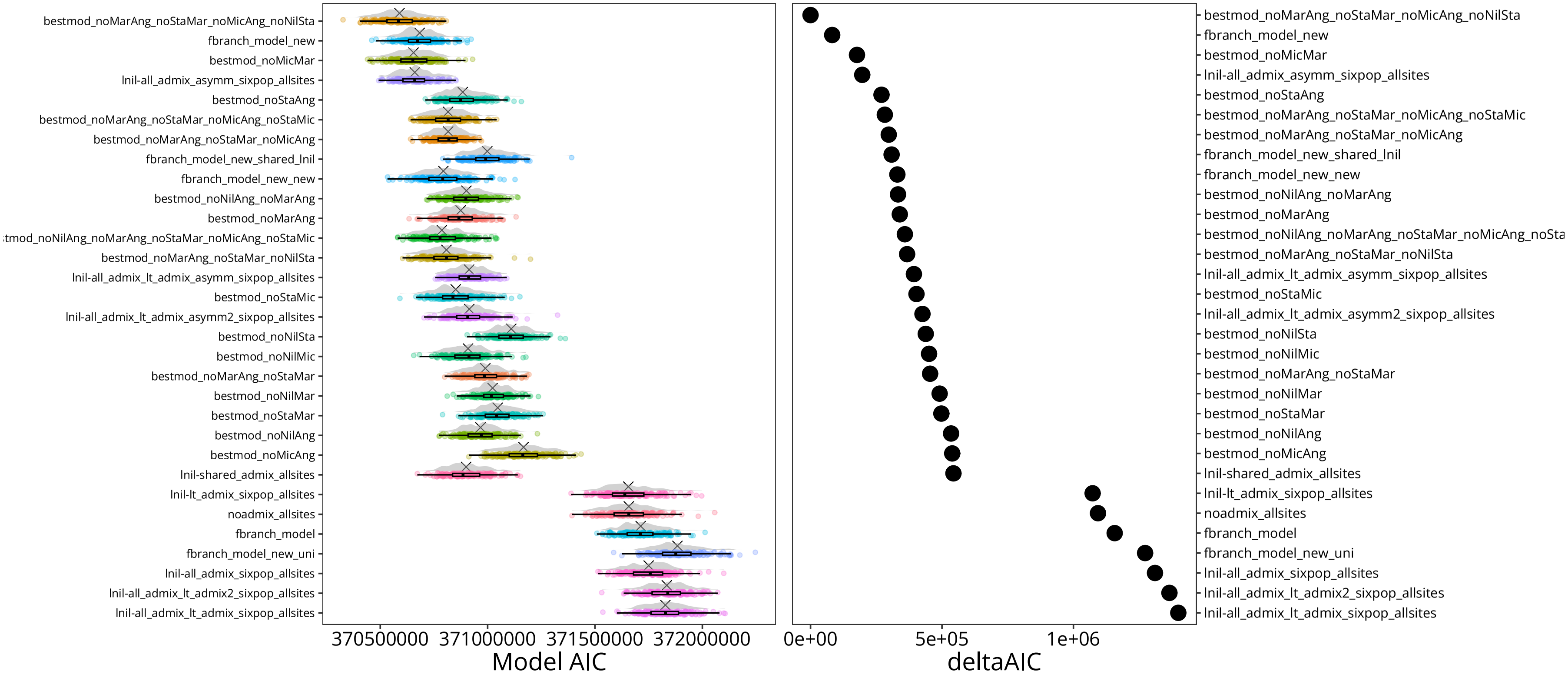


**Fig. S14.** A comparison of model likelihoods (left) and AIC (right) for the six-taxon models tested. Model parameters are detailed in Table S6. Models are ordered based on AIC estimates; the best models were chosen based on lowest AIC and highest likelihood distributions. For AIC calculations, parameters were penalized by 200. The vertical dashed line indicates the lower 95% quantile for the top model's distribution of likelihood values. Note that model numbers match the diagram in Fig. S2).

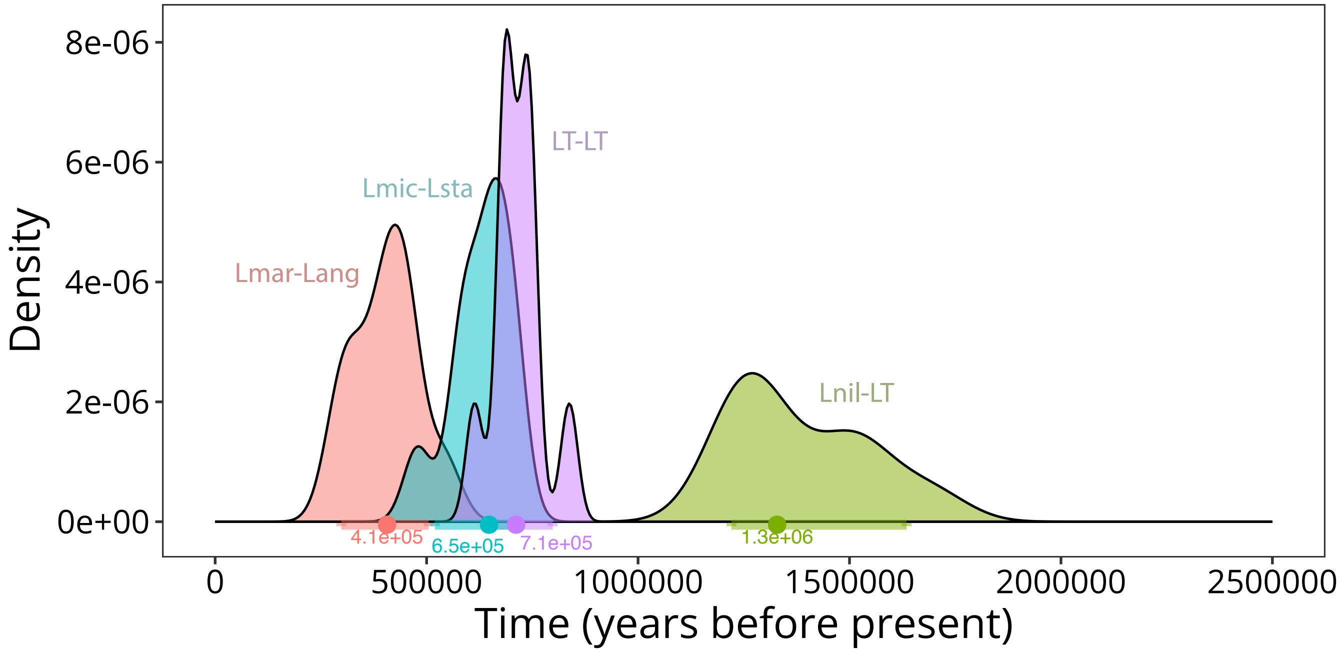


**Fig. S15.** A comparison of divergence time estimates from the top ten best `fastsimcoal2` models, demonstrating that our choice of top model does not significantly bias our divergence time results. The distribution of divergence estimates for the *L. niloticus* (Lnil)–Lake Tanganyika (LT) clade split has a median estimate of 1.3 Mya (range 1.2–1.7 Mya); the split between the two LT sister clades (LT–LT) has a median estimate of 712 Kya (range 613–837 Kya); the median estimate for the *L. mariae* (Lmar)–*L. angustifrons* (Lang) divergence time is 647 Kya (range 478–714 Kya); and the median for the *L. microlepis* (Lmic)–*L. stappersii* (Lsta) split is 407 Kya (range 287–542 Kya).

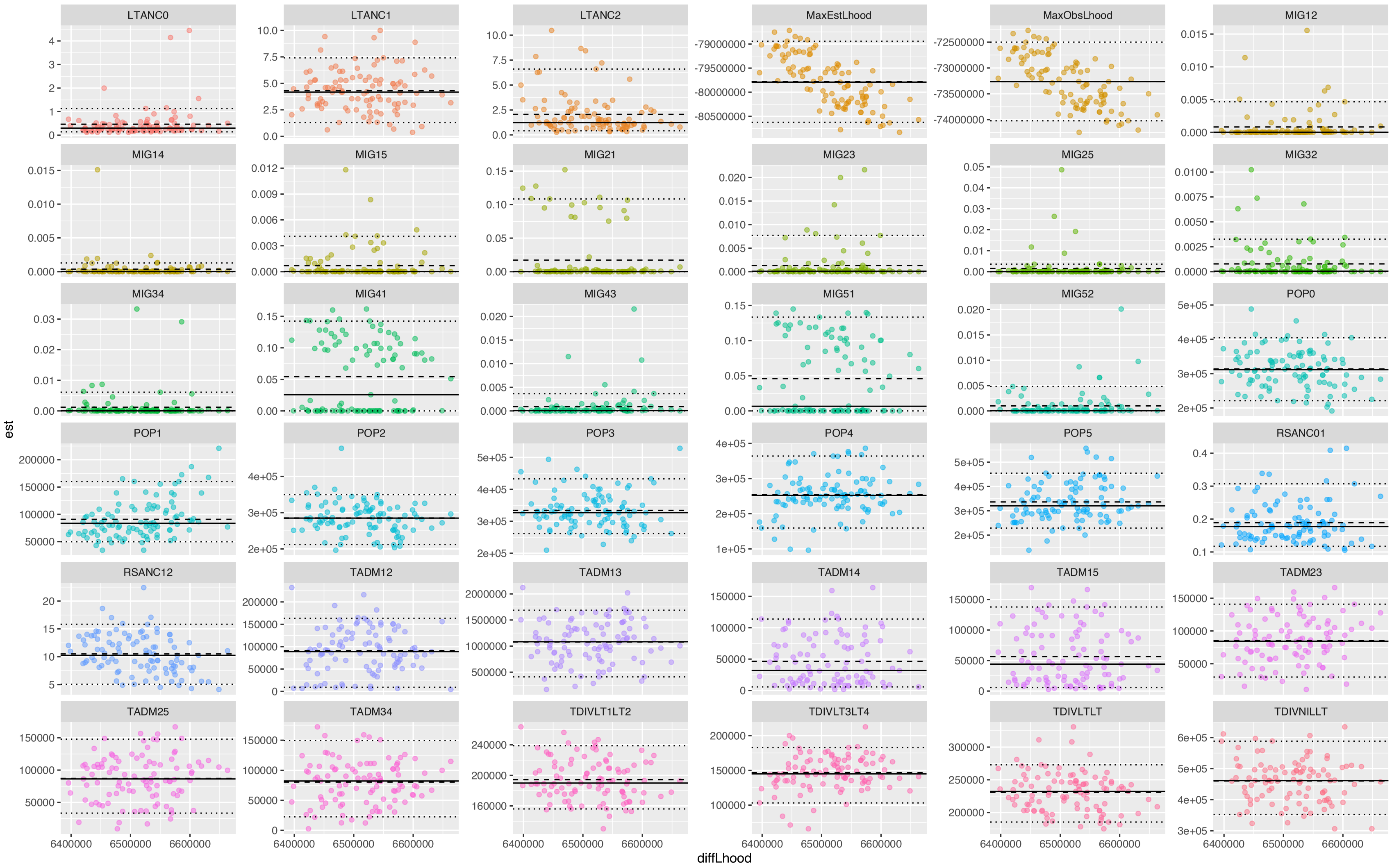


**Fig. S16.** Distribution of parameter estimates from nonparametric bootstrapping of 200 datasets in `fastsimcoal2` for our best model, with each parameter estimate plotted against the difference in observed and expected likelihoods for its containing model. Dashed lines indicate the 95% bootstrap confidence intervals for each parameter. Details of summary statistics for each parameter can be found in Table S6. Solid lines indicate median of the parameter distributions, which are the estimates plotted in Fig. 2.

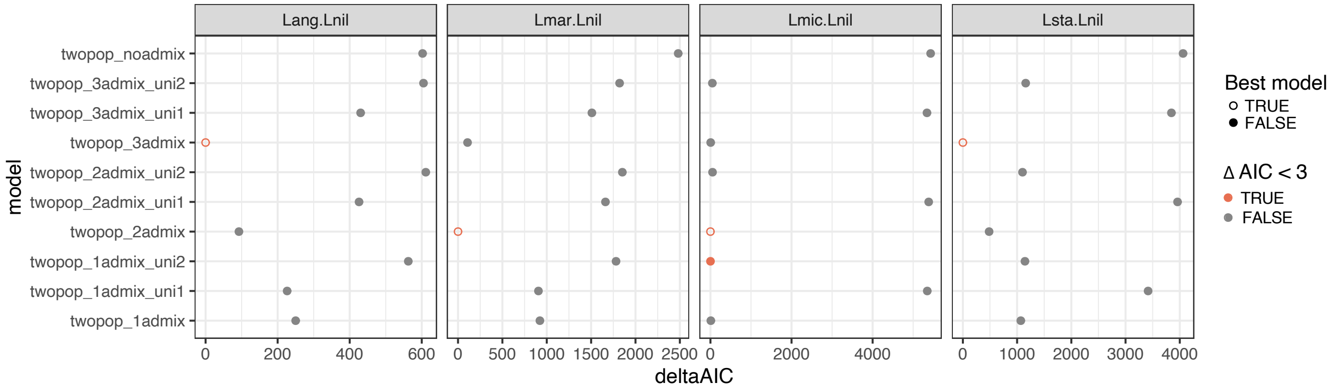


**Fig. S17.** A comparison of AIC values for the two-taxon `fastsimcoal2` models tested for comparisons between *Lates niloticus* (Lnil) and each of the Lake Tanganyika taxa: Lang (*L. angustifrons*), Lmar (*L. mariae*), Lmic (*L. microlepis*), Lsta (*L. stappersii*). The best models were chosen based on lowest AIC and highest likelihood values. For AIC calculations, parameters were penalized by 25. Models in red have an AIC value within 3 points of the top model's AIC. For each species pair, we tested models with no admixture ("noadmix"), a single admixture event ("1admix"), two admixture events ("2admix"), or three admixture events ("3admix"), and within each model involving admixture, we also tested models with gene flow occurring unidirectionally in one direction or the other ("uni1" and "uni2").

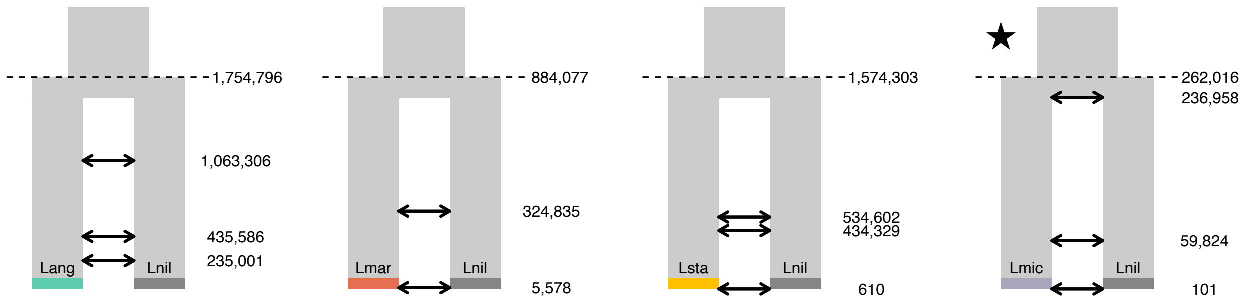


**Fig. S18.** Visual representations of the inferred best models for each two taxon `fastsimcoal2` comparison involving *Lates niloticus* and the Lake Tanganyika *Lates* species, including parameters inferred for the given model. Stars indicate species pairs for which there was at least one alternative model where $\Delta AIC < 3$ (Fig. S17). Divergence times are indicated for the split between the two taxa, as well as inferred timings of admixture events. The following taxon abbreviations are used in the figure: Lang (*Lates angustifrons*), Lnil (*L. niloticus*), Lmar (*L. mariae*), Lsta (*L. stappersii*), Lmic (*L. microlepis*).

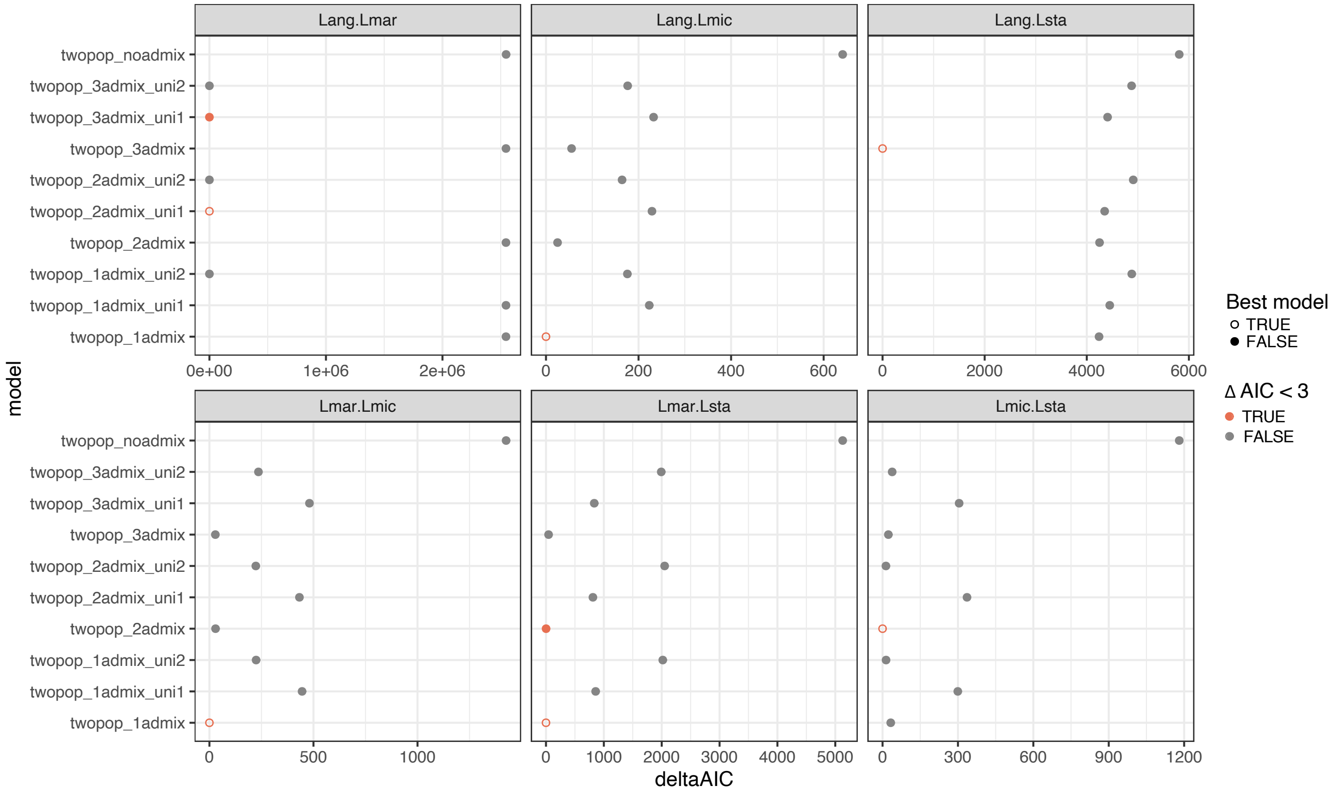


**Fig. S19.** A comparison of model likelihoods AIC values for the two-taxon `fastsimcoal2` models tested for comparisons within Lake Tanganyika taxa [Lang (*L. angustifrons*), Lmar (*L. mariae*), Lmic (*L. microlepis*), Lsta (*L. stappersii*)]. The best models were chosen based on lowest AIC and highest likelihood values. For AIC calculations, parameters were penalized by 25; models with a $\Delta AIC < 3$ are indicated in red. For each species pair, we tested models with no admixture ("noadmix"), a single admixture event ("1admix"), two admixture events ("2admix"), or three admixture events ("3admix"), and within each model involving admixture, we also tested models with gene flow occurring unidirectionally in one direction or the other ("uni1" and "uni2").

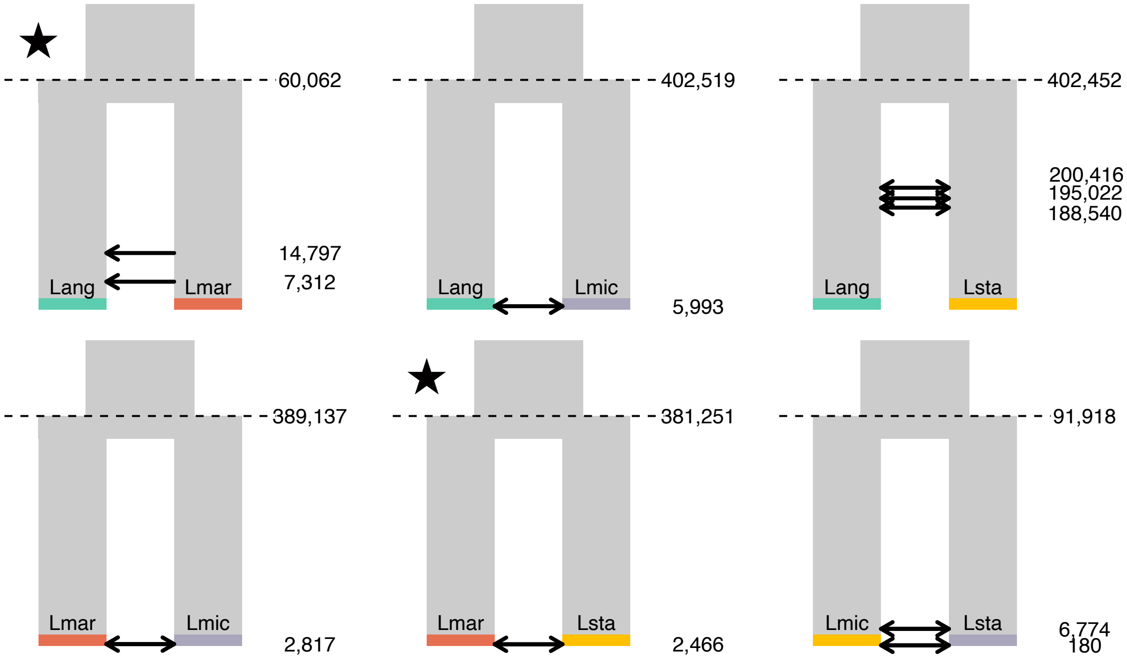


**Fig. S20.** Visual representations of the inferred best models for each two taxon `fastsimcoal2` comparison between LT *Lates* taxa, including parameters inferred for the given model. Stars indicate species pairs for which there was at least one alternative model where $\Delta AIC < 3$ (Fig. S19). Divergence times are indicated for the split between the two taxa, as well as inferred timings of admixture events. The following taxon abbreviations are used in the figure: Lang (*Lates angustifrons*), Lmar (*L. mariae*), Lmic (*L. microlepis*), Lsta (*L. stappersii*).

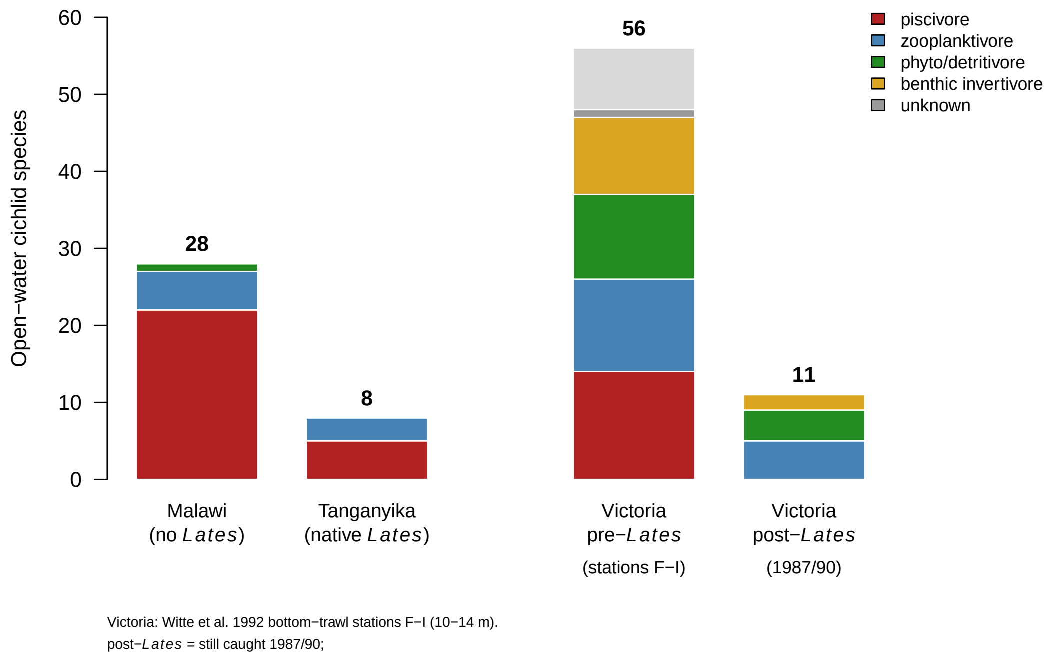


**Fig. S21.** Richness of pelagic cichlid species by feeding guild in lakes with different *Lates* histories: Lake Malawi (no *Lates*), Lake Tanganyika (native *Lates*), and Lake Victoria before and after the introduction of Nile perch (*Lates niloticus*). Numbers above bars give total species. For Malawi and Tanganyika, species included are those coded with "open water" as their complete or partial habitat description in **?**, and guilds are assigned based on classifications in that same that dataset. For Victoria, we included taxa recorded at offshore bottom-trawl stations F–I (10–14 m depth) of Mwanza Gulf in Witte et al. (1992), with trophic categories for these species matching those of McGee et al. (2015). The Victoria post-*Lates* bar shows the subset of those taxa still caught in 1987/90 in any gear or at any station. One taxon of uncertain post-*Lates* status is not counted. There are several caveats to these data and comparisons. First, the data from Malawi and Tanganyika are lake-wide lists whose open-water classification is a coarse habitat code. Victoria is a quantitative survey of one small region with a minimum depth filter. Because *Lates niloticus* in Lake Victoria are broadly demersal, the open-water subset is only part of the cichlid community that interacts with *Lates*. Many demersal Malawi genera are not included the Malawi data, in sum accounting for several hundred species; if we included those species as well the diversity contrast with Tanganyika would be even more extreme.
